# An extra-circadian function for human CLOCK in the neocortex

**DOI:** 10.1101/2023.03.13.531623

**Authors:** Yuxiang Liu, Miles R. Fontenot, Ashwinikumar Kulkarni, Nitin Khandelwal, Seon-Hye E. Park, Connor Douglas, Matthew Harper, Pin Xu, Nisha Gupta, Jay R. Gibson, Joseph S. Takahashi, Genevieve Konopka

## Abstract

Core circadian-related proteins such as the transcription factor CLOCK are ubiquitously expressed and important for regulating molecular pathways underlying circadian rhythms. Previous work has suggested that CLOCK has evolved human neocortex-specific gene regulation and therefore may have extra-circadian functions. To test this in vivo, we generated a mouse model that recapitulates human cortical expression of CLOCK. The CLOCK humanized (HU) mice show enhanced cognitive flexibility, which is associated with the alteration in spatiotemporal expression of CLOCK. Cell type specific genomic profiling of HU mice identified upregulated genes related to dendritic growth and spine formation in excitatory neurons. Consistent with this result, we found that excitatory neurons in HU mice have increased complexity of dendritic branching and spine density, as well as a greater frequency of excitatory postsynaptic currents, suggesting an increase in neural connectivity. In contrast, CLOCK knockout in human induced pluripotent stem cell-induced neurons showed reduced complexity of dendrites and lower density of presynaptic puncta. Together, our data demonstrate that CLOCK evolved extra-circadian gains of function via altered spatiotemporal gene expression and these functions may underlie human brain specializations.

## Introduction

The unparalleled advancement of human cognition can be attributed to the evolution of the human brain. Compared with non-human primates, the human brain has undergone neocortical expansion (Finlay and Darlington, 1995; Striedter, 2005), increased complexity of neuronal dendritic arborization and neural networks (Bianchi et al., 2013; Elston et al., 2001; Sherwood et al., 2020), and alterations in gene expression (Berto et al., 2019; Konopka et al., 2012; Ma et al., 2022; Sousa et al., 2017; Xu et al., 2018; Zhu et al., 2018). However, the underlying molecular mechanisms of these specializations is still not clear.

Comparative genomics among humans and non-human primates has demonstrated that the regulation of gene expression rather than modifications in protein coding sequences is a major facet of molecular evolution driving human brain specializations (Consortium, 2005; King and Wilson, 1975; Konopka and Geschwind, 2010). Thus, elucidating the key regulatory players driving these gene expression programs is critical to understand human brain evolution. Moreover, functional analyses on these candidate genes are required to understand their relative contribution to human brain evolution (Konopka and Geschwind, 2010; Necsulea and Kaessmann, 2014). Few studies have functionally tested the role of specific genes in evolved brain function. Studies of human FOXP2, SRGAP2, and CBLN2 have demonstrated changes in transcriptional programs or neuronal function that are fundamentally altered compared to mouse or non-human primate function (Charrier et al., 2012; Dennis et al., 2012; Enard et al., 2009; Enard et al., 2002; Fossati et al., 2016; Konopka et al., 2009; Schmidt et al., 2021; Shibata et al., 2021).

The transcription factor CLOCK (or circadian locomotor output cycles kaput) has increased human-specific expression in the cerebral cortex (Babbitt et al., 2010; Bakken et al., 2021; Berto and Nowick, 2018; Bozek et al., 2014; Khrameeva et al., 2019; Konopka et al., 2012). Of note, *CLOCK/Clock* is the only core circadian gene that itself lacks rhythmicity in the cortex (Chen et al., 2016; Kobayashi et al., 2015; Li et al., 2017; Mure et al., 2018; Rath et al., 2013; Rath et al., 2014). Transcriptome analyses discovered that a substantial number of downstream targets of mouse CLOCK are also nonrhythmic (Miller et al., 2007; Panda et al., 2002). We further showed that transcriptional targets of human CLOCK in neuronal cultures are also not regulated in a cyclical manner (Fontenot et al., 2017). Moreover, unlike most core circadian genes that have elevated expression following the opening of the eyes to coordinate light entry, *CLOCK/Clock* maintains stable expression across all developmental stages (Kobayashi et al., 2015). Together, these results support an extra-circadian function for CLOCK in the neocortex.

CLOCK has also been implicated in human brain function via its association with human cognitive disorders. We showed that *CLOCK* is a hub gene in a human-specific neocortical co-expression network that has significant enrichment for genes in involved in neuropsychiatric disorders such as seasonal affective disorder, depression, schizophrenia, and autism (Konopka et al., 2012). Single nucleotide polymorphism analyses have also linked *CLOCK* with a broad spectrum of neuropsychiatric disorders (e.g., autism spectrum disorder, bipolar disorder, schizophrenia, attention-deficit/hyperactivity disorder, major depressive disorder, and addiction) (Lamont et al., 2007; Schuch et al., 2018). Patients with epilepsy have decreased expression of *CLOCK* in the temporal cortex (Li et al., 2017). We showed that *CLOCK* knockdown in human neurons leads to altered expression of genes important for neurodevelopment (Fontenot et al., 2017). However, the mechanisms downstream of CLOCK that are important for human brain function are unknown.

Here, we generated a humanized mouse model that contains both the protein-coding and regulatory regions of human *CLOCK* in a mouse *Clock* knockout background. We then leveraged this mouse model to investigate how human CLOCK affects cognitive behaviors, cell type-specific gene expression, neuroanatomy, and electrophysiology. We discovered that human CLOCK has extra-circadian functions that alter gene expression in a spatiotemporal manner to promote connectivity of neurons for enhanced cognitive flexibility.

## Results

### Generation of CLOCK humanized mice

We generated humanized CLOCK mice (HU) by expressing human *CLOCK* together with its flanking non-coding genomic regions in mice lacking endogenous *Clock* (knockout, KO). A bacterial artificial chromosome (BAC) containing the full-length human *CLOCK* gene was edited through recombineering to include flanking non-coding regulatory regions (**Fig. 1A**) and delivered via pronuclear injection into embryos. Selected positive lines (*CLOCK^hum^;Clock^+/+^*) (see supplemental material and **Figs. S1A-B** for mouse line selection), which contained 7-8 BAC copies on average (**Fig. 1B**), were crossed with *Clock* knockout mice (KO:*Clock^-/-^*) (DeBruyne et al., 2006). F1 heterozygotes (*Clock^+/-^* and *CLOCK^hum^;Clock^+/-^*) were crossed to obtain littermates of humanized (HU:*CLOCK^hum^;Clock^-/-^*), KO, and wildtype (WT:*Clock^+/+^*) mice (**Fig. S1C**). A mouse *Clock*BAC transgenic line (*Clock^mus^;Clock^+/+^*) that we previously generated was similarly crossed to *Clock* knockout mice to obtain mouse *Clock* overexpression mice (MS:*Clock^mus^;Clock^-/-^*) in the same way as HU mice (Antoch et al., 1997).

**Figure 1.**
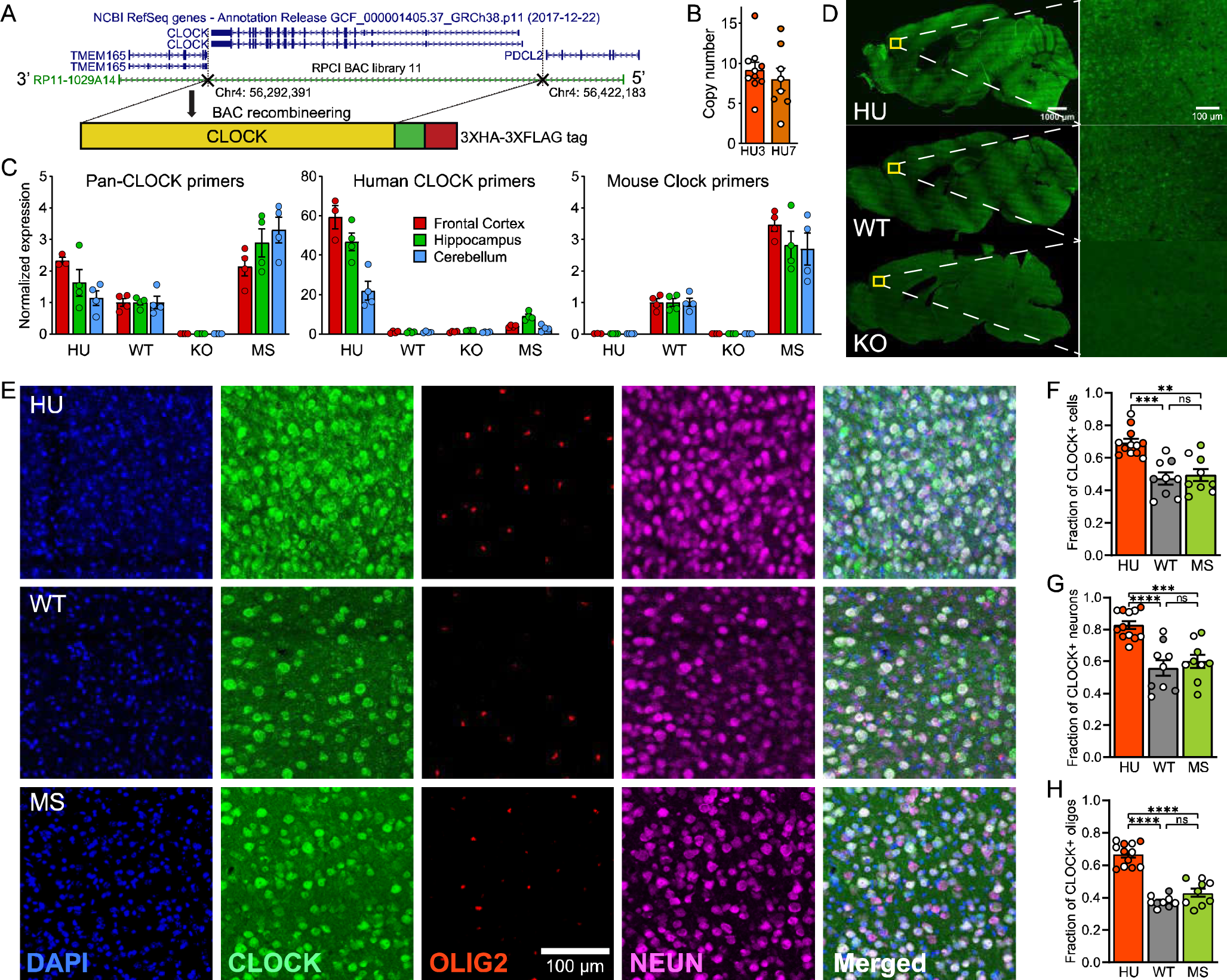
Human BAC transgenic mice express human CLOCK in a greater percentage of cells. **(A)** Recombineering of a BAC to obtain human CLOCK with flanking regulatory regions. **(B)** qRT-PCR results to calculate the number of BAC copies inserted into the genomes of two of the mouse lines. **(C)** qRT-PCR with species-specific *CLOCK/Clock* primers. **(D)** IHC staining for CLOCK in sagittal sections of brain. **(E)** Representative images of IHC staining for CLOCK (green), oligodendrocytes (OLIG2; red), and neurons (NEUN; magenta) in frontal cortex. DAPI indicates nuclei. **(F-H)** Quantification of CLOCK+ fractions in **(F)** all cells, **(G)** neurons, and **(H)** oligodendrocytes. Each data point is an image of subarea in the frontal cortex. We sampled 3-4 mice per genotype and 3 images per mouse. We applied a general linear mixed model (GLMM) with genotype and sex as fixed factors, image as random factor nested with individual, and Tukey’s test for post-hoc analysis. See also **Figure S1**.

Using qRT-PCR, we found that HU and MS mice exclusively expressed human *CLOCK* or mouse *Clock* with about 2.5-fold upregulation in the frontal cortex respectively compared to the WT mice (**Fig. 1C**). This 2.5-fold increase in *CLOCK* is similar to the relative increase we previously reported when comparing human and chimpanzee frontal cortex tissue (Konopka et al., 2012). Using immunohistochemistry (IHC), we observed protein expression of CLOCK in the frontal cortex of both HU and WT mice but not in KO mice (**Fig. 1D**). We then quantified the fraction of CLOCK positive cells in the frontal cortex and found a greater fraction in HU mice than in WT mice (**Fig. 1E,F**). We used anti-NeuN and anti-OLIG2 to label neurons and oligos (including both oligodendrocytes and oligodendrocyte precursor cells) respectively, and found that HU mice demonstrated expression of CLOCK in a greater fraction of both cell types than WT mice (**Fig. 1E,G,H**). The percentage of CLOCK+ neurons in the HU mice (89%) is similar to what has been observed in the human cortex (86%), while the percentage of CLOCK+ neurons in WT mice (62%) is similar to previous reports in mice (66%) (Li et al., 2017). To rule out the possibility that the increased fraction of human CLOCK positive neurons is due to BAC transgenic overexpression, we also carried out IHC in MS mice and found that the fractions of mouse CLOCK+ cells were similar to WT mice and significantly less than the percentages in HU mice (**Fig. 1E-H**). Therefore, human and mouse have different expression patterns of CLOCK/Clock in cortical neurons, and we generated a humanized mouse model that mimics the expression pattern of human CLOCK.

To test whether HU mice express functional CLOCK protein, we tested whether these mice sufficiently rescue *Clock* loss of function behaviors. We test mice in a classic wheel-running paradigm for circadian rhythms and observed a shortened locomotor activity period in both constant light and darkness in KO mice compared to the WT mice, which replicated previous results (Dallmann et al., 2011; DeBruyne et al., 2006). HU mice could partially rescue the reduction of period in both constant light and darkness conditions (**Fig. S1D-F**), which is consistent with the rescue of period in *ClockΔ19* mice with mouse *Clock* overexpression (Antoch et al., 1997). Like *ClockΔ19* mice, adult KO mice are heavier than WT mice, whereas HU mice partially rescue the increase in body weight normalized to body length (**Fig. S1G,H**) (Turek et al., 2005). Because *ClockΔ19* mice had previously been shown to demonstrate depression-like behavioral phenotypes (Easton et al., 2003; Roybal et al., 2007), we carried out a tail suspension test. KO mice showed decreased immobility, in line with previous results, and this was partially rescued in HU mice (**Fig. S1I**). Finally, we ran a resident-intruder test to quantify aggressive behavior and found that KO mice were more aggressive than WT mice while HU mice rescued the aggressive behavior completely (**Fig. S1J**). Performance of KO mice was consistent with the behavior of *Nr1d1* knockout mice (Chung et al., 2014). NR1D1 is a downstream target of CLOCK (Preitner et al., 2002; Takahashi, 2016) and our result suggests that human CLOCK can effectively regulate downstream targets in the mouse genome. In summary, we generated a humanized mouse model that expresses functional human CLOCK and mimics the expression pattern of CLOCK in human cortical neurons and therefore can be used as a valid model to study brain function of human CLOCK.

### Human CLOCK enhances cognitive flexibility

Because CLOCK has been linked with cognitive diseases, we next examined whether HU mice have altered performance in cognitive tests. We tested mice in a series of sensory-motor tests: an open field test that measures both activity and anxiety, an olfactory discrimination test, an eye blink reflex, an ear twist reflex, and a whisker twist orientation. All of these were carried out to determine if manipulation of *CLOCK/Clock* could alter sensory and motor functions of cognition. There were no differences across all genotypes in all of these tests (**Fig. S2A-C**). In the novel object recognition test, recognition memory was marginally impaired in KO mice, while there was no difference between HU and WT mice (**Fig. 2A**). In the fear conditioning test, we did not detect differences in associative learning of auditory cues across genotypes. KO mice showed a deficit in associative learning of context, but HU and WT mice did not differ in any type of associative learning (**Fig. 2B**). We conducted a 5-trial social memory test and found that KO mice were impaired in social learning, while the difference between HU and WT mice was not significant (**Fig. 2C**). These results suggest that human and mouse CLOCK play important and similar roles in general learning and memory. To further differentiate the performance between the HU and WT mice, we challenged them in a set-shifting assay that measures executive function, working memory, and cognitive flexibility. We found that the KO mice were significantly impaired in all tasks of the set-shifting assay, while HU mice required significantly fewer trials to reach criterion than WT mice during the reversal task (**Fig. 2D**). To determine whether the improved performance of HU mice was because of the increase in human CLOCK or random insertion effects of this particular mouse line (human CLOCK line 7, hc7), we repeated the discrimination and reversal tasks in a second humanized line (hc3). The results of hc3 replicated what we found in hc7 (**Fig. 2E**), and we did not find a significant difference between the two lines (**Fig. S2D**). To address whether overexpression of CLOCK in general drove the phenotype or whether human-specific CLOCK regulation and function are driving the behavioral results, we again repeated the discrimination and reversal tasks in MS mice. MS mice showed similar performances as WT mice (**Fig. 2F**) and required more trials to reach criterion than HU mice in the reversal task (**Fig. S2E**). In summary, our results indicate that loss of CLOCK in mice results in broad cognitive deficits, while overexpression of human CLOCK but not mouse CLOCK improves cognitive flexibility independent of motor activity, anxiety or sensorimotor ability.

**Figure 2.**
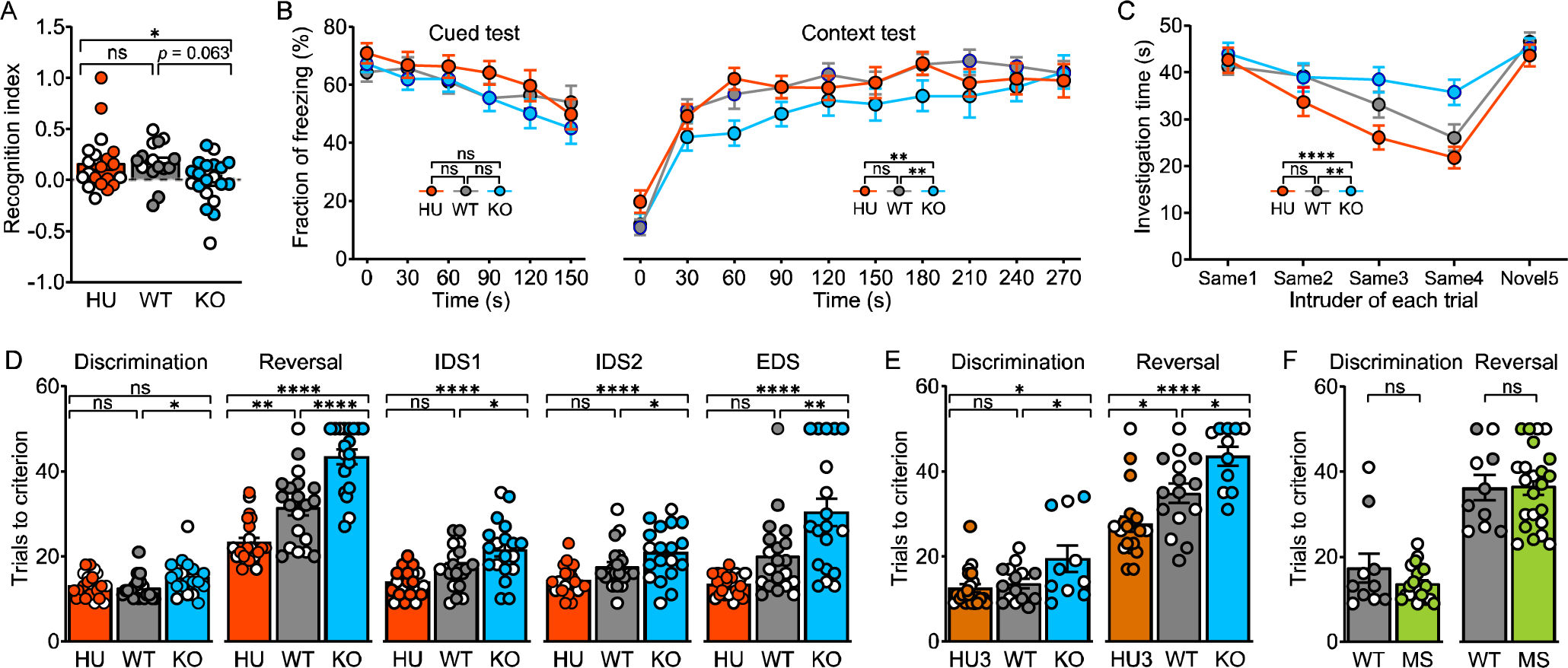
CLOCK affects performance in cognitive tests. **(A)** KO mice show a moderate deficit in a novel object recognition test. **(B)** KO mice have impaired memory of context but not the auditory cue in a fear conditioning test. **(C)** KO mice show social memory deficits in a 5-tńal social memory test. The same intruder was used for the first 4 trials, and then a novel intruder was used for the 5th trial. **(D)** Trials to criterion in 5 continuous tests of a set-shifting assay. IDS: intra-dimensional shift, EDS: extra-dimensional shift. **(E-F)** Performance of reversal learning in **(E)** an alternative humanized mouse line (HU3) and **(F)** MS mice. Each data point is an individual. We applied a general linear model (GLM) with genotype and sex as fixed factors and with Tukey’s test for post-hoc analysis for panel **A, D, E,** and **F.** We did a repeated measures ANOVA with genotype and sex as between-subject factors plus time and trial as within-subject factors for panel B and C respectively. See also **Figure S2** and **Table S2**.

### Human CLOCK increases neuron density without changing cortical lamination

CLOCK is expressed during cortical development (Kang et al., 2011; Loo et al., 2019). We therefore hypothesized that manipulation of CLOCK might result in neurodevelopmental alterations that underlie the observed changes in cognitive behavior. We measured brain mass with and without normalization to body mass, but we did not find any significant differences across genotypes at postnatal day 7 (P07) or P21 (**Fig. S3A-D**). However, although brain mass is unaltered, it was not clear whether CLOCK could alter the organization and structure of the cortex.

Our previous work showed that knockdown of *CLOCK* expression in human neural progenitor derived neurons led to altered migration in an *in vitro* neurosphere assay (Fontenot et al., 2017). Thus, we investigated potential outcomes of migration defects by measuring cortical thickness and lamination of somatosensory cortex in our mouse lines. We used anti-CUX1, anti-FOXP1, and anti-FOXP2 antibodies to label layers 2-4, layers 3-5, and layer 6, respectively (**Fig. S3E**). We found that the absolute cortical thickness was not significantly different across genotypes (**Fig. S3F**), while normalized cortical thickness is decreased in KO mice but not different between HU and WT mice (**Fig. S3G**). Layer thickness was normalized to cortical thickness to quantify lamination. The absolute and normalized thickness of layers 2-4 and layer 6 did not show differences across genotypes (**Fig. S3H-K**). Thus, CLOCK does not alter the laminar organization of the cortex. These results are consistent with other findings in *Emx-Cre Clock* conditional KO mice (Li et al., 2017).

CLOCK is enriched in progenitor-like cells (i.e., neuroepithelial, radial glial, and intermediate progenitor cells) during brain development (Kang et al., 2011; Loo et al., 2019). These progenitor cells ultimately give rise to multiple types of cortical cells. Therefore, we tested whether overexpression of human CLOCK would alter the number of cortical cells. Since we did not find alterations of overall brain mass or cortical thickness in the somatosensory cortex of HU mice, we examined the density of cortical neurons and oligodendrocytes in orbital prefrontal cortex, which is involved in cognitive flexibility of reversal learning (Bissonette and Powell, 2012; Hamilton and Brigman, 2015). We found a greater density of both neurons (anti-NEUN+) and oligodendrocytes (anti-OLIG2+ which marks both mature oligodendrocytes and oligodendrocyte precursor cells) in HU compared to WT mice (**Fig. 3**). To further confirm that this difference is due to the overexpression of human CLOCK specifically, we carried out the same quantification in MS mice and found similar densities as WT mice and significantly lower densities than HU mice (**Fig. 3**). Although the cell density did not differ between WT and KO mice, the decreased cortical thickness of KO mice results in fewer total cells. In summary, CLOCK increases the number of cells in mouse frontal cortex without altering lamination.

**Figure 3.**
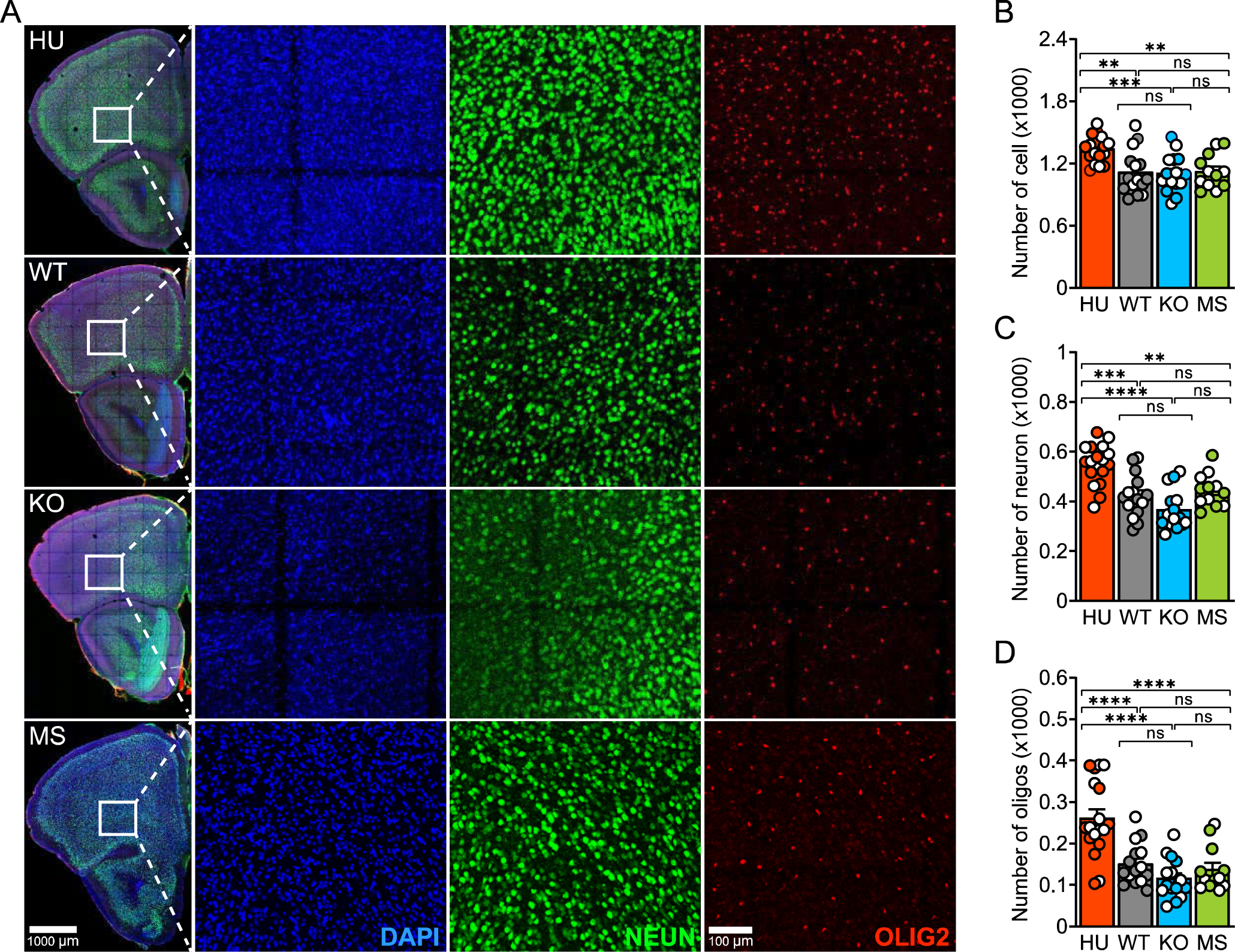
Human CLOCK increases cell density in adult orbital prefrontal cortex. **(A)** Representative image of IHC staining for all cells (DAPI, blue), neurons (NEUN, green), and oligodendrocytes (OLIG2, red) in in orbital prefrontal cortex. **(B-D)** Quantification of cell number in **(B)** all cells, **(C)** neurons, and **(D)** oligodendrocytes. Each data point is a section containing orbital prefrontal cortex. We sampled 4-6 mice per genotype and 3 sections per mouse. We did GLMM with genotype and sex as fixed factors, section as random factor nested with individual, and Tukey’s test for post-hoc analysis. See also **Figure S3**.

### snRNA-seq of frontal cortex reveals differential spatiotemporal expression of human and mouse CLOCK

CLOCK is a transcription factor that is expressed in most cortical cell types (Bakken et al., 2021; Kobayashi et al., 2015; Li et al., 2017). To understand the potential cell type-specific gene regulatory mechanisms underlying the observed alterations of cognitive flexibility, we performed snRNA-seq on the frontal cortex of P07 and P56 mice (n = 2 male and 2 female/genotype; 24 samples in total). Due to high levels of ambient RNA contamination (Caglayan et al., 2022), one sample of each genotype was removed at P07. All samples from both ages had similar quality control metrics including the number of nuclei, unique molecular identifiers (UMIs), and expressed genes, and percentage of mitochondrial transcripts (**Fig. S4A-H**). We integrated the nuclei from all genotypes of the same age and identified major cell types at P07 and P56, respectively (**Fig. 4A,B**). The genotype composition of each cluster is similar and consistent at both P07 and P56 (**Fig. S4I,J**), suggesting that CLOCK does not alter the relative proportion of cell types, although we observed a greater overall number of cells in the frontal cortex (**Fig. 3**). We identified all major cortical cell types (i.e., excitatory neurons, inhibitory neurons, astrocytes, mature oligodendrocytes (MOL), oligodendrocyte precursor cells (OPC), microglia, and endothelial cells) in both P07 and P56 samples (**Fig. S5**; **Table S4**). The number of UMIs and expressed genes are consistent with a previous study across cell types in adult mouse cortex (Hrvatin et al., 2018) (**Fig. S4K-N**).

**Figure 4.**
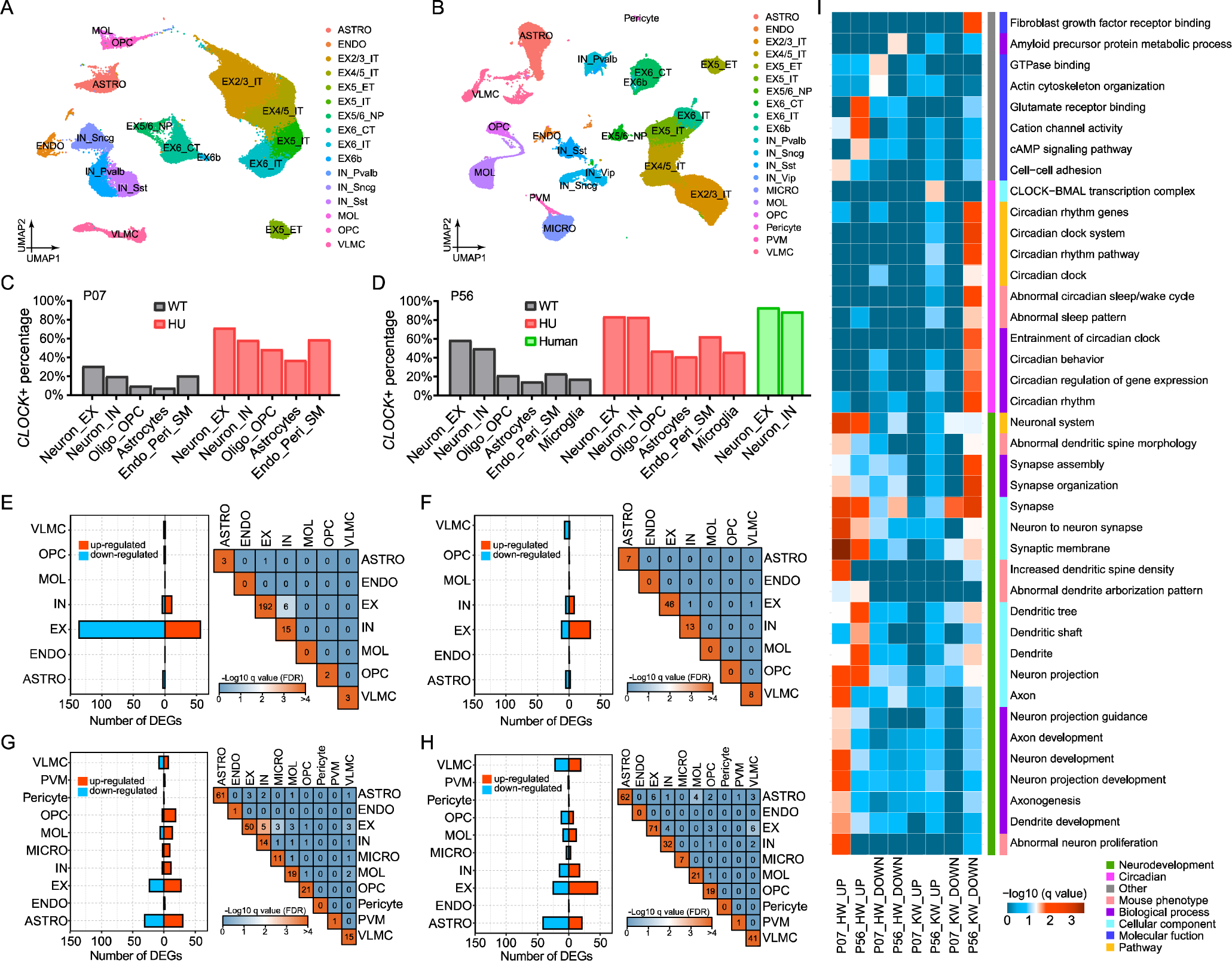
snRNA-Seq reveals that human CLOCK alters cell type-specific expression patterns in frontal cortex. **(A-B)** Uniform manifold approximation and projection (UMAP) plots show the major cell types of frontal cortex at **(A)** PO7 and **(B)** P56. ASTRO: astrocyte; ENDO: endothelium; EX2/3_IT, EX4/5_IT, EX5_IT, and EX6_IT: intratelencephalic projection neurons in layers 2-3, 4, 5, and 6, respectively; EX5_ET: extratelencephalic projection neurons in layer 5; EX5/6_NP: near projection neurons in layers 5 and 6; EX6_CT: corticothalamic projection neurons in layer 6; EX6b: excitatory neurons in layer 6b; IN_Pvalb, IN_Sncg, IN_Sst, and IN_Vip: inhibitory neurons subtypes exclusively expressing Pvalb, Sncg, Sst, and Vip, respectively; MICRO: microglia; MOL: mature oligodendrocyte; OPC: oligodendrocyte precursor cell; PVM: perivascular macrophage; VLMC: vascular leptomeningeal cell. **(C-D)** The percentage of CLOCK/Clock+ cells in each major cell type at **(C)** PO7 and **(D)** P56 based on snRNA-seq data. **(E-H)** The number of differentially expressed genes (DEGs) in each major cell type (bar plot), and hypergeometric tests between DEGs from each pair of cell types (heatmap) at **(E)** PO7 HU vs. WT comparison, **(F)** PO7 KO vs. WT comparison, **(G)** P56 HU vs. WT comparison, and **(H)** P56 KO vs. WT comparison. The color of each cell represents the −Log 10 of FDR corrected p value (q value). The number in each cell is the number of overlapped DEGs between two cell types. (I) Heatmap of gene ontology (GO) analysis to show enrichment of GO terms of the DEGs in EX2/3_IT. The color of each cell represents −Log10 q value of hypergeometric tests. GO terms are grouped based on their functions (left strip) and GO categories (right strip). HW_UP, human CLOCK upregulated genes; HW_DOWN, human CLOCK downregulated genes; KO_UP, Clock KO upregulated genes; KO_DOWN, Clock KO downregulated genes. See also **Figures S4**-**9**, **Tables S4-8** and **Table S13**.

The fraction of cells that express human CLOCK (CLOCK+ cells) in each major cell type are greater than the fraction of cells that express mouse CLOCK (i.e. in WT) (**Fig. 4C,D**). The results at P56 are consistent with our findings in IHC experiments (**Fig. 1E-H**), and the fraction of CLOCK+ neurons in HU mice is comparable with previous single-cell snRNA-seq data from human cortex (**Fig. 4D**) (Boldog et al., 2018). We also found that normalized expression of human CLOCK is greater than mouse CLOCK in all cell types at both P07 and P56 (**Fig. S6A,B**), this result is similar to the comparison between adult human and mouse across major cell types in cortex (Bakken et al., 2021). These results further confirm that HU mice mimic the expression pattern of CLOCK observed in human cortex. From P07 to P56, the fraction of CLOCK+ cells across the major cell types increased in both HU and WT mice (**Fig. 4C,D**). This result suggests that CLOCK might play different roles at different developmental stages. At both P07 and P56, the percentage of CLOCK+ cells varied across cell types, and neurons, especially excitatory neurons, showed the greatest percentage of both human and mouse CLOCK+ cells (**Fig. 4C,D**). Thus, CLOCK might have a major impact on excitatory neuronal function.

### CLOCK regulates gene expression in a cell type-specific manner

To identify the genes regulated by CLOCK, we did pairwise (i.e., HU vs. WT and WT vs. KO) comparisons within each major cell type at P07 and P56 separately. The HU vs. WT comparison detected genes that are specifically regulated by human CLOCK, while the WT vs. KO comparison determines genes that are regulated by mouse CLOCK. We overlapped the differentially expressed genes (DEGs) of each cell type at each age, and found that none of the cell types, except for excitatory and inhibitory neurons in HU vs. WT at P56 showed significantly overlapped DEGs (**Fig. 4E-H**; **Table S5-8**). These results suggest that CLOCK regulates gene expression in a cell type-specific manner. At P07, CLOCK primarily regulates genes in excitatory neurons (**Fig. 4E,F**). At P56, CLOCK has broader effects on gene expression across cell types, although excitatory neurons still show the greatest number of DEGs (**Fig. 4G,H**). These results, together with our finding that excitatory neurons are the cells with the greatest percentage of CLOCK+ cells (**Fig. 4C,D**) suggest that CLOCK might alter brain function mainly through regulation of gene expression in excitatory neurons.

Excitatory neurons can be divided into subtypes based on their projection patterns. To further narrow down to the subtype of excitatory neuron that is most affected by CLOCK expression, we identified DEGs for subtypes similar to what we did for major cell types. Projection neurons subtypes have distinct DEGs, while intratelencephalically projecting excitatory neurons (IT) from different cortical layers have significantly overlapping DEGs (**Fig. S7**; **Table S9-12**). At both P07 and P56, the greatest number of DEGs is in IT neurons, especially in layer 2-3 IT neurons (EX2/3_IT) (**Fig. S7**). This indicates that CLOCK might alter the cognitive behavior of mice mainly through regulation of gene expression of EX2/3_IT. Therefore, we focused on EX2/3_IT in the following analyses.

### Human CLOCK and mouse Clock regulate distinct sets of genes in EX2/3_IT in an age-specific manner

To investigate whether human and mouse CLOCK regulate different genes in EX2/3_IT, we distinguished up-and down-regulated DEGs, and then overlapped human and mouse CLOCK-regulated genes within age groups (**Fig. S8A,B**). The non-significant overlaps suggest that the expression of genes regulated by mouse CLOCK are rescued by human CLOCK, consistent with our observation of behavior (**Fig. S1D-J**). In addition, this lack of overlap suggests that overexpression of human CLOCK not only regulates and can rescue targets of mouse CLOCK but can regulate novel human CLOCK-specific genes. We also overlapped DEGs from the two time points, P07 and P56 (**Fig. S8C,D**). We found few overlapping genes, suggesting that both human and mouse CLOCK regulate genes in an age-specific manner.

To understand the function of the DEGs, we carried out Gene Ontology (GO) enrichment analysis on genes regulated by CLOCK in EX2/3_IT. Human CLOCK upregulated genes at both P07 and P56 are enriched for neurodevelopmental functions such as dendritic growth (e.g., neuron projection development) and spine formation (e.g., synapse organization) (**Fig. 4I**; **Table S13**). It suggests that these neurodevelopmental functions might be enhanced in HU mice. Consistent with the greater number of DEGs for HU at P07, the number of enriched GO terms for HU DEGs at P07 are more than those enriched at P56 (**Fig. 4I**), suggesting a more important role for CLOCK in early postnatal stage. In contrast, mouse KO led to enrichment of downregulated genes for only a few neurodevelopmental functions at P56 (**Fig. 4I**). In addition, DEGs of mouse KO at P56 but not P07 are enriched for circadian and sleep signal pathways (**Fig. 4I**). These results are consistent with a previous finding that showed that mouse visual cortex lacks rhythmic expression of circadian genes before eyes open at P10 (Kobayashi et al., 2015).

### Human CLOCK regulates genes with human-specific open chromatin

To investigate whether human CLOCK regulates human-specific expressed genes, we utilized a dataset that compared chromatin state from single-nuclei ATAC-seq in adult postmortem human cortex compared to chimpanzee and rhesus macaque (Emre et al., unpublished). We subsetted the dataset to include those genes with human-specific open patterns of chromatin state that also contained a CLOCK:BMAL1 binding motif and overlapped them with DEGs from HU mice in EX2/3_IT. We found a significant overlap (**Fig. S9A**), and 13 out of the 17 overlapping genes showed chromatin states consistent with their regulatory directions in HU mice (**Table S14**).

Some of these human-specific expressed genes are specialized for neurodevelopmental functions. For example, *Tenm2*, which is upregulated with greatest fold change in most of major cell types of P07 HU mice, enables cell adhesion molecule binding and facilitates axon growth (del Toro et al., 2020; Li et al., 2018). *Sorcs2*, which is upregulated in P56 HU mice, responds to the neurotrophic factor BDNF to maintain neurite growth and spine formation (Giza et al., 2018; Glerup et al., 2016). Finally, *Srgap2*, which is downregulated in P07 HU mice, inhibits dendrite branching and synaptogenesis (Charrier et al., 2012). Human-specific duplication of SRGAP2C suppresses activity of SRGAP2, resulting in complex dendrites and a greater density of spines in human (Schmidt et al., 2019; Schmidt et al., 2021).

We then validated the expression of TENM2 and SORCS2 proteins in the frontal cortex of P07 and P56 mice, respectively. We quantified the fluorescent intensity of antibody staining in IHC, and found that HU mice showed significantly greater expression of both proteins compared to WT mice (**Fig. S9B-F**). We examined the regulatory regions of both genes and detected CLOCK:BMAL1 binding motifs in the enhancer regions of both human and mouse genomes (**Table S15**) (Moore et al., 2020). Together, these results confirmed the snRNA-seq data and suggest that TENM2 and SORCS2 are direct downstream targets of CLOCK.

### Human CLOCK increases dendrite complexity and spine density

To assess potential neuronal morphological changes that are suggested by the snRNA-seq DEGs related to dendrites and spines, we sparsely transduced cortical neurons with a recombinant adeno-associated virus (rAAV) expressing mCherry through unilateral intracerebroventricular injection in P0-P1 pups to visualize cell morphology including the soma, neurite arborization, and synaptic spines. At P18 and P56, we performed IHC to amplify the mCherry signal and co-stain with cortical layer and cell type markers (**Fig. S10A,B**). We analyzed the morphology of the excitatory neurons in layers 2-4 of the frontal cortex. We found no difference in soma size across genotypes at both ages (**Fig. S11A** and **Fig. S12A**). We then quantified the complexity of dendritic trees in P56 mice. Compared with WT mice, HU mice have a greater number of branches and longer total dendritic length, while KO mice did not show a significant difference in either parameter (**Fig. 5A-C**). Sholl analysis further confirmed that the concentric rings encountered more intersections with the neuron branches in HU mice and fewer intersections in KO mice compared to WT mice (**Fig. 5D**). This is consistent with previous studies where neurons in brain tissue of epilepsy patients that expressed less CLOCK showed less complexity of dendritic arborization (Li et al., 2017). To summarize, our results suggest that CLOCK facilitates dendritic complexity of excitatory neurons in layers 2-4 of frontal cortex.

**Figure 5.**
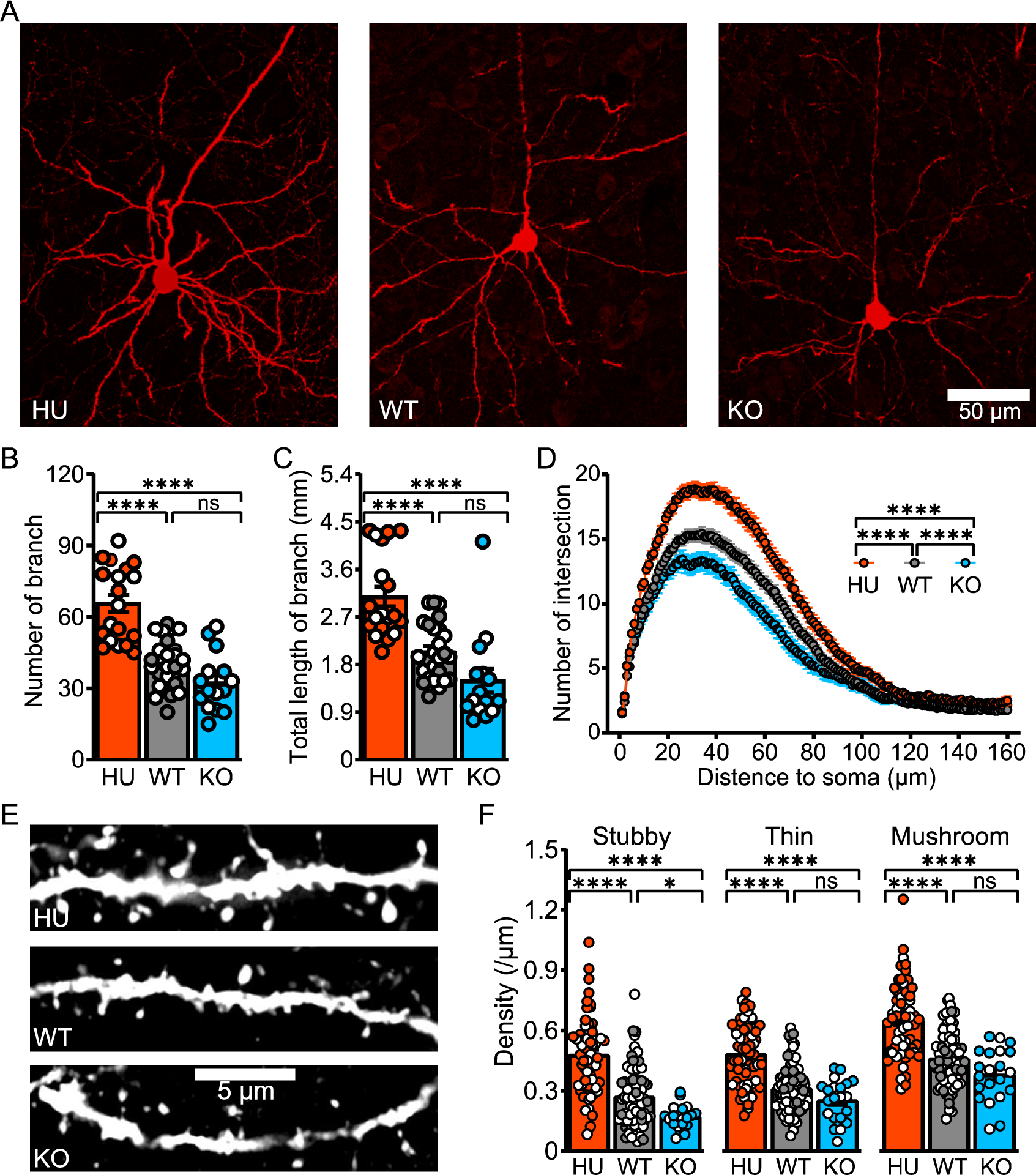
CLOCK increases dendrite complexity and spine density in adult mice. **(A)** Representative layer 2-4 excitatory neurons of frontal cortex in each genotype. **(B-C)** Quantification of **(B)** number and **(C)** total length of branch in each genotype. Each data point is a neuron. We sampled 3 mice per genotype and 5-9 neurons per mouse. We applied GLMM with genotype and sex as fixed factors, neuron as random factor nested with individual, and Tukey’s test for post-hoc analysis. **(D)** Sholl analysis to quantify encountered intersections between neuron branches and concentric rings from soma as measures of dendrite complexity. We did repeated measures ANOVA with genotype and sex as between-subject factors, distance to soma as within-subject factor, and Tukey’s test for post-hoc analysis. **(E)** Zoomed in segments to quantify density and morphology of spines. **(F)** Density of spines for each subtype of spine. Each data point is a segment, we sampled 2-4 segments from 3-8 neurons per mouse. We applied GLMM with genotype and sex as fixed factors, segment as random factor nested with neuron which nested with individual, and Tukey’s test for post-hoc analysis. See also **Figure S10, S11**, and **S12**.

We next quantified the density and morphology (stubby, thin, and mushroom) of spines in P56 mice. We measured the spine length in all spine types and the area of the spine head and width of the neck in thin and mushroom spines. Compared with WT mice, HU mice showed an increase in the density of all three types of spines, whereas KO mice instead demonstrated a decreased density of stubby spines (**Fig. 5E,F**). The results from KO mice are consistent with previous findings in *Emx-Cre Clock* KO mice and *Clock* knockdown in primary neuronal cultures (Li et al., 2017). A comparison of spine morphology did not detect significant differences across genotypes except for the elongated thin and mushroom spines in HU mice (**Fig. S11B-H**). Thus, these results indicate that CLOCK mainly affects spine density.

To determine whether the more complex dendritic trees and greater spine density results from an increased generation or decreased pruning of dendrites and spines, we also measured the spine properties at P18, which is before the peak of synaptogenesis at P21 (Micheva and Beaulieu, 1996; Semple et al., 2013). We found results similar to what we observed at P56 (**Fig. S12**). Thus, CLOCK is more likely to drive neuronal projection growth and spine formation rather than neurite elimination and synaptic pruning. Given the pervasive expression of CLOCK in the whole cortex (Kobayashi et al., 2015; Li et al., 2017), we also quantified all rAAV transduced excitatory neurons regardless of their cortical layers and found similar results to those in layer 2-4 neurons (data not shown). In summary, these results are consistent with our snRNA-seq DEGs and demonstrate that CLOCK facilitates neurite growth and spine formation of cortical excitatory neurons.

### Humanized mice show greater frequency of spontaneous EPSCs

To determine whether the increased complexity of dendrites and spine density result in enhanced neural connectivity in HU mice, we characterized the electrophysiological properties of layer 2-4 excitatory neurons in the somatosensory cortex of 3–4-week-old mice (**Fig. S13A**). We found that the firing versus current injection curves were similar across genotypes **(Fig. S13B)**, suggesting that the excitatory neurons do not have differences in intrinsic excitability. The subthreshold membrane properties (i.e., resting membrane potential (RMV), input resistance, and normalized conductance) also did not show any differences **(Fig. S13C-E)**. These results demonstrate that CLOCK does not affect the intrinsic properties of excitatory neurons. Similar results were found in inhibitory parvalbumin neurons of the visual cortex that had *Clock* deleted (Kobayashi et al., 2015).

We then measured the spontaneous and evoked excitatory postsynaptic currents (EPSCs) to determine if CLOCK changes the overall neuronal network. Neurons in HU mice showed an increased frequency but decreased amplitude of spontaneous EPSCs (**Fig. 6A-C**). The decreased amplitude could be attributed to a compensatory mechanism to the increased frequency (Turrigiano, 2012). Thus, we calculated the total charge transfer (frequency x amplitude) to represent the overall effect and found that it is greater in neurons from HU mice compared to WT mice (**Fig. 6D**). This result might be linked to *Nrxn2*, a gene that is associated with spontaneous EPSC frequency (Missler et al., 2003; Ramirez and Kavalali, 2011), and one of the DEGs that is upregulated in HU mice. We then measured evoked EPSCs from upper-layer excitatory cortical neurons by stimulating extracellular electrode placed in the same cortical layer (**Fig. S13F**). Consistent with the decrease in total charge transfer of spontaneous EPSCs, evoked EPSC amplitude was decreased in KO mice while there was no difference between HU and WT mice (**Fig. 6E**). This finding is consistent with the downregulation of *Snap25 in* KO mice, which is one of the plasma membrane proteins for synaptic vesicle fusion in evoked EPSCs (Südhof and Rothman, 2009). In summary, our results indicate that the increased frequency of spontaneous EPSCs in HU mice was neither because of altered intrinsic neuronal excitability nor due to greater vesicle release. Therefore, the most plausible explanation is enhanced neural connectivity in HU mice, which is consistent with our results from snRNA-seq and neuroanatomical measures.

**Figure 6.**
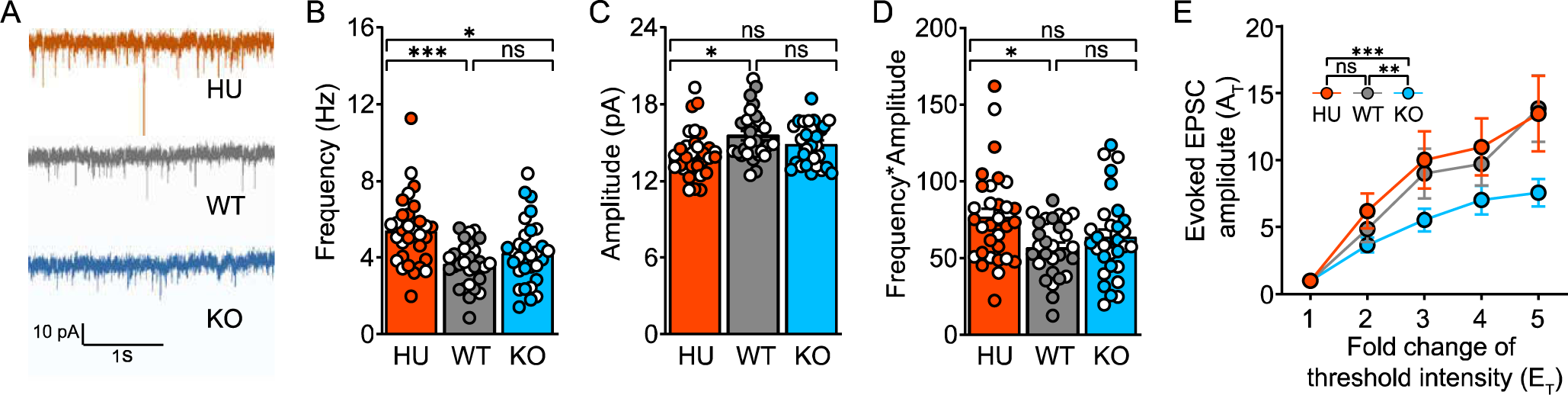
CLOCK alters EPSCs of layer 2-4 excitatory neurons. **(A)** Representative raw traces of spontaneous EPSCs were recorded from the primary somatosensory cortex of layer 2-4 excitatory neurons. **(B-D)** Comparison of **(B)** frequency, **(C)** amplitude, and **(D)** frequency*amplitude of spontaneous EPSCs for each genotype. Each data point is a neuron. We recorded from 5-8 neurons per mouse where 4-5 mice per genotype were used. **(E)** The amplitude of evoked EPSCs in response to the stimulus with a fold change of threshold intensity. We applied GLMM with genotype, sex, and stimulus intensity (x-axis for panel **E** only) as fixed factors, neuron as a random factor nested with individual, and Tukey’s test for post-hoc analysis. See also **Figure S13**.

### TENM2 overexpression rescues human iPSC-derived neurons with CLOCK knockout

We next wanted to investigate if human CLOCK exerts similar functions in human neurons. To determine whether the neuronal alterations in HU mice emulate CLOCK-associated phenotypes in human neurons, we applied CRISPR-Cas9 to delete 1-17 bp of nucleotides at Exon 7 of *CLOCK* to induce a frameshift in induced pluripotent stem cell (iPSC) clones (**Fig. S14A**). We selected clone #2 and #7, which represented two types of altered reading frames with the longest deletions. The results of qRT-PCR and western blot confirmed that CLOCK was not expressed in the KO clones and the core-circadian downstream targets were altered in expression (**Fig. S14B,C**), which suggested a successful knockout. We then followed an NGN2-overexpression protocol to rapidly differentiate the iPSCs into monolayers of excitatory cortical neurons (Ho et al., 2016; Zhang et al., 2013) (**Fig. S14D**). We did immunocytochemistry (ICC) on Day 28 of differentiation for TUJ1 and MAP2, which indicated the successful differentiation process of iPSCs to mature neurons (**Fig. 7A** and **Fig. S15**), while CAMKII staining indicated that these neurons were excitatory neurons (**Fig. S15A,B**). Positive staining of CUX1 but not FOXP2, suggests that the differentiated neurons are more similar to the properties of upper-layer neurons (layers 2-4) rather than deep layer (layer 6) cortical neurons (**Fig. S15C**). We quantified the length and segment of cell projections expressing TUJ1 and the density of puncta expressing VGLUT2 to analyze the dendritic complexity and spine density of the iPSCs-induced neurons respectively. We discovered that CLOCK KO neurons have decreased number of segments and density of puncta (**Fig. 7A-D**). These results are consistent with what we found in mouse excitatory neurons (**Fig. 5**) and suggest that CLOCK is functionally involved in the development of dendrites and spines in the human excitatory neurons.

**Figure 7.**
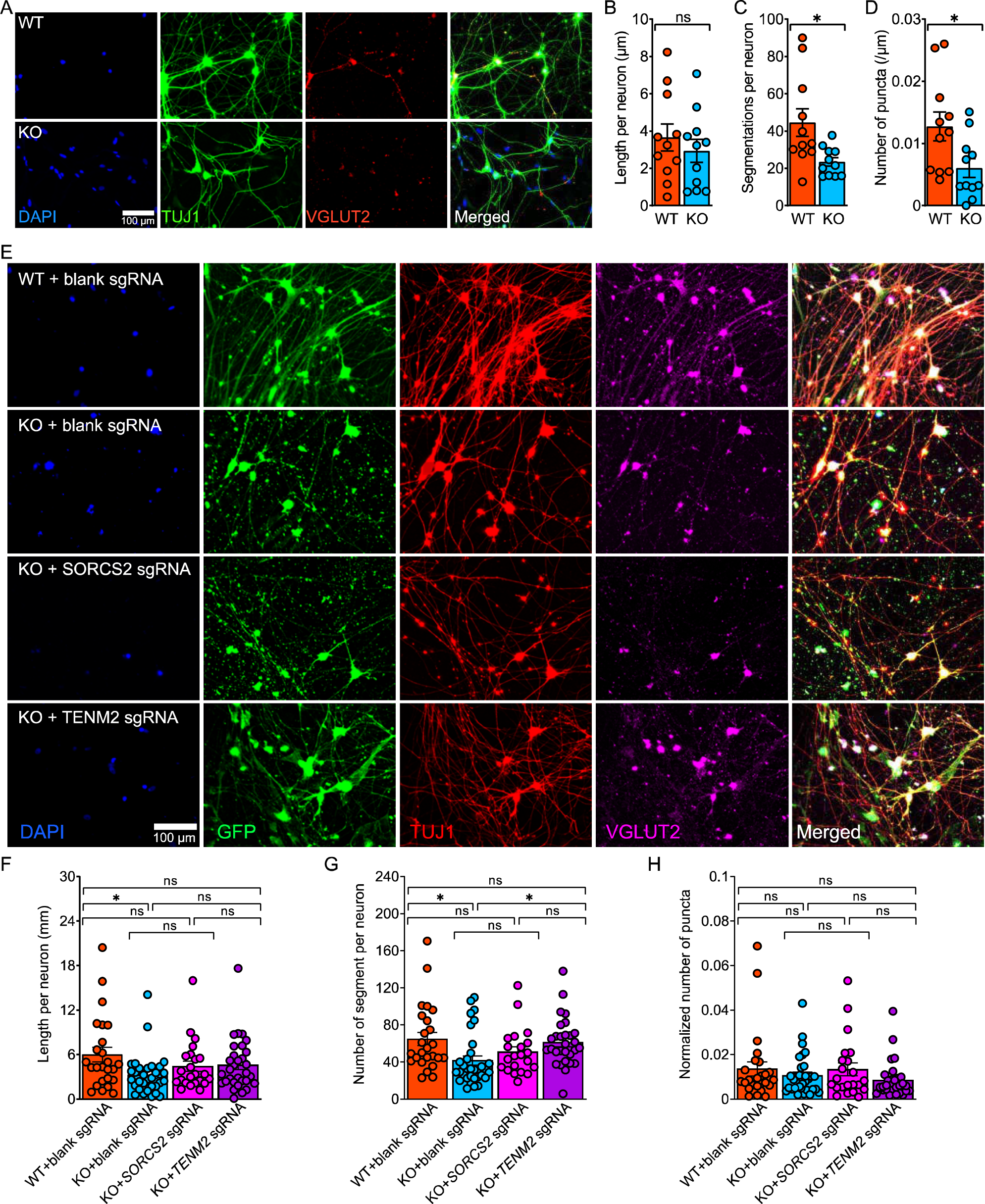
CLOCK KO reduces mature neuronal features in iPSC-derived neurons and is partially rescued by TENM2. **(A)** Representative ICC images of iPSC-derived neurons at Day 28 with nuclei marker DAPI (blue), neuronal dendrite mark TUJ1 (green), and presynaptic marker VGLUT2 (red). **(B)** and **(F)** Total length of dendrites, which was quantified from TUJ1 staining, normalized by the number of nuclei overlapped with TUJ1 signals. **(C)** and **(G)** Number of TUJ1 segments normalized by the number of nuclei overlapped with TUJ1 signals as a quantification of dendrite complexity. **(D)** and **(H)** Number of VGLUT2 puncta overlapped with TUJ1 signal and normalized by length of dendrites as a quantification of presynaptic spine density. **(B-D)** Quantification of iPSC-derived neurons at Day 28. **(E)** Representative ICC images of iPSC-derived neurons at Day 35 with nuclei marker DAPI (blue), CRISPR-Cas9 transfection marker GFP (green), neuronal dendrite mark TUJ1 (red), and presynaptic marker VGLUT2 (magenta). **(F-H)** Quantification on iPSC-derived neurons at Day 35. Each data point is a culture of neurons on a circle glass cover slip sitting on the bottom of a well in a 24-well plate. We did GLM with genotype/treatment as fixed factors with Tukey’s test for post-hoc analysis. See also **Figure S14** and **S15**.

We then tested whether the phenotypes of *CLOCK* KO in the human iPSCs-induced neurons could be rescued by overexpressing either of the two validated downstream targets of CLOCK (*TENM2* and *SORCS2*). We leveraged CRISPR/Cas9 to target the promoter region of *TENM2* and *SORCS2* for transcriptional activation (Matharu et al., 2019). qRT-PCR showed that *CLOCK* KO neurons decreased expression of both genes, while CRISPR-mediated activation resulted in similar expression levels of both genes as WT (**Fig. S14G**). We then quantified ICC results on Day 35 for dendritic complexity and puncta density. The results of *CLOCK* KO and WT neurons with empty plasmid (**Fig. 7E-H**) replicated the results of untreated neurons of Day 28 for dendritic arborization (**Fig. 7A-C**). The KO neurons with *TENM2* sgRNA were partially rescued, showing a more complex arborization than KO neurons (**Fig. 7E,G**). However, *TENM2* overexpression failed to rescue the decreased puncta in KO neurons (**Fig. 7E,H**), while *SORCS2* upregulation failed to rescue either phenotype (**Fig. 7E-H**). The monolayer iPSCs-derived neurons mainly capture the early developmental stage (Odawara et al., 2014). This could explain why *TENM2*, which is regulated by CLOCK at P7 in mice, could partially rescue dendritic arborization but *SORCS2*, a gene that is regulated by CLOCK in adult mice, could not rescue dendritic arborization.

## Discussion

In this study, we generated a humanized mouse model to mimic the neocortical expression of human CLOCK and demonstrate a robust extra-circadian function for human CLOCK in directing brain development and function. We find that human CLOCK regulates genes with human-specific expression to enhance dendritic growth and spine formation in EX2/3_IT. Moreover, the increased dendritic branches and synaptic spines in EX2/3_IT neurons likely facilitate the enhanced functional connections that we observe as a greater frequency of spontaneous EPSCs. Together, these genomic and physiological alterations downstream of human CLOCK present as the likely mechanism for the improved cognitive flexibility of HU mice.

It was unclear from previous studies, whether the observed increase in overall expression of CLOCK in the human cortex (Babbitt et al., 2010; Bozek et al., 2014; Konopka et al., 2012) could be attributed to higher cellular expression of CLOCK and/or a greater fraction of CLOCK+ cells. Using HU mice, we found that human CLOCK showed an increase in both measures of expression compared to mouse Clock. However, MS mice, which have increased cellular expression of mouse CLOCK but with similar percentages of CLOCK+ cell as WT mice, showed similar performance in reversal learning and cortical cell density as WT mice. These comparisons suggest that the possible cell-type specific expression of CLOCK rather than relative level of expression might determine the extra-circadian function of CLOCK in cortical excitatory neurons.

CLOCK also exhibited a strong temporal effect on neuronal functions. We found that mouse CLOCK regulated genes associated with circadian rhythms and some neurodevelopmental functions at P56, but we observed very few DEGs downstream of mouse CLOCK alteration at P07 (**Fig. 4I**). In contrast, human CLOCK regulated 192 genes in excitatory neurons at P07 (**Fig. 4E**). Moreover, the results of GO analysis of DEGs from EX2/3_IT highlighted dendrite growth and spine formation (**Fig. 4I**), which is consistent with the developmental events around P07 (Baloch et al., 2008; Semple et al., 2013). There are more than twice as many CLOCK+ excitatory neurons in HU compared to WT as P07 (**Fig. 4C**), which might be a mechanism driving gained neurodevelopmental functions.

Although functional alterations of human CLOCK could be attributed to its change of spatiotemporal expression, our study could not exclude the possibility that human CLOCK evolved novel molecular functions. Compared to human, CLOCK proteins are 98.7% and 90.3% identical in chimpanzee and mouse respectively (Sayers et al., 2021; Wilsbacher et al., 2000). Since the main functional binding domains (i.e. bHLH and PAS) in the N-terminal of CLOCK are highly conserved, a major change in functions is not expected. Species coding differences mainly occur in the C-terminal PolyQ region of CLOCK, and the function of this region is still controversial (O’Brien et al., 2013; Saino et al., 2015). A lack of polymorphisms in the human CLOCK PolyQ region suggests purifying selection (Saleem et al., 2001), but its function with respect to human brain evolution still needs elucidating.

A hallmark of human brain evolution is a greater complexity of dendrites and density of spines compared to other non-human primates (Bianchi et al., 2013; Elston et al., 2001; Schmidt and Polleux, 2022; Sherwood et al., 2020). Increased complexity of neural connectivity has also been linked to the altered computational properties of the human brain and might result in enhanced cognitive abilities (Hayashi and Majewska, 2005; Schmidt and Polleux, 2022; Schmidt et al., 2021). Our snRNA-seq data, dendritic morphology measures, and electrophysiology data consistently suggest that human CLOCK enhances neural connectivity in the frontal cortex of HU mice. Previous studies demonstrated that the complexity of dendrites and density of spines are positively correlated with rodents’ performance in cognitive tests (Leggio et al., 2005; Leuner and Gould, 2010). Therefore, the observed enhancement of cognitive flexibility in HU mice might result from the increased neural connectivity.

In addition to the increased neural connectivity of excitatory neurons, we also observed an increased neuron density in HU mice (**Fig. 3C**). The more complex neural connections of the human neocortex are also predicated on an increased abundance of neurons (Dicke and Roth, 2016; Elston et al., 2006). These data suggest that human CLOCK might regulate the proliferation of neural progenitors during embryonic development. Dividing cells have rhythmic expression of circadian proteins (Nagoshi et al., 2004). Some cell cycle genes (e.g. *c-Myc* and *Wee1*) are under the control of circadian genes such as *Cry1*, *Cry2*, *Per1*, and *Clock* (Fu et al., 2002; Matsuo et al., 2003; Miller et al., 2007). The greatest CLOCK expression levels are detected in progenitor-like cells in early embryonic development (Loo et al., 2019). Thus, future studies could investigate whether human CLOCK affects the proliferation of neural progenitors at embryonic stages.

In this study, differential gene expression in circadian-related genes was only detected in KO mice at P56, not HU mice. These results suggest that human CLOCK might regulate neural functions independently of circadian related pathways. Extra-circadian functions have also been suggested for CLOCK and other core circadian genes such as BMAL1 (Lipton et al., 2015). *Clock* and *Bmal1* mutant mice but not *Cry1* or *Cry2* mutant mice have deficits in place learning (Kondratova et al., 2010; Van der Zee et al., 2008). Similarly, astrogliosis was found in the cortex of aged *Npas2* and *Clock* double KO and *Bmal1* KO mice, while *Per1* and *Per2* double KO mice did not show this phenotype (Musiek et al., 2013). *Per1*, *Per2*, *Cry1* and *Cry2* are downstream targets of CLOCK and BMAL1 in the circadian signal pathway (Takahashi, 2017). These results suggest that CLOCK regulates place learning and astrogliosis through a non-circadian pathway, and therefore further demonstrated the extra-circadian function of CLOCK.

Human-specific changes, which are those traits that evolved after divergence from the common ancestor of chimpanzee and humans, are of great interest for understanding evolution (Konopka and Geschwind, 2010; Pattabiraman et al., 2020). Given the phylogenetic distance between mouse and human, however, we should be cautious that a humanized mouse model would inevitably capture primate-specific, ape-specific, and human-specific alterations. The other limitation of this study is that the human CLOCK gene was expressed in the background of the mouse genome. Given the high homogeneity of genomes between human and mouse, we found that gene expression, neuroanatomy, and cognition of HU mice captured some human-specific alterations. However, the mechanism we uncovered in humanized CLOCK mice might not be the same as in vivo human specializations due to the mouse background. This could be addressed by generating a non-human primate model system or studying some of the cellular and molecular mechanisms by using human and chimpanzee iPSCs systems.

In conclusion, we found that the spatiotemporal alteration of human CLOCK expression in a mammalian model system results in human-specific expression of genes that enhance dendritic growth and spine formation of excitatory neurons. The observed increased density of neurons and complexity of dendrites and spines form enhanced functional connectivity in the frontal cortex. These changes may underlie the advanced cognitive flexibility of the humanized mice. Our study suggests that gain of extra-circadian functions of CLOCK via altered spatiotemporal gene expression might be an important molecular mechanism for human brain evolution.

## Methods

### Mouse

All mice in this study were kept in C57BL/6J genetic background and housed in 12hr light/12hr dark cycle (LD), lights were on at 7 am Central Standard Time. All mice were raised at University of Texas Southwestern Medical Center (UTSW). We complied guidelines of UTSW Animal Care and Use Committees and followed procedures approved by UTSW Institutional Animal Care and Use Committee (IACUCC# 2015-100918) for animal care and experiments. We weaned pups around 4 weeks of age and housed with same-sex littermates for experiments.

### Cell culture

All cultured cells were kept at 37°C, 5% CO_2_, and above 90% humidity.

#### HEK293T cells

HEK293T cells (ATCC, #CRL-3216) were cultured in DMEM (Cytiva HyClone^TM^, #SH3024301) with 1% Antibiotic-Antimycotic (Thermo Fisher Scientific, #15240062) and 10% Fetal Bovine Serum (FBS) (Thermo Fisher Scientific, #10437028).

#### Human iPSC culture

We obtained a human iPSC line (WTC-11, #GM25256) from the Gladstone Institute. We cultured the cells following the Technical Manual (Stemcell Technologies, Document #28315). Cells were maintained in six-well plates (Corning, #4936) with mTeSR^TM^1 medium (Stemcell Technologies, #85870). The plates were coated with growth factor-reduced Matrigel matrix hESC-qualified (BD Biosciences, #354277). We passaged cells when the confluency of culture reached about 85% by using Gentle Cell Dissociation Medium (Stemcell Technologies, #07174) and ROCK inhibitor (Fisher Scientific, #50-863-6). We changed mTeSR^TM^1 medium every day to maintain the cell line.

#### Rat primary astrocytes

Rat primary cortical astrocytes (Invitrogen, #N7745-100) were cultured according to Invitrogen protocols (MAN0001679 for thawing and establishing culture; MAN0001680 for expanding culture). We cultured astrocytes in Matrigel coated 6-well plate with DMEM and 15% FBS medium. Medium was half changed every 5 days. We passaged astrocyte cultures when they reached 100% confluency using 1:1 1X PBS (Cytiva HyClone^TM^, #SH3025601) diluted Accutase (Gibco^TM^, #A1110501).

### Generation of humanized CLOCK mice

#### Recombineering human CLOCK BAC

We obtained a BAC (RP11-1029A14) that contained the full-length human CLOCK gene from BACPAC Resources (Children’s Hospital Oakland Research Institute). This BAC contains the full-length TMEM165 and part of PDCL2 genes flanking on the 3’ and 5’ end of CLOCK gene respectively (**Fig. 1A**). We first removed TMEM165 and PDCL2, and then we added a 3XFLAG-3XHA N-terminal tag through recombineering. The following primers were used to direct the recombination events (the underlined sequences are what binds the cassette, while the sequences before that are the 50bp homology arm overhangs):

5’ TMEM165 trim

Cassette Forward: 5’ CAAACAGGTATGA GGCATAATTCTGCTACTTACTAGCTGTGTAATTTGAGCCTGTTGACAATTAATCATCGGCA 3’

Cassette Reverse: 5’ ATTGGCCTATCAA TACTGATAATTTTAT ATGACACACAAGGAAAAACAAGTCAGCACTGTCCTGCTCCTT 3’

Oligo Forward: 5’ CAAACAGGTATGAGGCATAATTCTGCTACTTACTAGCTGTGTAATTTGAGCTTGTTTTTCCTTGTGTGTCATATAAAATTATCAGTATTGATAGGCCA AT 3’

Oligo Reverse: 5’ ATTGGCCTATCAATACTGATAATTTTATATGACACACAAGGAAAAACAAGCTCAAATTACACAGCTAGTAAGTAGCAGAATTATGC CTCATACCTGTTTG 3’

3’ PDCL2 trim

Cassette Forward: 5’ TTTCTGATCCCTAATATAAAAGCAGTCATAAGTAAAACATTCAGTACACACCTGTTGACAATTAATCATCGGCA 3’

Cassette Reverse: 5’ ATTCTGCTACATGCTACAACATAGTCGAACCTTGAAAACATGCTAAGCGATCAGCACTGTCCTGCTCCTT 3’

Oligo Forward: 5’ TTTCTGATCCCTAATATAAAAGCAGTCATAAGTAAAACATTCAGTACACATCGCTTAGCATGTTTTCAAGGTTCGACTATGTTGTAGCATGTAGCAGAAT 3’

Oligo Reverse: 5’ ATTCTGCTACATGCTACAACATAGTCGAACCTTGAAAACATGCTAAGCGATGTGTACTGAATGTTTTACTTATGACTGCTTTTATATTAGGGATCAGAAA 3’

N-terminal 3XFLAG-3XHA tag

Cassette Forward: 5’ TAAGGAGAAGTACAAATGTCTACTACAAGACGAAAACGTAGTATGTTATGCCTGTTGACAATTAATCATCGGCA 3’

Cassette Reverse: 5’ ACATACCTGTCAACAATCGAGCTCATTTTACTACAGCTTACGGTAAACAATCAGCACTGTCCTGCTCCTT 3’

Oligo Forward: 5’ TAAGGAGAAGTACAAATGTCTACTACAAGACGAAAACGTA GTATGTTATGGACTACAAAGACCATGACGG 3’

Oligo Reverse: 5’ ACATACCTGTCAACAATCGAGCTCAT 3’

#### Electroporation

We inoculated SW102 cells into 5ml LB Broth (Sigma, #L3022) with 25ug/ml chloramphenicol (Sigma, #C0378) and grew cells at 32°C overnight. We took 4mL of the saturated SW102 culture to inoculate into 120ml LB Broth with 25ug/ml chloramphenicol and grew the culture on a 32°C shaker to an OD600 between 0.2-0.3 for 2 hours. We placed the SW102 culture in a 42°C water bath for 15 minutes and then transferred to 50mL conical tubes and chilled on ice. The cultures were centrifuged at 4°C at 4000rpm for 10 minutes to pellet cells. The cell pellet was resuspended in 25mL ice-cold 10% glycerol (VWR, #EM-4750). These centrifuge-resuspension steps were repeated for two more times with ice-cold 10% glycerol. After decanting the supernatant of the last wash, the recombineered BAC DNA was electroporated into these cells using a 1mm cuvette at 1750 V, 25 uF, 200 Ohms. 1mL of LB Broth was added and the cells then recovered for 2 hours at 32°C with shaking. 1mL cultures were spun down and washed with minimal M9 salt wash. Finally, the cells were plated on minimal plates with 25ug/ml chloramphenicol and incubated at 32°C.

#### Cesium Chloride BAC preparation

We picked a single colony from the minimal plate and grew it in a 2ml LB Broth culture overnight at 32°C with chloramphenicol. We inoculated 2L LB Broth cultures and incubated ∼24 hours at 32°C with chloramphenicol. We harvested the bacteria by centrifugation at 4,000g for 20 minutes. The pellets were resuspended in 100ml Buffer P1 without RNase (45.5 ml 20% Glucose (Sigma, #G6152), 25ml 1M Tris pH8.0 (Fisher, #BP152-5), 20ml 0.5M EDTA (Sigma, #E-5134), ddH_2_O to 1L). We added 50ml Buffer P2 (20ml 10N NaOH, 50ml 20% SDS, ddH_2_O to 1L) and mixed by inversion, and then incubated at room temperature (RT) for 5 minutes. 37mL cold Buffer P3 to the lysate (600ml 5M Potassium Acetate (Sigma, #P1190), 115ml Glacial Acetic Acid (Sigma, #695092), ddH_2_O to 1L) was then added, mixed immediately 4∼6 times, and placed on ice for 10 minutes. Samples were centrifuged at 10,000 rpm for 20 minutes, then the supernatant was transferred to new tubes and 100ml isopropanol was added. We then centrifuged the sample at 10,000rpm for 30 minutes, followed by resuspending the pellet in 7ml TE. We then added 1.1g CsCl per ml of TE and 300ul Ethidium Bromide (EtBr; Fisher, #BP1302). The sample was separated into phases in a VTI 80 ultracentrifuge overnight at 95,000 rpm at 22°C. We then pulled the EtBr-marked DNA band with a 3ml syringe. The remainder was filled with TE mixed with 1.1g CsCl per ml of TE, and spun again for 4 hours at 95,000 rpm at 22°C. The BAC DNA band was pulled again and EtBr was removed with repeated 3ml washes of n-Butanol saturated with ddH2O. The resulting mix was diluted with 3X volume of TE, and 1/10 volume of 3M Sodium Acetate (Sigma, #71183) was added. 2X volume of 100% ethanol was added and samples were chilled on ice for 20+ minutes. DNA was pelleted at 4,500 rpm for 30 minutes at 4°C. The final ultrapure BAC DNA was diluted in BAC injection buffer (10mM Tris pH 7.4, 0.25mM EDTA).

#### Selection of mouse line

The CsCl purified human CLOCK BAC was integrated into C57BL/6N mice by direct pronuclear injection. Since we intended to generate a mouse model to mimic overexpression of human CLOCK, founders were screened by qRT-PCR, which will be described later, to generically amplify both human CLOCK and mouse Clock. Four founders showed increased *CLOCK* expression in all three brain regions (i.e. frontal cortex, hippocampus, and cerebellum, **Fig. S1A**). We then did an immunoprecipitation (IP) using cortex from the mice as previously described (Usui et al., 2017b). We used primary CLOCK antibody (MilliporeSigma, #ab2203) in 1:100 dilution in each sample. We then did Western Blotting on the IP’d sample as previously described (Usui et al., 2017a) using both primary FLAG antibody (Sigma, #F1804) in 1:400 dilution and a different primary CLOCK antibody (Abcam, #ab3517) in 1:200 dilution. The result showed that only three founders demonstrated bands representing FLAG-tagged human CLOCK at the expected size (**Fig. S1B**). The three founder lines (HUf2, HUf3, and HUf7) were kept and backcrossed onto a C57BL/6J genetic background for 10 generations. SNP genotyping was performed the same as described in previous work on the 5th generation of backcross (Kumar et al., 2012). It only detected about 8% of SNPs are heterogenous, while the other loci all showed C57BL/6J SNPs (**Table S1**). In our breeding, we found that the human CLOCK BACs of HUf2 were likely inserted into X chromosome. Thus, we used lines HUf3 and HUf7 for our experiments. Given the relative higher expression of CLOCK in frontal cortex of HUf7, we used HUf7 mice for all of our experiments and used HUf3 mice to confirm the results of critical experiments.

### Breeding of mice for experiments

The humanized CLOCK mouse lines (HUf3 and HUf7), which were denoted as *CLOCK^hum^;Clock^+/+^*, were crossed with exon 5-6 Clock knockout mice (KO: *Clock^-/-^*), which were generated by the lab of David R. Weaver (DeBruyne et al., 2006). The F1 heterozygotes (*CLOCK^hum^;Clock^+/-^* and *Clock^+/-^*) were crossed to obtain littermates of humanized (HU: *CLOCK^hum^;Clock^-/-^*), KO, and wildtype (WT: *Clock^+/+^*) mice (**Fig. S1C**). Similar to the generation of HU mice through breeding, we crossed mouse Clock BAC line (*Clock^mus^;Clock^+/+^*), which we previously generated to overexpress mouse Clock (Antoch et al., 1997), with KO mice to generate heterozygotes and then obtained Clock BAC mouse without endogenous Clock (*Clock^mus^;Clock^-/-^*, MS).

### Genotyping of human CLOCK and mouse Clock KO

We extracted genomic DNA from an ear or tail snip. The PCR mixture contains 5 ul EmeraldAmp GT PCR Master Mix (Takara, #RR310B), 1 ul genomic DNA, 1ul 10uM primers mix and 3 ul molecular grade H_2_O. We amplified a 298bp fragment to detected human *CLOCK* with following primers: F: 5’ GTCGTCACCCTCGTTTGAGT and R: 5’ TTGCTGCATCCTACAGTGCT; and PCR cycle settings: 5 min at 95°C; 35 cycles of 15 sec at 95C, 30 sec at 60°C, and 30 sec at 68°C; 5 min at 72°C; hold at 12°C. We detected the WT and KO of endogenous mouse Clock by using the following primer set: F_WT_: 5’GGTCTATGCTTCCTGGTAACG, F_KO_: 5’ATTCCCCATCCAAAGATATTTGC, and R: 5’CCAGGCTTACGCTGAGAGC; and PCR cycle settings: 3 min at 94°C; 35 cycles of 30 sec at 94°C, 30 sec at 60°C, and 90 sec at 72°C; 10 min at 72°C; hold at 4°C to amplify 280bp and 506bp fragments for WT and KO allele respectively (DeBruyne et al., 2006). We amplified a ∼300bp fragment to detected mouse Clock BAC with following primers: F: 5’ ATACGACTCACTATAGGGCGAATTCG and R: 5’ TTAGCCTCTTCTGTGCTACATCG; and PCR cycle settings: 2 min at 96°C; 35 cycles of 15 sec at 95°C, 30 sec at 55°C, and 15 sec at 72°C; 3 min at 72°C; hold at 12°C. The PCR products were analyzed on 1% agarose gel. However, for genotyping of MS mice, the primers to detect endogenous mouse Clock could also amplify mouse Clock BAC, so MS (*Clock^mus^;Clock^-/-^*) and *Clock^mus^;Clock^+/-^* mice could not be distinguished by using above mentioned PCR tests. Therefore, we further distinguished the two genotypes by quantitatively determining the copy number of LoxP in Clock knocked out allele through quantitative real-time PCR (qRT-PCR).

### Tissue preparation and qRT-PCR

#### Quantification of copy number of LoxP for genotyping of MS mice

In order to distinguish MS mice (*Clock^mus^;Clock^-/-^*) from *Clock^mus^;Clock^+/-^* mice, We designed the primers (F: 5’ AGTGGGTGGCACCGATAAG and R: 5’ CACCAAATGTTTGTTGAATGAAA) to amplify a 166bp fragment containing a LoxP sequence in the Clock KO allele. We extracted DNA from 0.7 cm tail snips and performed phenol-chloroform purification (Sambrook and Russell, 2006), and then we diluted the DNA to 0.1 ng/ul. The DNA solution was further diluted 4X as the DNA template. We always ran these DNA samples with known KO, *Clock^+/-^*, and WT samples as a reference to determine copy number of LoxP. The qRT-PCR mixture contains 5ul iTaq Universal SYBR Green Supermix (Bio-Rad Laboratories, #172-5121), 2ul DNA template from phenol-chloroform purification, 1ul 10uM primers mix and 2ul molecular grade H_2_O in one well of a 384-well plate (MicroAmp™, # 4343370), each sample was run in replicated reactions in 4 wells. We performed qRT-PCR using a Touch Real-Time PCR Detection System (Bio-Rad Laboratories, #CFX384) with the following PCR cycle settings: 2 min at 50°C; 10 min at 95°C; 39 cycles of 15 sec at 95°C, 45 sec at 65°C; 5 sec at 65°C; to 95°C and stop. The Cq value of the 4 replicates from the same sample were averaged, and we compared the Cq values of the samples with the genotype-known reference samples to determine their genotypes.

#### TaqMan qRT-PCR quantification of BAC copy number

We did phenol-chloroform extraction of DNA from 0.7 cm tail snips of WT mice and then diluted them to 100 ng/ul as the genome background. We prepared the sample DNA template in the same way and concentration. Purified BAC solution was diluted to 2 ng/ul in BAC injection buffer. Then 1.684 ul of BAC solution was added to 40 ul of genome background solution. This would result in 16 BAC copies per diploid mouse genome. We further diluted the BAC copies to 8, 4, 2, 1 in the genome background solution for a standard curve. The qRT-PCR mixture contains 5 ul TaqMan™ Universal Master Mix II no UNG (Applied Biosystems, #4440040), 0.9 ul 10 uM primer mix (F: 5’ GCCAGCCCCTTCTTGTCA and R: 5’ ACTGCAGTCTGAGCGGTTAGTG), 0.25 ul 10 uM TaqMan probe (5’ TTAGATCCTAGCTCACGTGTC; Applied Biosystems, #5371391), 1 ul DNA template, and 2.85 ul molecular grade H_2_O in one well of a 384-well plate, each sample was run in replicated reactions in 4 wells. We performed qRT-PCR using the same platform with the following PCR cycle settings: 10 min at 95°C; 39 cycles of 15 sec at 95°C, 1 min at 60°C and stop. The Cq value of the 4 replicates from the same sample were averaged, and we generated a linear model between the Cq values and the known copy number per diploid mouse genome. Then we calculated the copy number of the samples by substituting Cq value into the model.

#### RNA isolation and quantification of gene expression

Mice were rapidly decapitated and brains quickly removed. The fresh brain was quickly transferred to ice-cold coronal Acrylic Mouse Matrice - 1mm Coronal (Braintree Scientific, # BS-A-5000C) and washed with ice-cold 1X PBS. The boundary between the olfactory bulb and frontal cortex was aligned to the first dent where the first razor blade (Fisher Scientific, #12-640) was inserted to remove the olfactory bulbs. The second razor blade was inserted into the third dent. We then removed the subcortical region from this section and separated the left and right hemisphere cortical samples into different Eppendorf tubes. The sampled frontal cortex corresponds to the cortical regions from #23 to #37 sections in the mouse P56 Coronal Atlas from the Allen Brain Atlas released in 2011. We also dissected out hippocampus and cerebellum and put them in separate tubes. The tubes were flash frozen in liquid nitrogen.

We extracted the total RNA of frontal cortex, hippocampus, cerebellum, 293T cells, and iPSC-induced neurons by using the miRNeasy Mini kit (Qiagen, #217004) according to the manufacturer’s protocol. The total RNAs were depleted of genomic DNA by applying DNaseI Amplification grade (Invitrogen, #18068015), and then reverse transcribed used SuperScript III First-Strand Synthesis SuperMix (Invitrogen, #18080400) to amplify cDNA following the manufacturer’s protocol. The qRT-PCR mixture contains 5ul iTaq Universal SYBR Green Supermix, 3ul cDNA template, 0.3ul 5uM primers mix and 1.7ul molecular grade H_2_O in one well of a 384-well plate, each sample was run in replicated reactions in 4 wells. We performed qRT-PCR using the same setup as LoxP copy quantification. Cq values of the 4 replicates from the same sample were averaged, and data of qRT-PCR were calculated ΔΔCq using a Touch Real-Time PCR Detection System and CFX Manager software (CFX384, Bio-Rad Laboratories, Hercules, CA). We used 18S and β-actin as the reference for normalization of expression. We designed the primers (F: 5’ AGCATGGTCCAGATTCCATC and R: 5’ CCCACAAGCTACAGGAGCAG) to amplify 146bp fragments for both mouse Clock and human CLOCK to check generic *CLOCK/Clock* expression in different brain regions of human CLOCK founder mice. To compare the expression of *CLOCK/Clock* in different brain regions of HU, WT, KO, and MS mice, we designed the primers (F: 5’ CACAAAACAGCACCCAGAGT and R: 5’ ATGACTGCCCCACAAGCTAC) to amplify 128bp fragments of human CLOCK exclusively, the primers (F: 5’ AGGAGCTGGGGTCTATGCTT and R: 5’ AGCATCTGACTGTGCAGTGG) to amplify 116bp fragments of mouse Clock exclusively, and the primers (F: 5’ GAGAAGTACAAATGTCTAC and R: 5’ CCATCAAAAATACTACTGTC) to amplify 114bp fragments of generic CLOCK/Clock. We also designed the primers (F: 5’ TGTCTCCTAGGTGCCTGCTT and R: 5’ CTCTCACGGGCTTCTCAGAC) to amplify 115bp fragments of mouse 18S, and the primers (F: 5’ CCAACCGTGAAAAGATGACC and R: 5’ CCATCACAATGCCTGTGGTA) to amplify 120bp fragments mouse β-actin as reference genes in mouse brain tissues. We then designed the primers (F: 5’ TACTGCAGCTGGAAATGTGC and R: 5’ TTGTTGCCACATCATCGTTT) to amplify 184bp fragments of TENM2 (primer1), the primers (F: 5’ GGCTTACCAGGAAACGATGA and R: 5’ CACCAAAGAGAGCGTCCTTC) to amplify 217bp fragments of human TENM2 (primer2), the primers (F: 5’ GTCCTCGCCTACACAAAGGA and R: 5’ GCAGGTGACGTACCGAAAAT) to amplify 186bp fragments of human SORCS2 (primer1), the primers (F: 5’ CTCGGTGGAGATTTTCGGTA and R: 5’ AATGCATACTTCGGCAGCTT) to amplify 212bp fragments of human SORCS2 (primer2), and the primers (F: 5’ GAGGGAGCCTGAGAAACGG and R: 5’ GTCGGGAGTGGGTAATTTGC) to amplify 68bp fragments of human 18S in HEK293T cells and human iPSCs-derived neurons. At last, we designed the primers (F: 5’ TGCCTACGGTTGTCATCTTT and R: 5’ CTCAACTCGCAATATCGCCA) to amplify 303bp fragments of human CLOCK, the primers (F: 5’ AGTAGGTCAGGGACGGAGGT and R: 5’ CAGGATTCCACATTGGAAAAG) to amplify 170bp fragments of human ARNTL, the primers (F: 5’ CTCTGCCTTCCCCATCGTC and R: 5’ TGGAACATGCTGAACTGTGC) to amplify 116bp fragments of human NPAS2, the primers (F: 5’ AACCCCGTATGTGACCAAGA and R: 5’ CCGCGTAGTGAAAATCCTCT) to amplify 138bp fragments of human PER1, the primers (F: 5’ GTCCCAGGTGGAGAGTGGT and R: 5’ AAATTCCGCGTATCCATTCA) to amplify 159bp fragments of human PER2, the primers (F: 5’ ACTTCGAGAAGCAGCTCACC and R: 5’ GCACCGGTAGTAGCAGGAC) to amplify 161bp fragments of human PER3, the primers (F: 5’ CTACATCCTGGACCCCTGGT and R: 5’ CACATCTGCTGGTTGTCCAC) to amplify 150bp fragments of human CRY1, the primers (F: 5’ GAACTCCCGCCTGTTTGTAG and R: 5’ CATCTTCATGATGGCTGCAT) to amplify 144bp fragments of human CRY2, the primers (F: 5’ CCAGTTTGAATGACCGCTCT and R: 5’ AAGCTGCCATTGGAGTTGTC) to amplify 160bp fragments of human NR1D1, the primers (F: 5’ GAGCAGGGGATCTGCTAAACT and R: 5’ GCACGAATGAGAGTTTCCTGC) to amplify 172bp fragments of human NR1D2, and the primers (F: 5’ CCCAGCTGATCTTGCCCTAT and R: 5’ CTCGTTGTTCTTGTACCGCC) to amplify 182bp fragments of human DBP in human iPSCs. All primers were designed by using Primer3 (Untergasser et al., 2012).

### Immunohistochemistry (IHC)

We anesthetized P18 and adult mice by injection of 80-100 mg/kg Euthasol. All mice were transcardially perfused with 4% ice-cold paraformaldehyde (PFA) (Sigma-Aldrich, #P6148), and we kept brains in 4% PFA in 4°C overnight for postfixing. Brains of P0 pups were kept in 4% PFA in 4°C overnight and then transferred to 4°C 30% sucrose (Sigma-Aldrich, S5016) with 0.01% sodium azide (Fisher, #S227I) until sectioning. For sagittal sections, we embedded adult brains in Tissue-Tek CRYO-OCT Compound (Thermo Fisher Scientific, #14-373-65) and cryosectioned onto slides at 20 μm. For coronal sections, we embedded brains in 2% Low Melting Temperature Agarose (UltraPure™, #15517-022) and sectioned with a Compresstome (Precisionary Instruments, #VF-300). The thickness of section depends on the experiments which are specified in corresponding parts. We stored free-floating sections in 1X PBS with 0.01% sodium azide. We used 1X TBS, which is diluted from 10X TBS (30.5g Tris base, 45g NaCl (Sigma-Aldrich, S3014), ddH_2_O to 500ml final volume, pH 7.6), for all washes. We diluted primary antibodies in 3% normal donkey serum (Jackson ImmunoResearch, #017-000-121) and 1% bovine serum albumin (BSA) (Sigma-Aldrich, #A9647) in 0.4% Triton X-100 (Thermo Fisher Scientific, #BP151-100) in TBS (TBS-T). Secondary antibodies were diluted in 1% BSA in TBS-T.

#### Lamination

We coronally sectioned adult mouse brains at 40 μm. Sections from frontal cortex, which are from #40 to #52 sections in the mouse P56 Coronal Atlas released as the Allen Brain Atlas in 2011, were used for lamination analysis. We put sections in citrate buffer (10 mM Sodium Citrate Tribasic Dihydrate (Sigma, #C8532), 0.05% Tween-20 (Fisher, #BP337), pH 6) for 10 min at 95°C for antigen retrieval. Aldehydes were quenched with 0.3 M Glycine (Sigma-Aldrich, #G8898) in TBS-T for 1h at RT. We then incubated sections in primary antibody solution overnight at 4°C. Secondary antibody incubations were performed for 1h at RT. We mounted sections onto slides (Fisherbrand™, #12-550-15) and waited until dry. #1 Coverslips (Fisherbrand™, #12-545-MP) were mounted using ProLong Gold Antifade Mountant with DAPI (Thermo Fisher Scientific, #P-36931). We used rabbit anti-CUX1 (1:200; Proteintech, #11733-1-AP) to label layers 2-4, mouse anti-FOXP1 (1:500; Abcam, #ab32010) to label layers 3-5, goat anti-FOXP2 (1:1000; Santa Cruz Biotechnology, #sc-21069) to label layer 6. We used species-specific secondary antibodies produced in donkey and conjugated to Alexa Fluor 488, Alexa Fluor 555, or Alexa Fluor 647 (1:2000). 20X images with fixed 13 z stacks and tile scans to capture the hemisphere of the coronal section were collected using a Zeiss confocal laser scanning microscope (LSM880) at the UT Southwestern Neuroscience Microscopy Facility. Image analysis was performed in FIJI (Schindelin et al., 2012) with self-written Macro codes. For each section, we did three measurements spanning the width of the primary somatosensory cortex for all layers, layers 2-4, and layer 6 respectively). We also measured the area of the hemisphere in which we measured layers. We divided the length of all layers to area for relative cortical thickness, and we divided the length of particular layers to cortical thickness for relative thickness of layers. All three measurements were averaged to represent the section.

#### Cell density

We coronally sectioned adult mouse brains from frontal cortex, which correspond to images from #27 to #36 in the mouse P56 Coronal Atlas released as the Allen Brain Atlas in 2011, into 40 μm slices for cell density analysis. We put sections in 3% H_2_O_2_ (dilute in ddH_2_O from 30% Hydrogen Peroxide Solution; Sigma-Aldrich, #H1009) for 1h at RT to block intrinsic peroxidase activity. Aldehydes were quenched with 0.3 M glycine in TBS-T for 1 hour at RT. We then incubated sections in blocking solution (3% normal donkey serum and 1% BSA in TBS-T) for 1h at RT. Sections were transferred to primary antibody solution for 48 hours at 4°C. Secondary antibody incubations were performed overnight at 4°C. We then incubated sections in 300 nM DAPI solution in 1X PBS for 5 min at RT. Sections were mounted onto slides, and #1 coverslips were mounted using ProLong Gold Antifade Mountant (Thermo Fisher Scientific, #P-36970). We used rabbit anti-CLOCK (1:200; Santa Cruz Biotechnology, #sc-25361) to label CLOCK+ cells, mouse anti-NEUN (1:1000; Millipore, #MAB377) to stain neurons, and goat anti-OLIG2 (1:500; R&D Systems, #AF2418) to stain oligodendrocytes (including both mature oligodendrocyte and OPC). We used species-matched secondary antibodies produced in donkey and conjugated to Alexa Fluor 488, Alexa Fluor 555, or Alexa Fluor 647 (1:2000). A 20X objective lens with fixed 13 z stacks and tile scans was used to capture the hemisphere of the coronal section using a Zeiss LSM 880 confocal microscope at the UT Southwestern Neuroscience Microscopy Facility. We performed cell count analysis in a 1513 x 1411 pixels (623 x 587 um) field of orbitofrontal cortex (**Fig. 3A**) by using FIJI (Schindelin et al., 2012) with self-written Macro codes. We counted DAPI_ NEUN double positive cells as neurons and DAPI_OLIG2 double positive cells as oligodendrocytes. At last, all cells, neurons, and oligodendrocytes were colocalized with CLOCK + cells for CLOCK+ neurons and CLOCK+ oligodendrocytes respectively to calculate CLOCK + fractions.

#### Fluorescent intensity

P07 and P56 coronal sections of brain from similar regions as cell density analysis were used for TENM2 and SORCS2 imaging respectively. The IHC and imaging steps are similar to those in lamination. We used sheep anti-TENM2 (1:100, Invitrogen, #PA5-47638) and sheep anti-SORCS2 (1:500; R&D Systems, #AF4238) to stain the two proteins, and mouse anti-CAMKII (1:100; Abcam, #ab22609) for excitatory neurons in P07 sections. Image capture used the same settings as described above. The fluorescent intensity was quantified from excitatory neurons for TENM2, and it was quantified from a 511 x 441 pixels (212 x 183 um) field of frontal cortex for SORCS2 and normalized to the number of cells.

#### In vivo lateral ventricle injection, dendrite and spine morphology

We performed *in vivo* lateral ventricle injection with recombinant adeno-associated virus (rAAV/hSyn-mcherry, Virus Vector Core, University of North Carolina at Chapel Hill) that express mCherry to visualize dendrites and spines of neurons as previously described (Kim et al., 2014). We prepared the needles for virus injection by pulling glass pipettes (World Precision Instruments, # 1B100-6) on a puller (Narishige International, #PC-10). We connected the needle to the micro syringe of Standard Infusion Only Pump 11 Elite Syringe Pumps (Harvard Apparatus, #70-4500) and fixed the needle on a stereotaxic (DAVID KOPF INSTRUMENTS, Model 940). P0-P1 mice were wrapped with a disposable rubber glove and buried in ice for 5-10 min for cryo-anesthetization. We placed the mouse on an ice-cold platform and fixed the head. We adjusted the needle to the intersection dot of transverse and sagittal suture lines and set this dot as the initial position (i.e. x=0, y=0, z=0). We then injected the rAAV in the left lateral ventricle (x=0.8mm left, y=1.5mm rostral, z=1.5mm deep) with infusion rate 3.33nl/second for a total of 50nl. The injected mouse was rubbed with its home cage bedding material gently and put back to its home cage which was placed on a heating pad. Mice were transcardially perfused as described above and the brains collected on P18 and P56.

We coronally sectioned P18 and adult mice at 100 μm. We used sections from frontal cortex, which correspond to images from #37 to #47 in the mouse P56 Coronal Atlas released as the Allen Brain Atlas in 2011, for dendrite and spine analysis. The IHC procedure was the same as the above description for cell density. Instead of 1# coverslip, we used #1.5 coverslip (Globe Scientific, #1415-15) for use with a high magnitude oil object. We used mouse anti-NEUN to label neurons, and goat anti-OLIG2 to stain oligodendrocytes, chicken anti-GFAP (1:500; Abcam, #ab4674) to label astrocytes, rabbit anti-CUX1 to label layers 2-4, mouse anti-CAMKII to stain excitatory neurons, and rabbit anti-DsRed (1:500; Clontech, # 632496) and goat anti-tdTomato (1:500; LifeSpan BioSciences, #LS-C340696) to enhance the mCherry signal. We used species-specific secondary antibodies produced in donkey and conjugated to Alexa Fluor 405, Alexa Fluor 488, Alexa Fluor 555, or Alexa Fluor 647 (1:100 for Alexa Fluor 405, 1:2000 for the others). A 63X objective lens with fixed 13 z stacks and tile scans was used to capture the entire arborization of a given neuron using the airyscan function of the Zeiss LSM 880 confocal microscope at the UT Southwestern Neuroscience Microscopy Facility. We used self-written Macro codes to apply Simple Neurite Tracer (Arshadi et al., 2021; Longair et al., 2011) and Sholl analysis (Ferreira et al., 2014; Sholl, 1953) plugins in FIJI to analyze the complexity of dendritic arborization (i.e. length of dendrites, number of branches, number of terminal, and number of intersections). Spine analysis was conducted by using self-written Macro codes and SpineJ (Levet et al., 2020) plugin in FIJI. We measured the morphological traits of spines (i.e. spine length, head area, head width, and neck width) classified spines into three categories (Berry and Nedivi, 2017; Harris et al., 1992; Peters and Kaiserman-Abramof, 1970): stubby (head width ≤ neck width), thin (2x neck width > head width > neck width), and mushroom (head width ≥ 2x neck width), and quantified the density for each type of spine.

### Immunoprecipitation (IP)

We performed IPs as we did previously (Usui et al., 2017b). We homogenized mouse cortex in 1X RIPA buffer (250 mM Tris-HCl pH 7.4, 750 mM NaCl, 5 mM EDTA, 5mM Na_3_VO_4_, 2.5% sodium deoxycholate, 0.5% SDS, 5% Igepal) with 10ul/ml of 100mM PMSF, 25ul/ml of 200mM Na3VO4, and 10ul/ml Protease Inhibitor Cocktail (Sigma-Aldrich, #P8340) by using TissueLyser LT (QIAGEN, #85600). We then sonicated the homogenized tissue using a Bioruptor (Diagenode, #UCD-200) in the following procedure: 10 seconds pulses of 200 W with an interval of 20 seconds for 5 min twice. Samples were centrifuged at 12,000 rpm for 15 min at 4°C. The supernatant of each sample was transferred to a new tube and incubated with 100ul Dynabeads Protein G (Invitrogen, #10004D) and rabbit anti-CLOCK (1:100; MilliporeSigma, #AB2203). At last, we used 40ul 1X SDS (1mM EDTA, 0.5% SDS, 20mM Tris-Cl) to elute proteins from Dynabeads for western blot.

### Western blot

In addition to using eluates from IP, we also collected protein from cultured cells. We washed cells with 1X PBS, collected in lysis buffer (0.5% Igepal, 50mM Tris pH 8, 10% glycerol, 0.1mM EDTA, 250mM NaCl, 50mM NaF, 0.7mM PMSF, 100X diluted Protease Inhibitor, 10mM DTT, 100uM Na3VO4). and centrifuged at 12,000 rpm for 10 min at 4°C. We performed Bradford assays (Bio-Rad) on the supernatants to quantify the protein concentration. We ran 50ug protein of each sample on 10% SDS-Page gels, and then transferred proteins to PVDF membrane (Bio-Rad, 162-0177). PVDF membrane was incubated in blocking solution (TBS with 1% Skim milk and 0.1% Tween-20) for 30min at RT, and then we added rabbit anti-CLOCK (1:200; Abcam, #ab3517), mouse anti-FLAG (1:400; Sigma, #F1804), rabbit anti-CLOCK (1:200; in house antibody raised in rabbit with epitope 377-556 a.a), and mouse anti-GAPDH (1:5000; Millipore, #MAB374) overnight at 4°C. PVDF membrane was washed in TBS with 0.1% Tween-20, and then incubated with corresponding secondary antibodies similar to IHC in blocking solution for 1h at RT, washed in TBS with 0.1% Tween-20 and imaged on an Odyssey infrared imaging system (LI-COR Biosciences).

### Wheel-running rhythm

Mice (N = 6-10/genotype) at 8-12 weeks of age were housed in wheel-running cages (Siepka and Takahashi, 2005) to monitor their locomotor activity rhythm. We initially exposed mice to a 12:12 hours of light and darkness cycle (LD) for two weeks followed by 4 weeks of constant darkness (DD), and then we returned to a LD cycle for two weeks followed by 4 weeks of constant light (LL). We recorded wheel-running activity and analyzed the free-running period of DD and LL by using χ^2^ periodogram analysis in ClockLab (Actimetrics, Wilmette, IL).

### Behavioral tests

All tested mice were handled 5 min/day by the experimenter for two weeks prior to experiments to alleviate their handling stress in experiments (Ueno et al., 2020). We tested young adult mice (8-12 weeks of age) in the behavioral test battery. Mice underwent tests in the following order: open field test, novel object recognition test, social memory and aggressiveness test, set-shifting test, olfactory test, neurological reflex test, and tail suspension test. The order of the behavioral tests was organized to go from least to most stressful. We also applied two weeks of inter-test intervals to mice between two adjacent tests to minimize the effects from the previous test (McIlwain et al., 2001). Fear conditioning was conducted using a different cohort of animals.

#### Open field test

The open field was conducted in the same apparatus and with the same procedure as previously described (Xu et al., 2021). We individually released mice into the center of the arena and allowed them to explore for 1 hour. The first 30 min was video-recorded and analyzed using Limelight4 (Actimetrics, Wilmette, IL). We quantified the total distance moved as a measure of activity. We divided the whole square arena into 25 (5X5) small square areas. We defined the 16 areas on the edge as the outside areas, while the 9 areas in the middle were defined as the center areas. We quantified the duration in the outside areas and the center of areas as a measure of anxiety.

#### Novel object recognition test

We did the novel object recognition test in the same arena as open field test. 24 hours after the open field test, we provided two identical glass vials in the arena as a training session. We individually released mice and allowed 30 min to investigate the objects. We video-recorded the first 15 min for analysis. The testing session happened 24 hours later; we replaced one of the glass vials with a nail polish bottle, and then individually released the mouse to investigate the objects for 15 min. Performance was recorded and analyzed using Limelight 4. A recognition index of the ratio of nose investigation duration between the novel and familiar object in the testing session was calculated (Antunes and Biala, 2012).

#### Social memory and aggressiveness test

We used a five trials social memory test to mimic the type of short but frequent interactions that occur in a colony setting (Winslow and Camacho, 1995). Juvenile (4-6 week old) sex-matched mice were used as social stimuli. We placed the test mouse into a new cage for a 30 min habituation period. A stimulus mouse was introduced into the cage for 1min. After a 10min delay, this was repeated for a total of five 1min exposures, with the same stimulus mouse in the first 4 exposures, and a novel stimulus mouse in the fifth exposure. We video-recorded and manually quantified the frequency and duration of social interaction (i.e. body investigation and attack).

#### Set-shifting test

Mice were calorie restricted. Food was only provided for 4 hours in the first 24 hours. For the second 24 hours, food was provided for 2 hours. For the remainder of the testing period, food was only provided for 1 hour per day. The mice maintained 80-85% of their ad libitum feeding body weight during the test. In the acclimation session before the real test, mice were trained to dig bedding medium in a bowl to obtain a ∼10mg piece of peanut butter chip. This session consisted of two days of training. On the first day, we trained mice to recognize the peanut butter chip as food, and successfully obtain 23 pairs of chips from two empty feeding bowls. On the second day, mice started with obtaining 13 pairs of chips from two empty feeding bowls, followed by obtaining 13 pairs of chips from bowls with kay-kob (Kaytee, #1851118) gradually added, and at last obtaining 23 pairs of chips that are fully covered by kay-kob. The set shifting test consisted of four consecutive tests in the following order: discrimination, reversal, intra-dimensional shift (IDS), and extra-dimensional shift tests (EDS). We did all tests in home cages. Two sensory modalities (visual and olfactory) were used. For each test, we paired 2 odors with 2 mediums (4 possible combinations) to the buried chip. A food reward was only associated with a particular stimulus in one modality. The mouse had to not only discriminate reward and non-rewarded modalities, but also discriminate and associate the correct stimulus in the correct modality with food reward. In discrimination, reversal, and intra-dimensional shift tests, the reward modality was consistent, while the reward modality was switched in the extra-dimensional shift test. Except for the discrimination and reversal tests, all stimuli were changed from test to test. Discrimination and reversal tests shared the same stimuli, but their association to reward were opposite. **Table S2** shows an example of combinations of stimuli across tests, and we counterbalanced all combinations of reward modalities and stimuli. A mouse was trained for up to 50 trials in each test. Criterion to learn was considered as 8 consecutive correct trials. A mouse that reached the criterion or 50 trails was forwarded to the next test. We quantified their performance by using the number of trials to criterion (Heisler et al., 2015).

#### Olfactory test

Since the set shifting test relies on olfactory discrimination, we ran an olfactory test to see if genetic modification of Clock/CLOCK affects olfactory discrimination. Mice were individually kept in a new cage for 30 minutes with a dry cotton swab attached to the center of the cage ceiling. We then tested mice in nine consecutive trials in the following order: three water dipped cotton swab trials, three vanilla cotton swab trials, and three lemon cotton swab trials. Each trial lasted for 2 minutes with 1 minute inter-trial intervals. We quantified the duration of nose investigation on the cotton swab (Yang and Crawley, 2001).

#### Neurological reflex test

We conducted eye blink reflex, ear twitch reflex, and whisker orienting reflex with a cotton swab to examine any sensory-motor deficits. We quantified these data in a binary manner as blink or not, twitch or not, and orienting to touched whisker or not (Heyser, 2003).

#### Tail suspension test

We first attached a 5cm Tygon tubing to the base of the mouse tail to prevent climbing onto its own tail, and then we attached a 20cm piece of yellow lab tape to the tail tip of each mouse. Mice were then placed at a height of 30cm height from the surface allowing them to hang from their tails. We tested 4 mice per batch. Adjacent individuals were about 20cm apart from each other, and there was no barrier between them. We video-recorded their performance for 6 min, and then manually quantified duration of immobility as previously described (Can et al., 2012).

#### Fear conditioning test

We carried out fear conditioning testing as previously described (Xu et al., 2021). We trained young adult mice in the fear-conditioning chambers. Mice were allowed to freely explore the chamber for 3 min. Then we applied a tone (2.8 KHz, 85 dB) for 30 sec, and a 2 second 0.85 mA foot shock overlapped with the last 2 sec of the tone. We applied the paired tone and foot shock stimuli for 4 times with 1 min inter-trial intervals. After 24 hours, we re-exposed the mice to the context (the same chamber in training) for 5 min, followed by the sound cue memory test (in a different chamber with different context and odor cues in another room but with the same tone) for 5 min. We used Ammeter (Med Associates, # ENV-421) to measure the shock intensity from each pair of neighboring wire bars on the grids, and sound was calibrated with a sound meter. The metal grid floors (for training) and plastic floors (novel context) were washed with water, sprayed with 70% ethanol and dried before each trial to avoid odor cues. All sessions were video-recorded and freezing of mice was scored by FreezeFrame3 (Actimetrics, Wilmette, IL).

### Body and brain mass

We measured the body mass of living mice at P7, P21, and young adult age (8 weeks to 12 weeks). From euthanized mice, we removed the brains, which included medulla and olfactory bulb, from P7 and P21 mice, and weighed immediately. All measurements were conducted by using a tabletop scale (Denver Instruments, #MXX-412).

### Single-nuclei RNA-seq

#### Tissue collection, nuclei isolation, and library preparation

Our nuclei isolation procedure was modified from our previous work (Ayhan et al., 2021). The frozen tissue was transferred to a Dounce homogenizer (DWK Life Sciences, # 885300-0002) with 2ml of ice-cold Nuclei EZ lysis buffer (Sigma, #NUC-101). We then inserted pestle A for 23 strokes followed by pestle B for 23 strokes on ice. The homogenized sample was transferred to a 15ml conical tube. We added 2ml of ice-cold Nuclei EZ lysis buffer and incubated on ice for 5min. Nuclei were collected by centrifuging at 500 g for 5min at 4°C. We discarded the supernatant and added 4ml of ice-cold Nuclei EZ lysis buffer to resuspend nuclei. We then repeated the incubation and centrifuge steps and resuspended the nuclei in 200ul of nuclei suspension buffer (NBS) (1x PBS, 1% ultrapure BSA (Thermo Fisher Scientific, #AM2618), and 0.2 U/ul RNAse inhibitor (Thermo Fisher Scientific, #AM2694)), and finally filtered the nuclei suspension twice through a Flowmi Cell Strainer (Bel-Art, H13680-0040).

We mixed 10ul of nuclei suspension with 10ul of 0.4% Trypan Blue (Gibco, #15250061) and loaded onto a hemocytometer (INCYTO, #DHC-N01) to determine the nuclei number concentration. We then diluted each sample to 1000 nuclei/ul in NBS. 10,000 nuclei/sample was used to generate snRNA-Seq libraries using 10X Genomics Single Cell 3’ Reagent Kits v3 (10x Genomics, #1000153) following the manufacturer’s protocol (Zheng et al., 2017). We processed 24 mice (2 males and 2 females per genotype per age) for snRNA-Seq. Libraries were sent to the McDermott Sequencing Core at UT Southwestern for sequencing using Illumina NovaSeq 6000.

#### Data analysis

##### Data processing

Raw BCL data for sequenced libraries was downloaded from the McDermott Sequencing Core at UT Southwestern. Self-written codes were used for data processing. FASTQ files were extracted using CellRanger *mkfastq* (v3.0.2) (Zheng et al., 2017). FASTQ files were next processed through UMI-Tools (v0.5.4) *whitelist* with ‘--set-cell-number=50000’ to identify the top 50,000 cell barcodes per sample (Smith et al., 2017). Reads corresponding to identified cell barcodes were then extracted using *extract* from UMI-Tools suite. Extracted FASTQ reads were then aligned to the reference mouse genome (MM10 GRCm38p6) and reference mouse Gencode annotation (vM17) using STAR (v2.5.2b) (Dobin et al., 2012). Samples corresponding to genotype HU were additionally aligned to a custom reference assembly containing only human *CLOCK* that was extracted from the reference human genome (HG38 GRCh38p12) and human Gencode annotation (v28) using STAR (v2.5.2b) (Dobin et al., 2012). Uniquely aligned BAM reads were further assigned to genes using featureCounts from Subread package (v1.6.2) using reference Gencode annotations (Liao et al., 2013). BAM files with alignment and gene assignment information were then sorted and indexed using Samtools (v1.6) (Li et al., 2009). Sorted and indexed BAM files were further used to generate a raw counts matrix (per gene per cell barocde) using *count* from UMI-Tools suite. Raw count matrices per sample were re-organized using R SeuratDisk package (https://github.com/mojaveazure/seurat-disk) in order to match the input requirements for CellBender. CellBender was then run on each sample to remove noise from ambient RNA (Fleming et al., bioarxiv). Potentials doublets were filtered out using DoubletFinder R package (McGinnis et al., 2019). Cleaned and filtered count matrices were then used to generate Seurat objects and QC plots per age group per genotype. Before generating Seurat objects for samples that belong to genotype HU, data were updated to retain gene expression information for both the endogenous copy and humanized copy of *Clock/CLOCK*.

##### Clustering analysis

R package Seurat (v4.0.6) (Hao et al., 2021) and self-written scripts were used for downstream analyses. After filtering, the raw UMI counts were normalized with a scaled factor of 10,000 and regressed to covariates such as the number of UMIs per cell, percent mitochondrial content per cell, batch, and sex as described in the Seurat analysis pipeline (Hao et al., 2021). Expression of the top 2000 variable genes was used to calculated principal components (PCs). Based on the result of JackStraw analysis, 30 and 28 PCs, which include the majority variation for P07 and P56 samples respectively, were used to identify cell clusters by using the original Louvain algorithm as described in the Seurat analysis pipeline (Hao et al., 2021). Dimension reduction and cluster visualization was conducted using Uniform Manifold Approximation and Projection (UMAP) (Becht et al., 2018; Kulkarni et al., 2019). 5 and 4 clusters of P07 and P56 datasets were removed due to their low number of UMIs (<10).

##### Cell type annotation

Gene enrichment per cluster was determined by Log_2_ fold change ≥ 0.3 and FDR adjusted p value ≤ 0.05 for a given gene/cell compared to the rest of the cells in a cluster. We then applied self-written code to do pairwise Fisher Exact Tests between cell types of this study and previously published brain single-cell datasets (Hrvatin et al., 2018; Yao et al., 2021) (**Fig. S6**). We assigned the cell types to clusters based on the statistical results and location of cluster on the UMAP. One cluster of P07 did not show enrichment with either published dataset, but it showed enrichment of striatal neurons using Cell-type Specific Expression Analysis tool (http://genetics.wustl.edu/jdlab/csea-tool-2/) (Dougherty et al., 2010). Hence, we removed this cluster.

##### Differentially expressed gene (DEG) analysis

DEG analyses were conducted pairwise (i.e. between HU and WT, and between KO and WT) within either P07 or P56. We subsetted cells of major cell types (i.e. astrocyte, endothelium, excitatory neuron, inhibitory neuron, mature oligodendrocyte, oligodendrocyte precursor cell, and vascular leptomeningeal cell) as well as the subtypes of excitatory neurons (i.e. EX2-3_IT, EX4-5_IT, EX5_IT, EX6_IT, EX5_ET, EX5-6_NP, EX6_CT, and EX6b). We used MAST (v1.8.2) (Finak et al., 2015) with linear mixed model for a zero-inflated regression analysis (Velmeshev et al., 2019) to calculate an FDR adjusted p value for each gene. The linear mixed model tested genotype as fixed factor, sample as random factor, and other parameters as covariates (such as batch, sex and so on). We also applied the Seurat analysis pipeline to calculate log_2_ fold change and percentage of expressed cells (Hao et al., 2021). Genes expressed in at least 30% of cells with |log_2_ fold change| ≥ 0.1 and FDR adjusted p value ≤ 0.05 were determined as DEGs.

##### Hypergeometric overlap test

A hypergeometric test was used to assess the significance of overlaps of DEGs across cell types, between different ages, and between human CLOCK and mouse Clock. It was also used to test overlap between DEGs of HU-WT comparison in EX2-3_IT and human-specific open/closed chromatin (Emre et al., unpublished). FDR adjusted p values were calculated based on the number of overlaps that occurred in a set of pairwise overlaps (e.g. **Fig. 4E-H** and **Fig. S7**).

##### Gene ontology (GO) enrichment analysis

ToppGene Suite (Chen et al., 2009) was used for GO analysis to determine the functions that were enriched by the DEGs. We used the expressed genes in P07 and P56 as the background input to ToppGene Suite. We did not apply a p value cutoff for output. The outputs were downloaded and processed using self-written code for further analysis and plot generation. Significant enrichment was set using a Benjamini-Hochberg FDR adjusted p value < 0.05.

##### Motif analysis in enhancer regions

Enhancer regions of TENM2/Tenm2 and SORCS2/Sorcs2 were obtained from Search Candidate cis-Regulatory Elements by ENCODE (SCREEN) (Consortium et al., 2020), and we further downloaded the sequences of these enhancer regions from UCSC Genome Browser (Kent et al., 2002). We then scanned the sequences of the enhancer regions for CLOCK:BMAL1 motif by using Mulan (Ovcharenko et al., 2005).

### Electrophysiology

#### Brain slices and recordings

P21-27 mice of both sexes were used for the electrophysiological recordings. We anesthetized mice with a mixture of ketamine (125 mg/kg) and xylazine (25 mg/kg), and then quickly decapitated to dissect out the whole brain. The brain was glued on the section panel of a Vibratome (Leica, #VT1200S) and immersed in ice-cold dissection buffer (110mM Choline-Cl, 2.5mM KCl, 1.25mM NaH_2_PO_4_, 25mM NaHCO_3_, 25mM Dextrose, 3mM Ascorbic Acid, 3.1mM Sodium Pyruvate, 7mM MgCl_2_, 0.5mM CaCl_2_.2H_2_O, and 0.2mM Kyneuric Acid, pH 7.4, 285 mOsm osmolality). The brain was sectioned into 300μm thickness slices (corresponding to the images from #39 to #47 in the mouse P56 Coronal Atlas released as the Allen Brain Atlas in 2011). The slices were immediately transferred to nominal Artificial Cerebrospinal Fluid (ACSF) (125mM NaCl, 3mM KCl, 1.25mM NaH_2_PO_4_, 25mM NaHCO_3_, 10mM Dextrose, 2mM MgCl_2_, and 2mM CaCl_2_.2H_2_O, pH 7.4, 295 mOsm osmolality, bubbled with 5% CO_2_ and 95% O_2_) at 35°C for 30 min, and then kept at RT for at least 30 min before recordings. Excitatory neurons from layers 2-4 were patched from the somatosensory cortex region of these slices using glass pipettes of 4-6 MΩ resistance, filled with K-Gluconate containing internal pipette solution (130mM K-Gluconate, 6mM KCl, 3mM NaCl, 10mM HEPES, 0.2mM EGTA, 4mM ATPMg, 0.4mM GTP-Tris, and 14mM phosphocreatine-Tris, pH 7.25, 305 osmolality). Neurons with a stable resting membrane potential and series resistance less than 20 MΩ were used for analysis.

#### Current steps for the excitability measurements

To measure the intrinsic excitability of the recorded neurons, we injected incremental current steps of 50pA amplitude and 500 ms duration in the current clamp at resting membrane potential. The number of action potentials at each current step were counted and plotted in the firing versus injected current curve. Also, the resting membrane potential of each neuron was determined and plotted.

#### Input resistance and normalized conductance

Input resistance of the neuron was measured in voltage clamp by applying single 500 ms duration, −10 mV voltage step at −85, −65 and −55 holding potentials. The recorded data from the voltage step was used to calculate the conductance and capacitance of each neuron was also calculated. Finally normalized conductance was determined by dividing the conductance by the capacitance.

#### Spontaneous and evoked EPSCs

Spontaneous EPSCs were recorded from layer 2-4 excitatory neurons in voltage clamp at −70 mV for 60 seconds while circulating oxygenated normal ACSF in the bath solution. Amplitude and frequency of the spontaneous EPSCs were analyzed using MiniAnalysis (Synaptosoft, Decatur, CA). Events were first detected automatically by the software after setting up cutoffs for the area (30) and height (10) of an event.

AMPA-receptor mediated evoked EPSCs were recorded from the layer 2-4 excitatory neurons in response to the extracellular electrical stimulation of horizontal intercortical afferents. We placed a cluster bipolar stimulating electrode (FHC, Bowdoin, ME) parallel to the recorded layer 2-4 excitatory neurons and applied biphasic pulse of 100 us interval to evoke an EPSC. ET (threshold intensity) is the minimum current intensity that evoked an EPSC, whereas AT is the amplitude of the EPSC response at ET. We then applied stimulus greater than the ET (2-fold, 3-fold, 4-fold and 5-fold of the ET) and measured the amplitude of the evoked EPSC. Finally fold change increases in the amplitude of the EPSC (with respect to AT) were calculated and plotted with the fold change increase in the stimulus intensity.

### CRISPR-Cas9 generation of CLOCK KO iPSC line

The CLOCK KO cell lines were generated by targeting the exon 7 of CLOCK with a pair of sgRNAs following the published guideline (Ran et al., 2013). The following set of sgRNAs were used: 5’-CACCGCTAGTGAAATTCGACAGGAC-3’ and 5’-AAACGTCCTGTCGAATTTCACTAGC −3’. To maximize the efficiency and mitigate potential off-target effects of gene editing, sgRNA candidates were designed and analyzed using CHOPCHOP v3 (Labun et al., 2019) and CRISPOR (Concordet and Haeussler, 2018) respectively. SgRNA sequences with fewest off-target sites in the human genome were selected for use. Each sgRNA was inserted into pSpCas9(BB)-2A-GFP (PX458) (Addgene, #48138) (Ran et al., 2013). Cells were incubated with ROCK inhibitor for 1-3 hours and then electroporated with the pair of sgRNAs. 48-72 hours post electroporation, cells were subjected to fluorescence-activated cell sorting (FACS) for GFP+ signals and re-plated as single clones for the subsequent 10-12 days. To verify deletion of CLOCK, positive clones from both strategies were confirmed via Sanger sequencing, qPCR, and western blot (**Fig. S14A-C**).

### Generation of iPSC-induced neuron through Neurogenin-2 (NGN2) fast differentiation

We differentiated iPSCs into induced neurons following a protocol modified from the original NGN2 fast differentiation experiment (**Fig. S14D**) (Zhang et al., 2013).

#### Generation of transduced iPSC line

We obtained plasmids of reverse tetracycline-controlled transactivator (rtTA) (FUdeltaGW-rtTA; Addgene, #19780) and NGN2 (pTet-O-Ngn2-puro; Addgene, #52047). We amplified and harvested these plasmids by using EndoFree Plasmid Maxi Kit (Qiagen, #12362), and sent them to the Neuroconnectivity Core at Baylor College of Medicine Intellectual and Developmental Disabilities Research Center for lentivirus packing. We dissociated iPSCs to single cells using Accutase and plated them into 12-well plates at 3.8 x 10^5^ per well in mTeSR^TM^1 medium and ROCK inhibitor. 1 day later, lentiviruses rtTA (titre: 3.108 x 10^10^) and NGN2 (titre: 1.578 x 10^10^) were added in medium with 1:2500 and 1:10000 dilution respectively to transduce iPSCs. The transduced iPSCs were expanded for differentiation and frozen down for future studies following the Technical Manual.

#### NGN2 differentiation

We dissociated transduced iPSCs in Accutase and plated 1.5 x 10^4^ /well and 1 x 10^5^ /well into 24-well and 6-well plates respectively. We placed an acid cleaned glass coverslip (Carolina Biological Supply, #633029) into each well of a 24-well plate. Cells in 24-well plates were used for morphological analysis, while cells in 6-well plates were collected for qRT-PCR. All wells were coated with Matrigel. The confluency was monitored every 12 hours. Once the confluency reached about 50%, we started Day 1 and replaced stem cell medium to differentiation medium KSR (415ml KnockOut DMEM (Gibco^TM^, #10829018), 75ml Knockout Replacement Serum (Invitrogen, #10828-028), 5ml MEM NEAA (Invitrogen, #11140-050), 5ml Glutamax (Gibco^TM^, #35050), and 0.5ml beta-mercaptoethanol (Invitrogen, #21985-023) with 4ug/ml Doxycycline (Sigma-Aldrich, #D9891), 10uM SB431542 (Cellagen Technology, #C7243-5), 100nM LDN-193189 (Selleck Chemicals, #S2618), and 2nM XAV939 (Tocris Bioscience, #3748)). The iPSCs started to differentiate while many cells continued to proliferate (**Fig. S14D**). 24 hours later (Day 2), we changed the medium to a mixture of half KSR and half N2B (500ml DMEM/F-12 (Life Technologies, #11330057), 5ml Glutamax, and 7.5ml 20% Dextrose (Sigma, #G6152) with 1:100 N-2 Supplement (Thermo Fisher Scientific, #17502-048), 4ug/ml Doxycycline, and 0.7ug/ml Puromycin (Sigma-Aldrich, #P8833)); cells differentiated into neural progenitor-like cells and grew until about 90% confluency (**Fig. S14D**). On Day 3, about 70% of cells died due to lack of successfull transduction with rtTA and NGN2 (**Fig. S14D**). Medium was changed to N2B. On Day 4, cells continued die until about 10% of newborn neurons survived (**Fig. S14D**). We changed to NBM medium (485ml NBM (Gibco, #21103), 5ml Glutamax, 2.5ml MEM NEAA, and 7.5ml 20% Dextrose with 4ug/ml Doxycycline, 1:50 B27 (Thermo Fisher Scientific, #17504-044), 10ng/ml GDNF (PeproTech, #450-10), 10ng/ml BDNF (PeproTech, #450-02), and 10ng/ml NT3 (Fisher Scientific, #267N3005CF)), and added 10uM Ara-C (Sigma-Aldrich, #C1768) to eliminate progenitors. On Day 5, medium was changed, and 20,000 astrocytes were added to each well in 24-well plate. After that, cell cultures were divided into three groups. In group 1, cells were kept in NBM until Day 28. The other cultures were CRISPR transfected on Day 6, cells in group 2 and 3 were kept in NBM until Day 21 and Day 35 respectively. After changing of NBM medium on Day 6 for group 1 and Day 7 for group 2,3, NBM was half changed every other day, and 2.5% FBS was add to NBM once per week.

### CRISPR-Cas9 overexpression of TENM2 and SORCS2

We performed CRISPR-Cas9 to overexpress TENM2 and SORCS2 similar to a previous study (Matharu et al., 2019). We designed pairs of forward and reverse sgRNAs by using the above mentioned tools (Concordet and Haeussler, 2018; Labun et al., 2019). F: 5’ ACCGTTTAATTGCCTAAGCGGTCT and R: 5’ AACAGACCGCTTAGGCAATTAAAC target the transcription start site (TSS) of TENM2, F: 5’ ACCGGTTTGCTTTATATGCTCATT and R: 5’ AACAATGAGCATATAAAGCAAACC target the site 33bp ahead of TSS of TENM2, F: 5’ ACCGAGATTGTGCAGGGCTGACCA and R: 5’ AACTGGTCAGCCCTGCACAATCTC target the site 137bp ahead of TSS of TENM2, F: 5’ ACCGAGTGACAAGTGCCAAAACTC and R: 5’ AACGAGTTTTGGCACTTGTCACTC target Exon 2 of TENM2, F: 5’ ACCGCTGGCGACCATGGCGCACCG and R: 5’ AACCGGTGCGCCATGGTCGCCAGC target TSS of SORCS2, F: 5’ ACCGCGGTGCGCGCTCGGCGTCGC and R: 5’ AACGCGACGCCGAGCGCGCACCGC target the site 60bp ahead of TSS of SORCS2, and F: 5’ ACCGGACCGGAGGAACCACCGAGC and R: 5’ AACGCTCGGTGGTTCCTCCGGTCC target the site 168bp ahead of TSS of SORCS2. Each pair of sgRNAs was cloned into pAAV-U6-sgRNA-CMV-GFP (Addgene, #85451) by using the SapI restriction enzyme (BioLabs, #R0569). To confirm the successful insertion of sgRNAs into the plasmids, we sequenced them in the Sanger Sequencing Core at McDermott Center of UTSW by using the forward primer: 5’ ACTATCATATGCTTACCGTAAC.

We transfected HEK293T cells (at about 70% confluency) and iPSC-induced neurons on Day 6 (**Fig. S14D**) by following the Lipofectamine^®^ LTX DNA Transfection Reagents Protocol. We diluted 3ul of Lipofectamine^®^ LTX Reagent (Invitrogen, #15338030) in every 50ul Opti-MEM™ I Reduced Serum Medium (Gibco, #31985088). Vectors of pMSCV-LTR-dCas9-VP64-BFP (Addgene, #46912) and pAAV-U6-sgRNA-CMV-GFP with/without sgRNAs insertion were diluted into 1ug/ul and mixed. 1ug of vector and 1ul of PLUS^TM^ reagents (Invitrogen, #15338030) were added to every 50ul Opti-MEM™ I Reduced Serum Medium. The diluted lipofectamine reagent and vectors were mixed in a 1:1 ratio and incubated for 15 min at RT to generate DNA-lipid complexes. Then 50ul and 250ul of the DNA-lipid complex were added to each well of a 24-well and 6-well plate respectively. After 12 hours for HEK293T cells and 24 hours for iPSC-induced neurons, we stopped the transfection by changing to fresh medium.

### Immunocytochemistry (ICC)

We washed wells containing cover slips with 1X TBS and then fixed them with 4% PFA and 2% sucrose in TBS for 20 min at RT. Aldehydes were quenched with 0.3 M glycine in TBS-T for 1 hour at RT. We then incubated cover slips in blocking solution (5% normal donkey serum and 3% BSA in TBS-T) for 1 hour at RT. We removed blocking solution and incubated cover slips in primary antibody solution (blocking solution with primary antibody) for 24 hours at 4°C. Secondary antibody incubations were performed overnight at 4°C followed by incubation in 300 nM DAPI solution for 5 min at RT. Coverslips were mounted in ProLong Gold Antifade Mountant. We used chicken anti-MAP2 (1:2000; Abcam, #ab5392) to label mature neurons, mouse anti-CAMKII (1:100) to label excitatory neurons, sheep anti-TENM2 (1:200) to validate expression of TENM2, sheep anti-SORCS2 (1:500) to validate expression of SORCS2, mouse anti-CUX1 (1:500; Santa Cruz Biotechnology, #sc-514008) to mark layer 2-4 like neurons, goat anti-FOXP2 (1:500) to mark layer 6 like neurons, chicken anti-GFP (1:2000; Aves Labs, #GFP-1010) to enhance GFP signal of transfected plasmid, mouse anti-TUJ1 (1:1000; Covance, #MMS-435P) to label dendrites of neurons, and rabbit anti-VGLUT2 (1:500; Synaptic Systems, #135402) to label the presynaptic puncta. We used species-matched secondary antibodies produced in donkey and conjugated to Alexa Fluor 488, Alexa Fluor 555, or Alexa Fluor 647 (1:2000). A 20X objective lens was used to capture images from cover slips on a Zeiss Axio Observer Z1 florescent microscope and a 10X objective lens was used to capture images from 6-well plates on a Leica Thunder Imager 3D Cell Culture with Tokai Stage Top Microincubator at the UT Southwestern Neuroscience Microscopy Facility. We analyzed the dendritic complexity in a 1388 x 1040 pixels (447 x 335 um) field which contains neurons (**Fig. 7A,E**) by using FIJI (Schindelin et al., 2012) with self-written Macro codes. We quantified the TUJ1 signal and normalized it to the number of DAPI+ cells that colocalized with GFP and TUJ1 signal to determine the complexity of dendritic branching. VGLUT1 puncta that overlapped with GFP and TUJ1 were counted and normalized to the length of dendrites as a quantification of spine density.

## Statistical analysis

The detail of statistical analyses were described in figure legend.

## Main Figures

The open circles always represent female samples, while closed circles represents male samples. In all analyses, we included sex as a factor in the statistical model. Since we do not find significant sex differences in most of the comparisons except for body weight and aggression, our results primarily focus on genotype or treatment effects. All data are shown as means ± SEM. Detail of statistical results can be found in **Table S3**. *\*p* < 0.05, *\*\*p* < 0.01, *\*\*\*p* < 0.001, and *\*\*\*\*p* < 0.0001.

## Data and code availability

Data of single-nuclei RNA-Seq has been deposited at GEO. We have deposited all original code to GitHub. We will provide their links and make them publicly accessible since the day of publishing. Additional information is available from the lead contact upon request.

## Acknowledgments

We thank Emily Oh for their critical comments on the manuscript, Dr. David R. Weaver for helping with mouse genotyping protocols, Emre Caglayan for help in analysis of snRNA-Seq data, Dr. Noheon Park for verifying the antibody for CLOCK WB, Dr. Filipa Ferreira, Lisa Thomas and Shelley Dixon for analysis and collection of wheel-running data. We also thank Dr. Shin Yamazaki and the UTSW Neuroscience Microscopy Facility for helping with imaging. G.K. is a Jon Heighten Scholar in Autism Research and Townsend Distinguished Chair in Research on Autism Spectrum Disorders at UT Southwestern. J.S.T. is an Investigator in the Howard Hughes Medical Institute. This work was supported by the James S. McDonnell Foundation 21^st^ Century Science Initiative in Understanding Human Cognition Scholar Award (#22002046) and National Institutes of Health (HG011641, MH207672 and MH103517) to G.K. and an American Heart Association Postdoctoral Fellowship (915654) to Y.L.

## Author contributions

Y.L., M.R.F., J.S.T., and G.K. conceptualized the project and designed experiments. Y.L. carried out most of the bench work, including mouse work, genotyping, behavior tests, AAV injection, brain dissection, brain slicing, IHC, qRT-PCR, snRNA-Seq, iPSC culture, CRIPSR-Cas9 and ICC. Y.L. collected and analyzed the data. M.R.F generated the CLOCK humanized mouse model. A.K. and Y.L. analyzed the snRNA-Seq data. N.K. and J.R.G. designed and performed electrophysiology experiments as well as data analysis. S.E.P. generated CLOCK KO iPSC lines and helped in iPSC culture and CRIPSR-Cas9 experiments. C.D. and M.H. did mouse work, genotyping, and helped with AAV injections. P.X. designed and helped with some behavior tests. N.G. contributed to brain slicing, IHC, and image taking. N.K. and S.E.P. edited the manuscript. Y.L., J.S.T., and G.K. wrote the manuscript.

## Supplementary Tables

**Table S1 (Related to Figure 1A).** SNP genotyping to determine heterogeneity after 5 generations of backcrossing with C57BL/6J mice. JJ and NN represent homozygous C57BL/6J and C57BL/6N loci, respectively, while NJ represents heterozygous loci.

**Table S2. (Related to Figure 2D).** Examples of odor and medium combinations for the set-shifting tests.

**Table S3**. Statistical results in figures.

**Table S4.(Related to Figure 4).** Gene enrichment in clusters at P07 and P56.

**Table S5. (Related to Figure 4E).** List of differentially expressed genes between HU and WT by major cell types at P07.

**Table S6. (Related to Figure 4F).** List of differentially expressed genes between KO and WT by major cell types at P07.

**Table S7. (Related to Figure 4G).** List of differentially expressed genes between HU and WT by major cell types at P56.

**Table S8. (Related to Figure 4H).** List of differentially expressed genes between KO and WT by major cell types at P56.

**Table S9. (Related to Figure S7A).** List of differentially expressed genes between HU and WT by subtype of excitatory neuron at P07.

**Table S10. (Related to Figure S7B).** List of differentially expressed genes between KO and WT by subtype of excitatory neuron at P07.

**Table S11. (Related to Figure S7C).** List of differentially expressed genes between HU and WT by subtype of excitatory neuron at P56.

**Table S12.(Related to Figure S7D).** List of differentially expressed genes between KO and WT by subtype of excitatory neuron at P56.

**Table S13. (Related to Figure 4I).** GO term enrichment of EX2/3_IT DEGs.

**Table S14. (Related to Figure S9).** Consistency between DEGs of human CLOCK in EX2/3_IT and human-specific chromatin status.

**Table S15. (Related to Figure S9).** Enhancer regions of *Tenm2/TENM2* and *Sorcs2/SORCS2* containing CLOCK:BMAL1 binding motifs.

---

**Figure S1 (Related to Figure 1).**
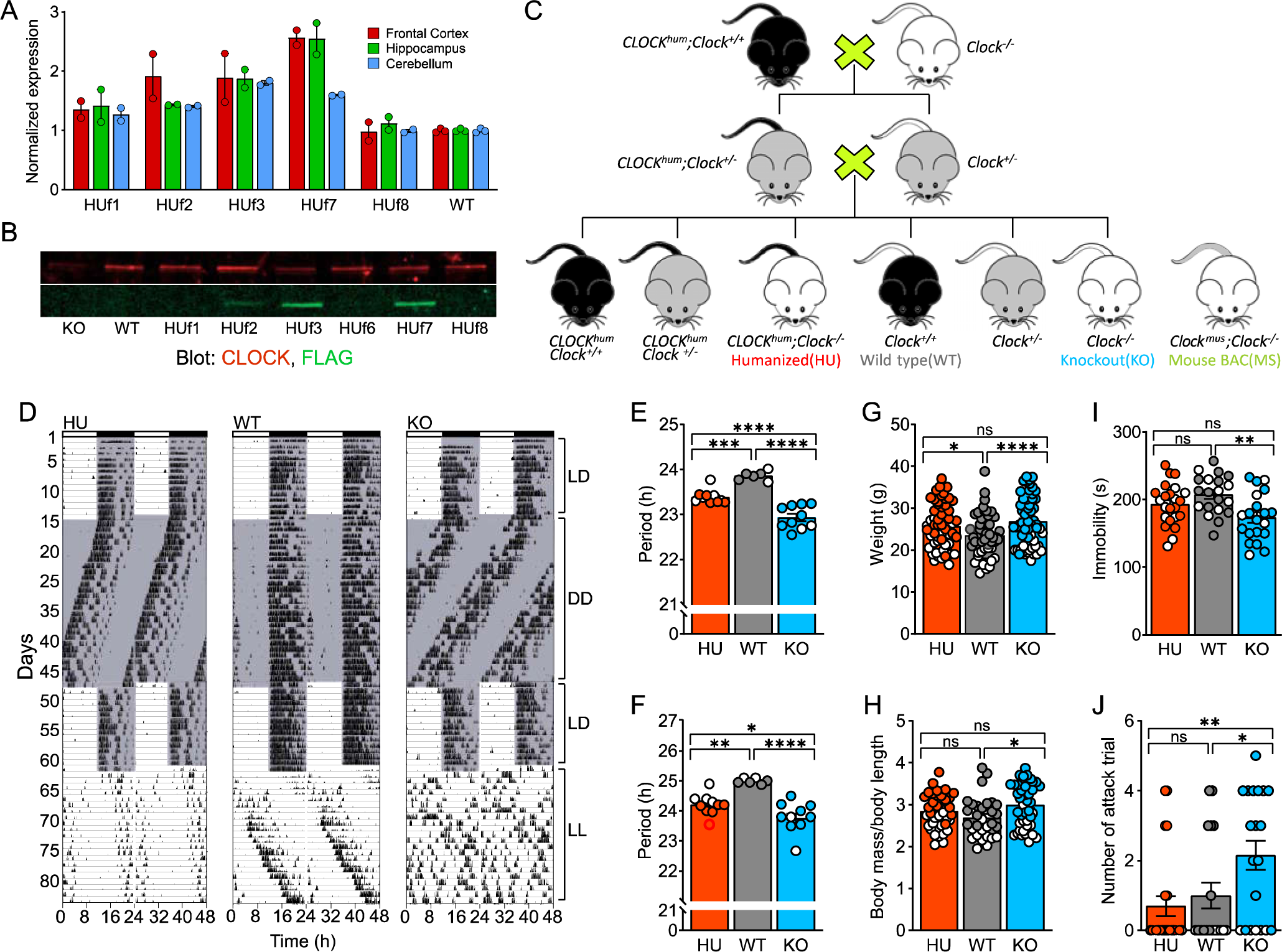
Generation and validation of mouse model. **(A)** qRT-PCR with pan-CLOCK primers for founders of each BAC transgenic mouse line. **(B)** Western blot of CLOCK (red) and FLAG (green) after immunoprecipitation with CLOCK antibody. The “f” added to the name of each mouse line represents the founder of each line. They were not bred onto the Clock KO background. **(C)** Breeding scheme of mice in this study. **(D)** Actogram of activity rhythm in wheel running test. Data are double-plotted, resulting in 48 hours for each line of trace. The black vertical lines represent activity. The shaded and unshaded areas represent lights off and on, respectively. LL represents constant light on, while DD represents constant darkness. **(E-F)** Period of activity across genotypes in **(E)** DD and **(F)** LL. **(G-H)** Body mass across genotypes with quantification of **(G)** absolute body mass and **(H)** normalized body mass to body length. **(I)** Duration of immobility in a tail suspension test. **(J)** Aggressive behavior to same-sex juvenile intruders quantified as number of trials with attack. Each data point is an individual, and we did GLM with genotype and sex as fixed factors with Tukey’s test for post-hoc analysis. See also **Table S1**.

**Figure S2 (Related to Figure 2).**
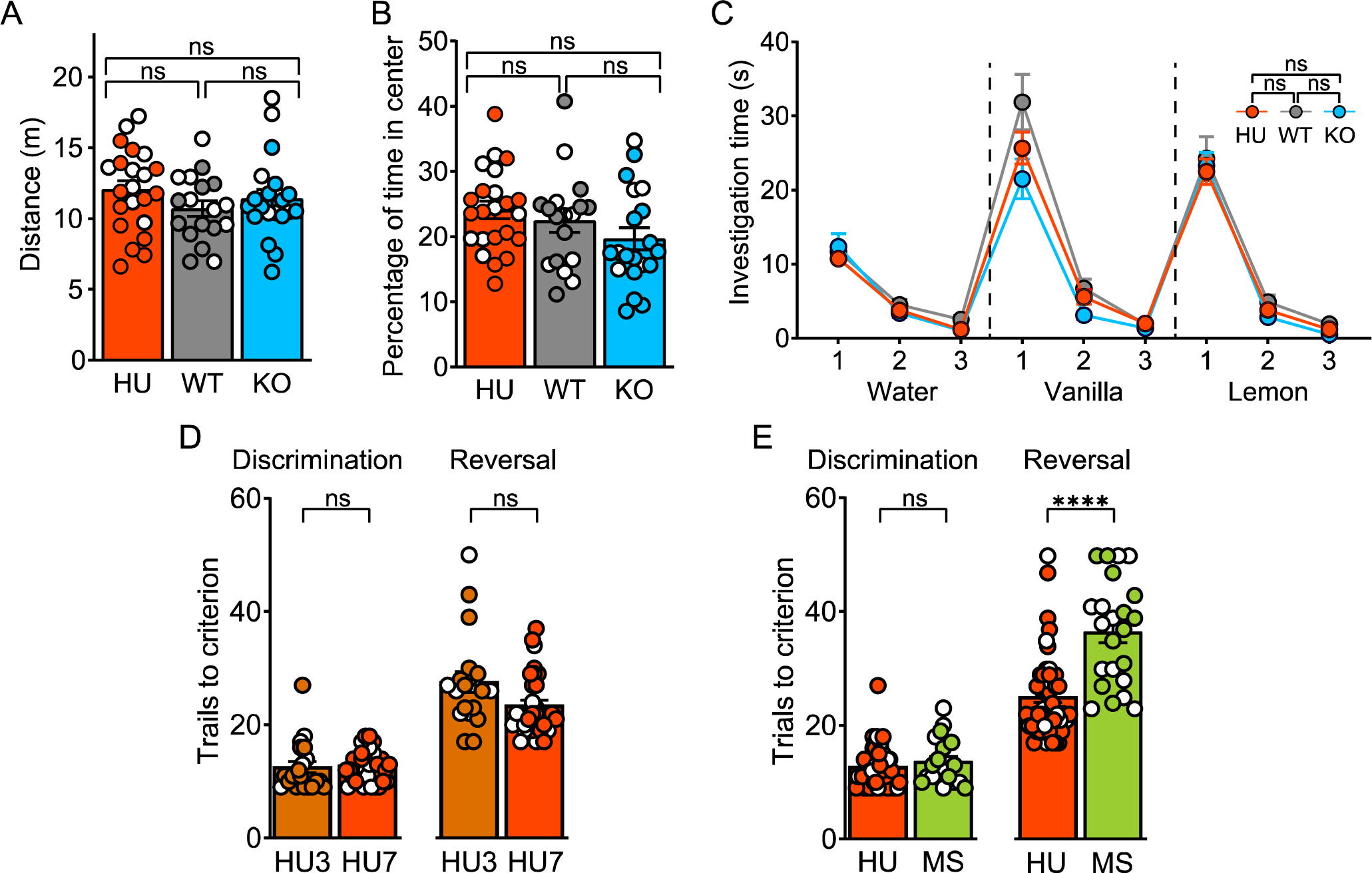
Behavioral tests. **(A-B)** CLOCK did not alter **(A)** activity and **(B)** anxiety level in an open field test. **(C)** Olfactory sensory test reveals that CLOCK does not affect olfactory recognition. **(D-E)** Comparing performance of reversal learning **(D)** between two humanized mouse lines HU3 and HU7, and **(E)** between HU and MS mice. Each data point is an individual, and we applied a one-way ANOVA with Tukey’s test for post-hoc analysis for panels **A, B, D,** and **E.** We did repeated measures ANOVA with genotype and sex as between-subject factors and trial as within-subject factor for panel **C.**

**Figure S3 (Related to Figure 3).**
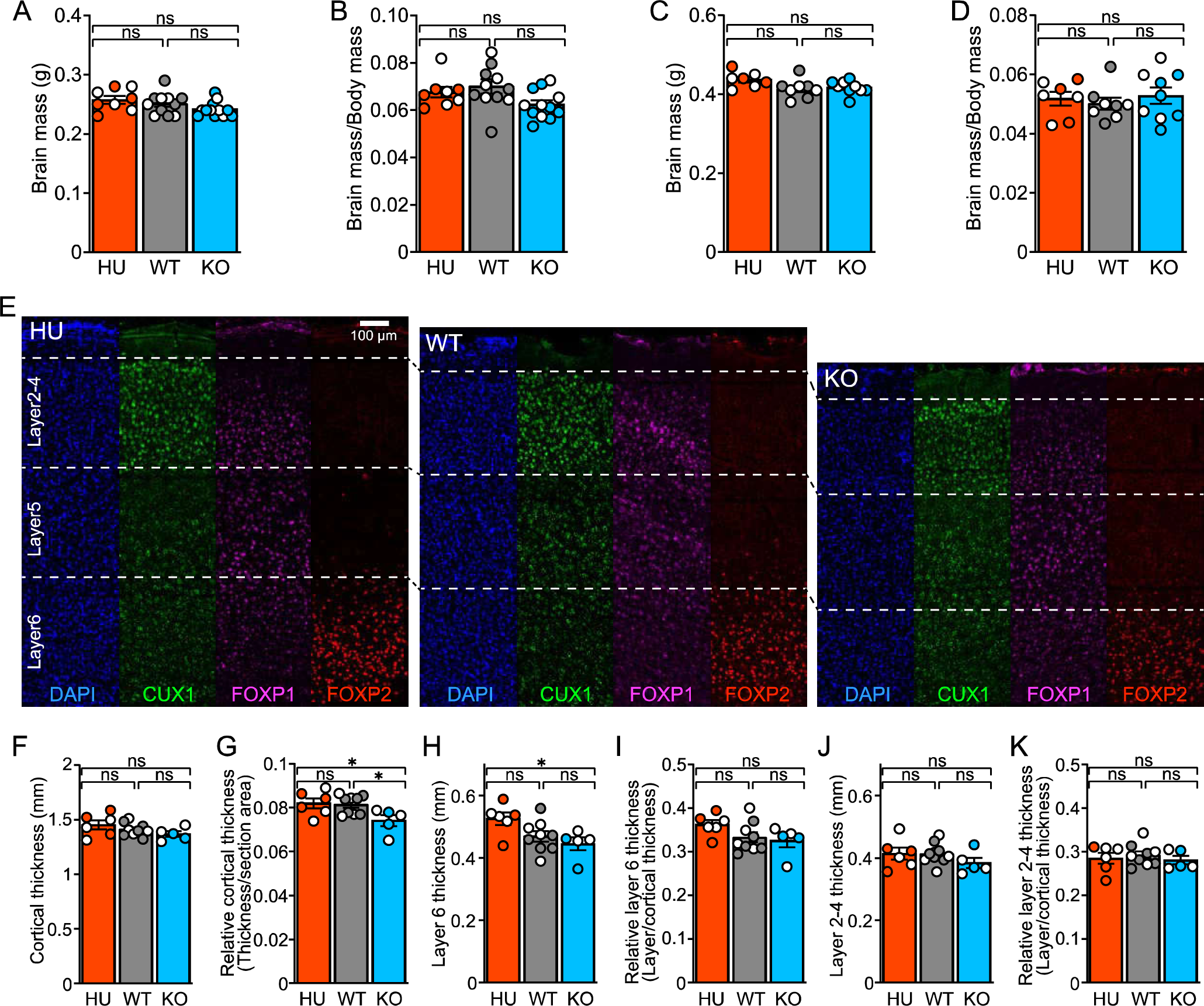
CLOCK effects on brain mass and cortical lamination. **(A)** Absolute brain mass and **(B)** normalized brain mass to body mass at PO7. **(C)** Absolute brain mass and **(D)** normalized brain mass to body mass at P21. **(E)** IHC to visualize layers 2-4 (anti-CUXΊ; staining outside of this area was in non-nuclei and considered non-specific), layers 3-5 (anti-FOXPI), and layer 6 (anti-FOXP2) in the primary somatosensory cortex. **(F)** Absolute thickness of cortex. **(G)** Relative cortical thickness normalized to area of coronal section. **(H)** Absolute thickness of layer 6. **(I)** Relative layer 6 thickness normalized to cortical thickness. **(J)** Absolute thickness of layers 2-4. **(K)** Relative layers 2-4 thickness normalized to cortical thickness. Each data point is an individual, and we applied GLM with genotype and sex as fixed factors and with Tukey’s test for post-hoc analysis.

**Figure S4 (Related to Figure 4).**
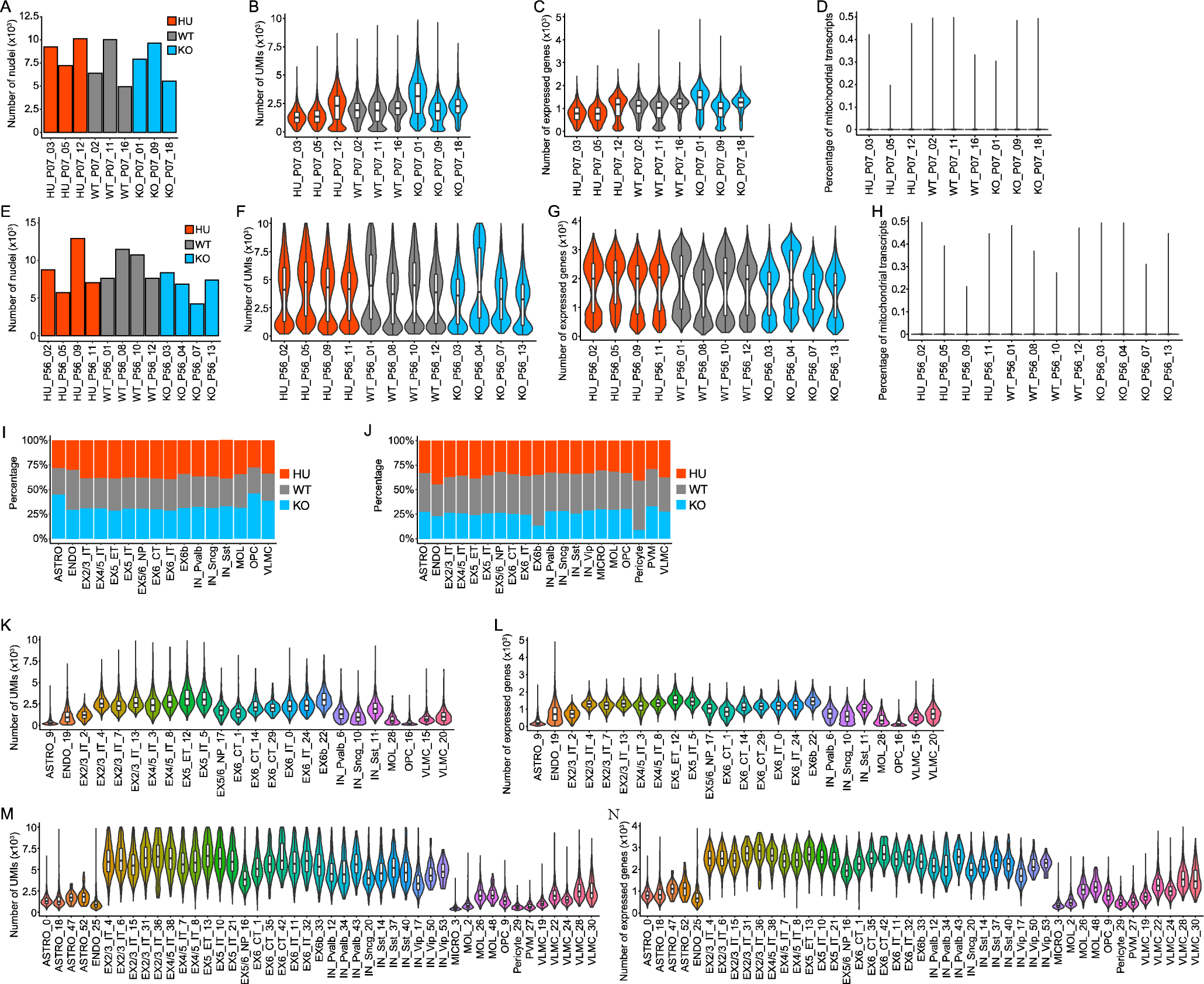
snRNA-Seq quality control plots. **(A,E)** Number of detected nuclei in each sample after sequencing at **(A)** PO7 or **(E)** P56. **(B,F)** Distribution of number of detected UMIs within each sample at **(B)** PO7 or **(F)** P56. **(C,G)** Distribution of number of expressed genes within each sample at **(C)** PO7 or **(G)** P56. **(D,H)** Distribution of percentage of mitochondrial transcripts within each sample at **(D)** PO7 or **(H)** P56. **(I-J)** Relative contribution of each genotype to the major cell at **(I)** PO7 or **(J)** P56. **(K,M)** Distribution of number of detected UMIs within each cell type at **(K)** PO7 or **(M)** P56. **(L,N)** Distribution of number expressed genes within each cell type at **(L)** PO7 or **(N)** P56.

**Figure S5 (Related to Figure 4).**
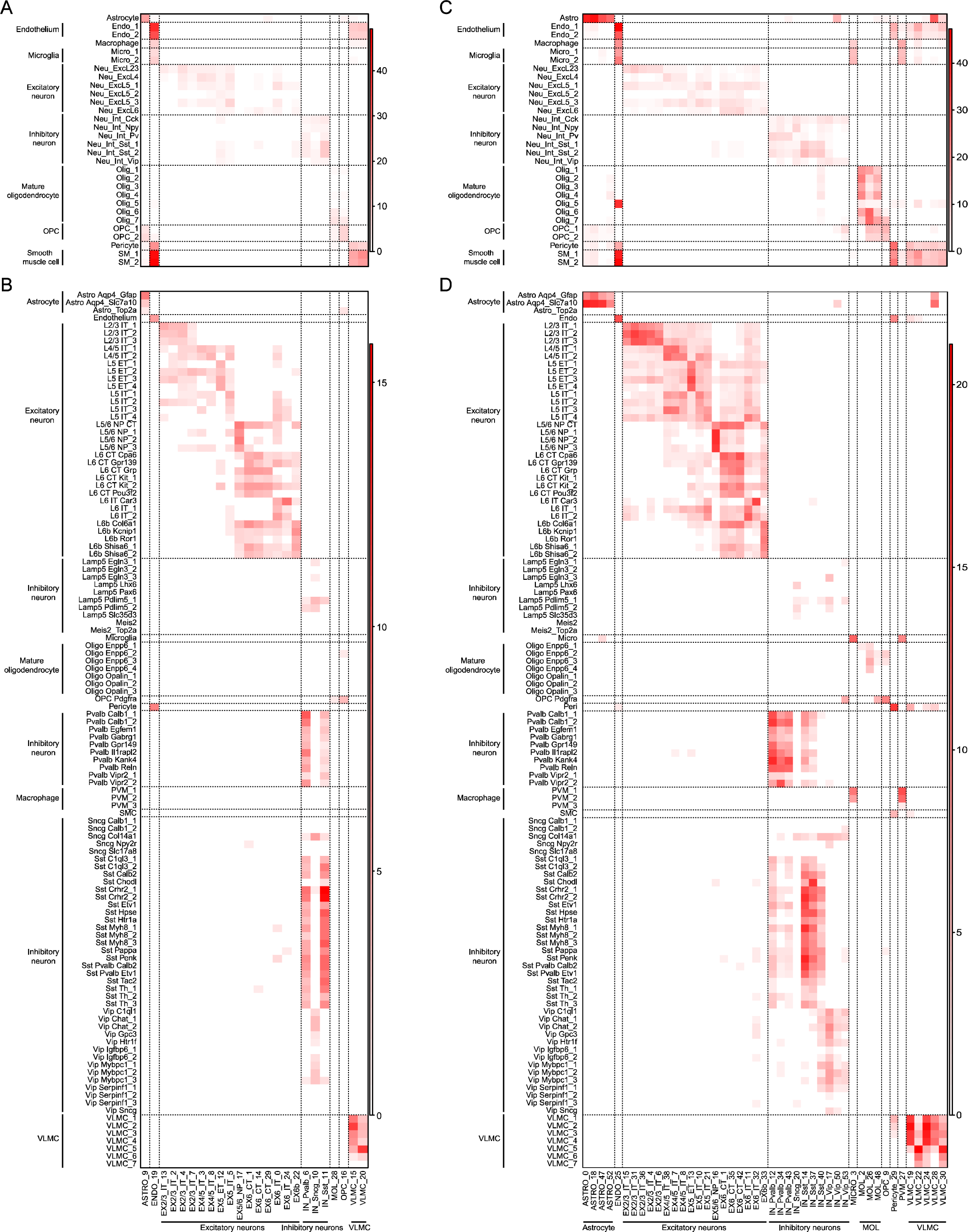
Annotation of clusters by cell types using published datasets. **(A,C)** Heatmaps show correlation between cell types from an adult mouse visual cortex dataset (Hrvatin et al., 2018) and clusters of this study at **(A)** PO7 or **(C)** P56. **(B,D)** Heatmaps show correlation between cell types from an adult mouse primary motor cortex dataset (Yao et al., 2021) and clusters of this study at **(B)** PO7 or **(D)** P56. Color reflects the −Log10 q value of a hypergeometric test. The dashed lines outline the boundaries of major cell types.

**Figure S6 (Related to Figure 4).**
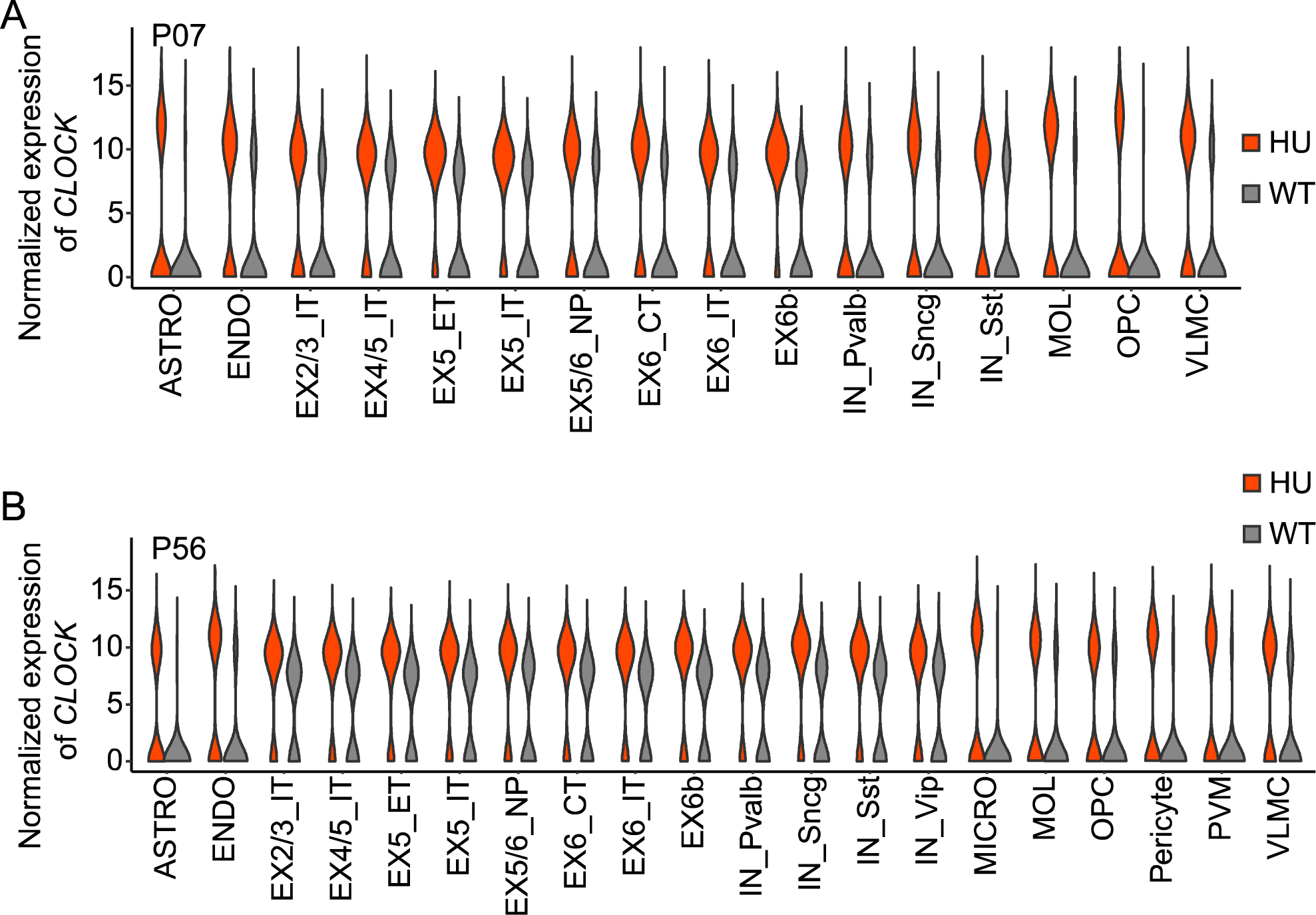
Cell type-specific expression of *CLOCK* in each genotype. Cellular expression of *CLOCK* in each cell type at **(A)** PO7 or **(B)** P56.

**Figure S7 (Related to Figure 4).**
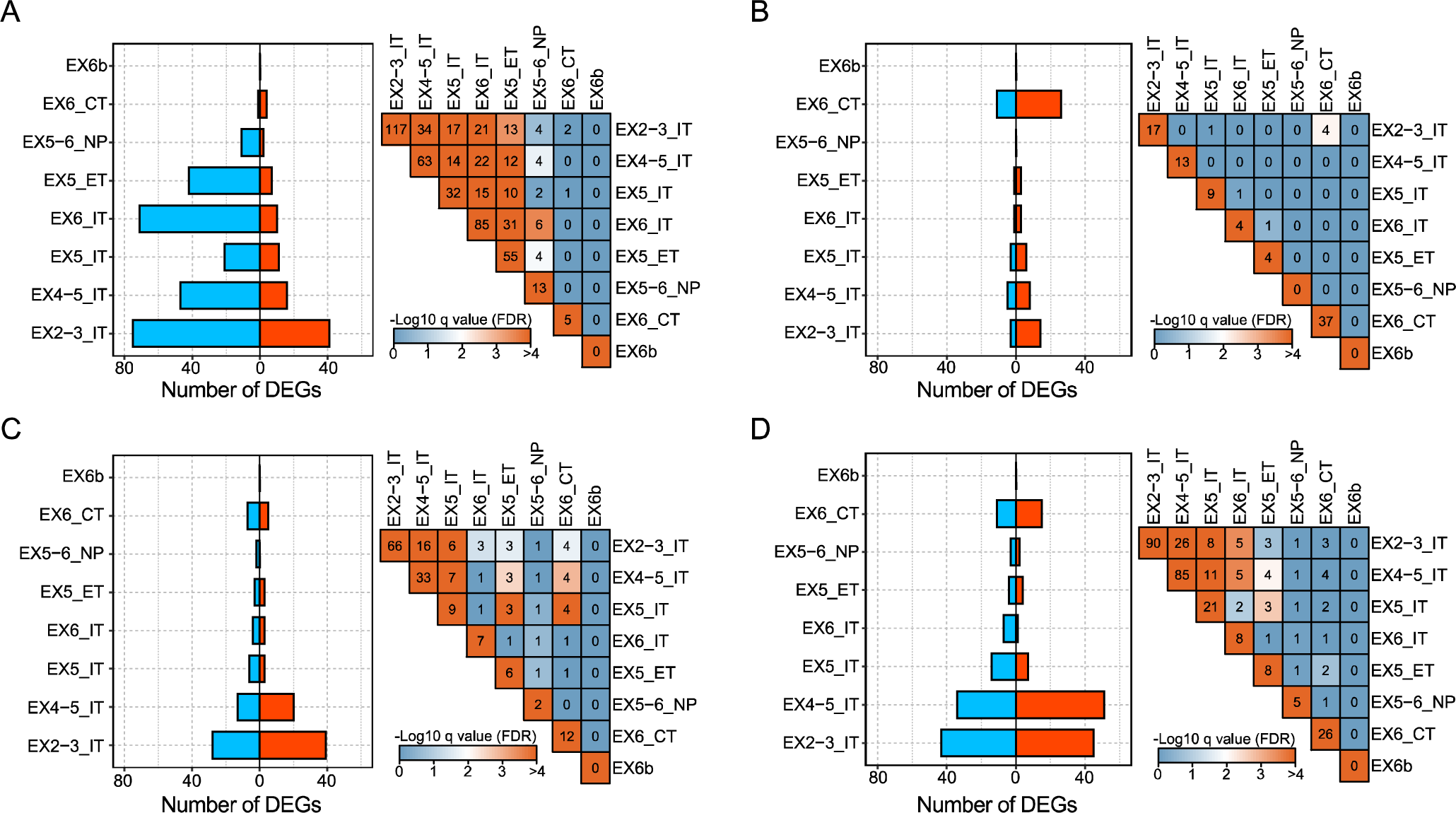
CLOCK regulates similar genes in subtypes of IT neurons with EX2-3_IT neurons most affected. Number of DEGs in each subtype of excitatory neurons (bar plot), and hypergeometric tests between DEGs from each pair of cell types (heatmap) at **(A)** PO7 HU vs. WT comparison, **(B)** PO7 KO vs. WT comparison, **(C)** P56 HU vs. WT comparison, and **(D)** P56 KO vs. WT comparison. Color reflects the −Log10 q value of hypergeometric test in heatmaps. See also **Tables S9-12**.

**Figure S8 (Related to Figure 4).**
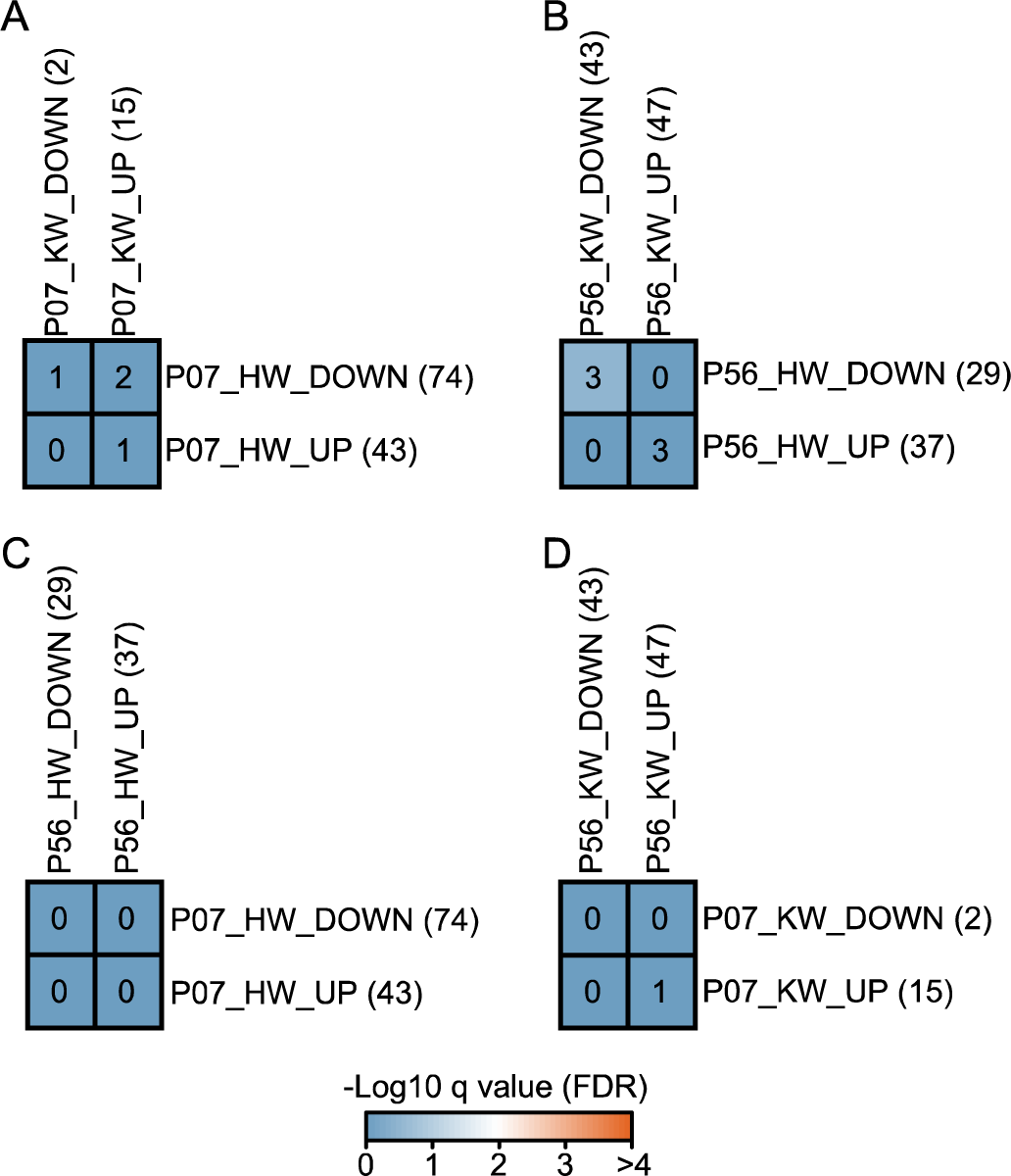
Human and mouse CLOCK regulate different genes in EX2-3_IT neurons in an age-specific manner. Overlap of DEGs from human and mouse CLOCK at **(A)** PO7 or **(B)** P56. **(C)** Overlap of DEGs result from human CLOCK between PO7 and P56. **(D)** Overlap of DEGs from mouse Clock between PO7 and P56. The color of each cell represents the −Log10 q value of hypergeometric test. The number in each cell is the number of overlapped DEGs. The number in brackets are the number of DEGs.

**Figure S9 (Related to Figure 4).**
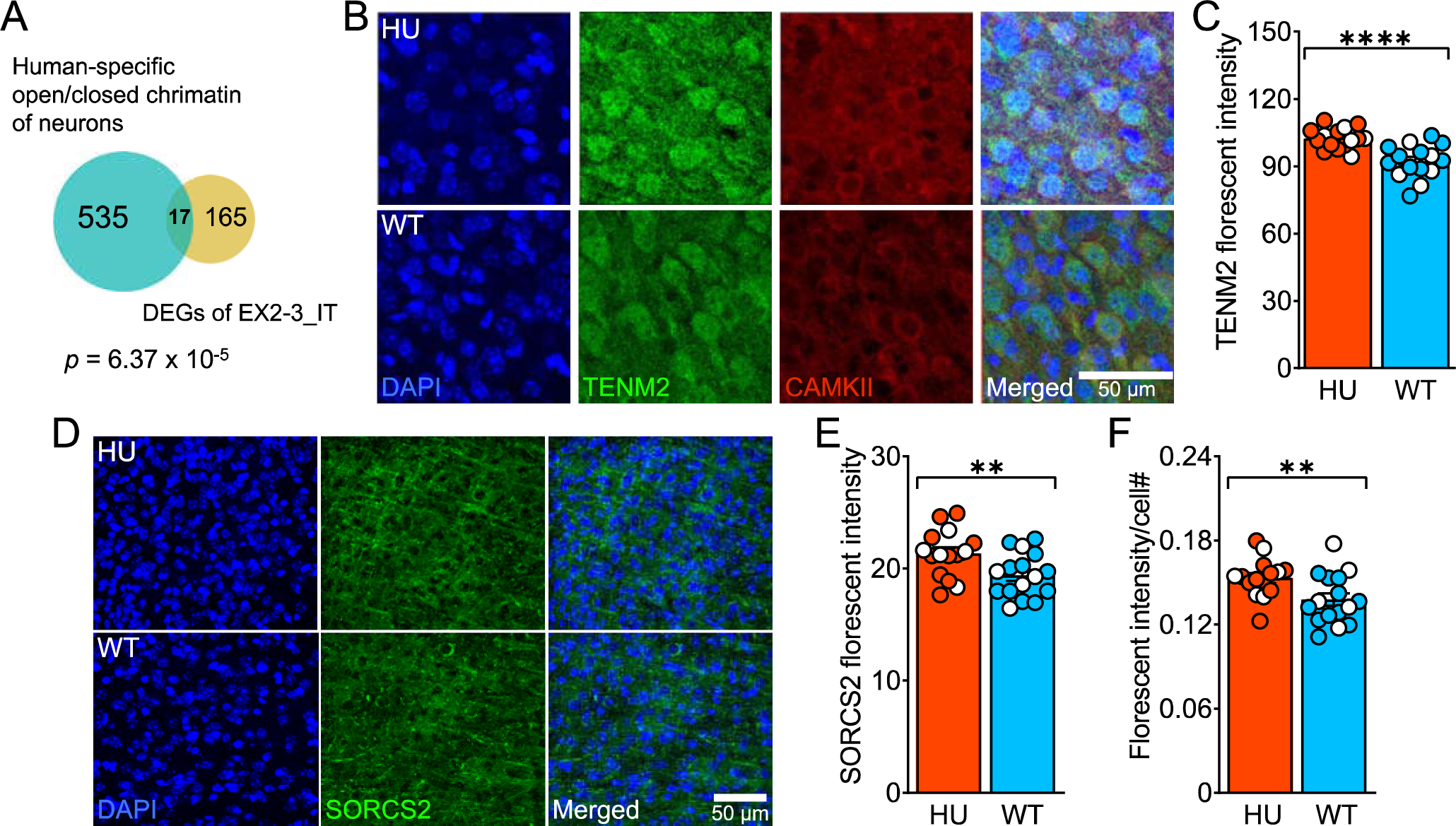
Validation of protein expression of human CLOCK downstream targets. **(A)** Venn diagram to show overlap between DEGs of EX2/3_IT and human-specific open chromatin in neurons. **(B)** Representative images to compare TENM2 fluorescent intensity (green) between HU and WT mice in excitatory neurons (CAMKII, red) of frontal cortex at PO7. **(C)** Quantification of TENM2 fluorescent intensity. Each data point is an excitatory neuron, and we sampled 3 mice per genotype and 5 neurons per mouse. **(D)** Representative images to compare SORCS2 fluorescent intensity (green) between HU and WT mice in the frontal cortex at P56. **(E-F)** Quantification of **(E)** absolute and **(F)** normalized SORCS2 fluorescent intensity. Each data point is an image of a frontal cortex section, and we sampled 3 mice per genotype and 5-6 images per mouse. We did GLMM with genotype and sex as fixed factors, image as random factor nested with individual, and Tukey’s test for post-hoc analysis. See also **Table S14-15**.

**Figure S10 (Related to Figure 5).**
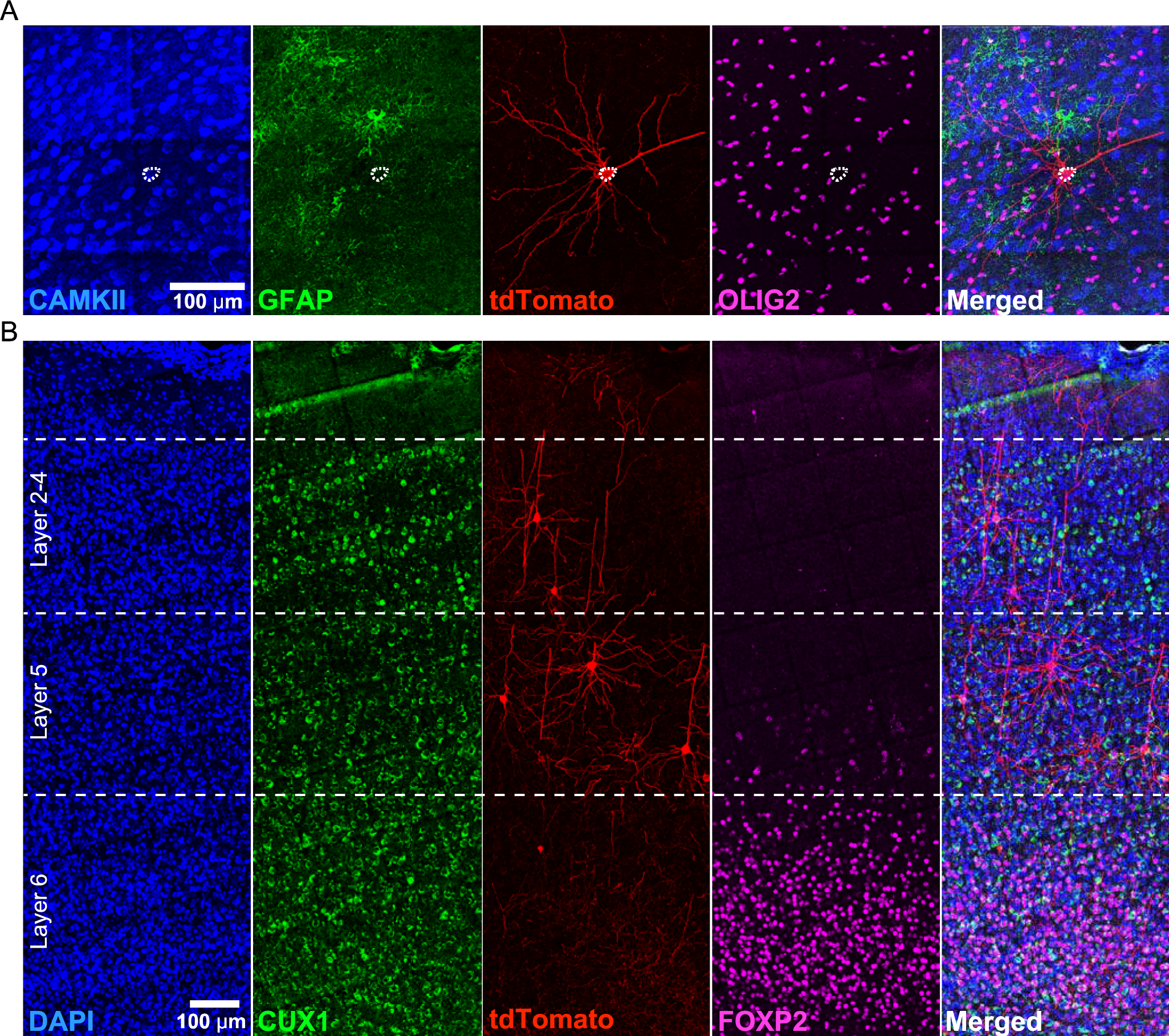
Identity and location of the neurons for morphological quantification. **(A)** AAV transduced (tdTomato, red) excitatory neurons, but not astrocytes (GFAP, green) and oligodendrocytes (OLIG2, magenta). **(B)** AAV transduced (tdTomato, red) excitatory neurons across cortical layers (layers 2-4: CUX1, green; layer 6: FOXP2, magenta) of adult frontal cortex.

**Figure S11 (Related to Figure 5).**
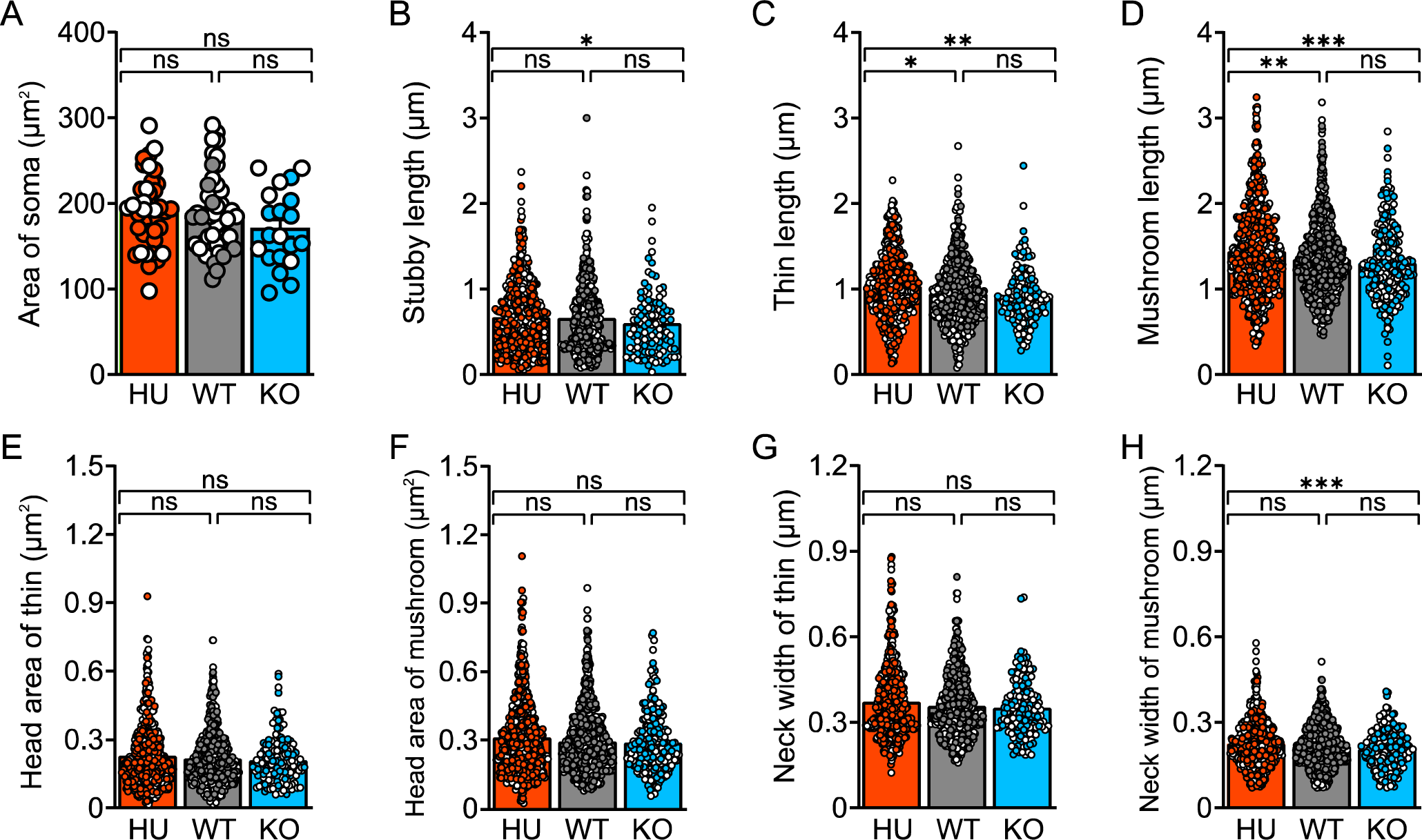
Soma area and spine morphology of layer 2-4 excitatory neurons in adult mice. **(A)** Comparison of soma area. Each data point is a neuron, and we sampled 3-4 mice per genotype and 6-12 neurons per mice. **(B-D)** Comparison of length across genotypes in **(B)** stubby, **(C)** thin, and **(D)** mushroom spines. **(E-F)** Head area of **(E)** thin and **(F)** mushroom spines. **(G-H)** Neck width of **(G)** thin and **(H)** mushroom spines. Each data point is a spine, and we sampled all spines from 2-3 segments from 3-7 neurons per mouse. We did GLMM with genotype and sex as fixed factors, spine as random factor nested with segment which again nested with neuron which further nested with individual, and Tukey’s test for post-hoc analysis.

**Figure S12 (Related to Figure 5).**
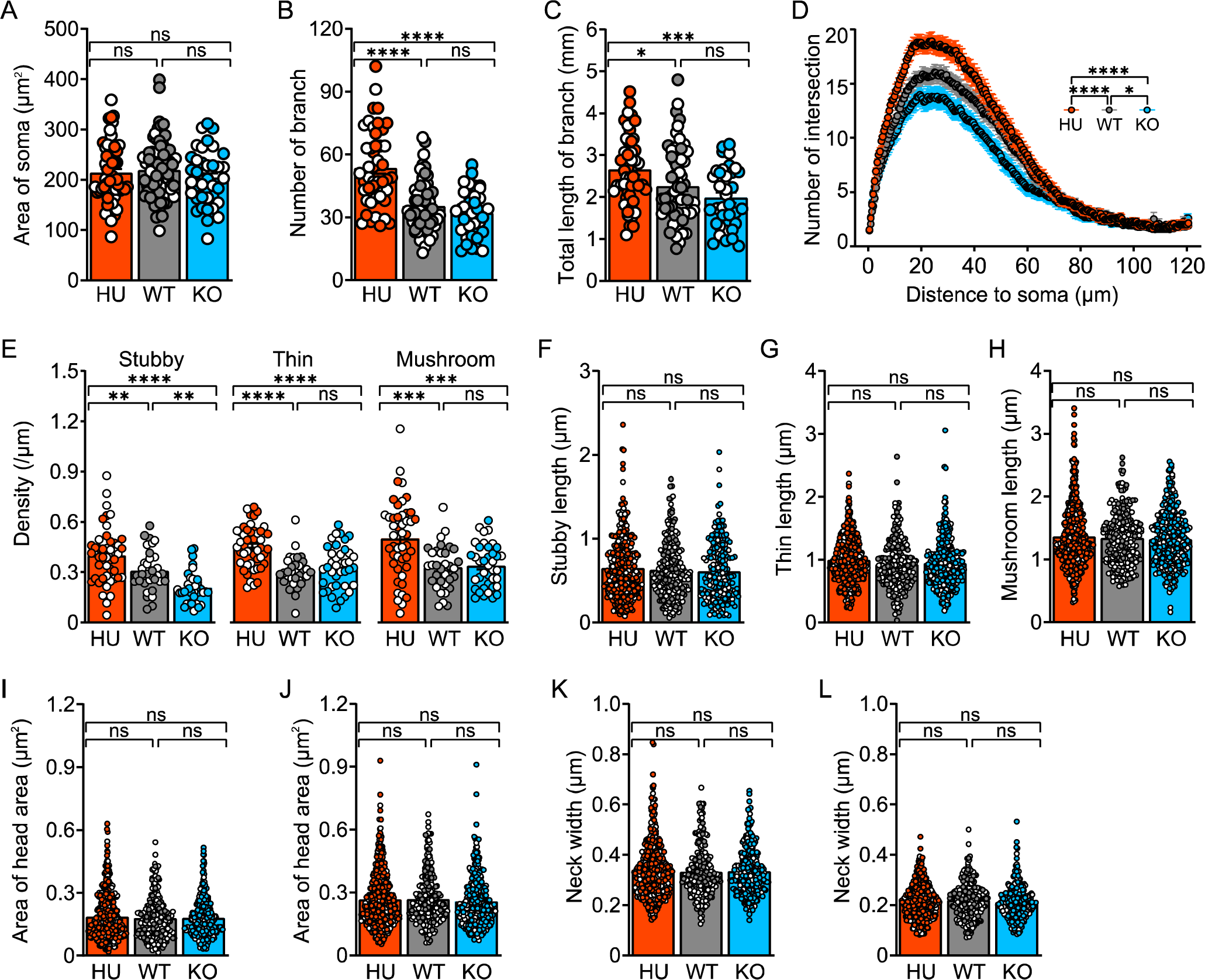
Neuronal morphology of layer 2-4 excitatory neurons in frontal cortex at P18. **(A)** CLOCK does not alter soma area. **(B-C)** Quantification of **(B)** number and **(C)** total length of branch in each genotypes. Each data point is a neuron, and we sampled 4 mice per genotype and 7-17 neurons per mouse. We did GLMM with genotype and sex as fixed factors, neuron as random factor nested with individual, and Tukey’s test for post-hoc analysis. **(D)** Sholl analysis to quantify encountered intersections between neuron branches and concentric rings from soma for dendrite complexity. We applied GLMM with genotype, sex and distance to soma (x-axis) as fixed factors, segment as random factor nested with neuron which nested with individual, and Tukey’s test for post-hoc analysis. **(E)** Density of each subtype of spine in each genotype. Each data point is a segment, and we sampled 2-3 segments from 3-5 neurons per mouse. **(F-H)** Comparison of length across genotypes in **(F)** stubby, **(G)** thin, and **(H)** mushroom spines. **(I-J)** Head area of **(I)** thin and **(J)** mushroom spines. **(K-L)** Neck width of **(K)** thin and **(L)** mushroom spines. Each data point is a spine, and we sampled all spines from 2-3 segments from 3-5 neurons per mouse. We did GLMM with genotype and sex as fixed factors, spine as random factor nested with segment which again nested with neuron which further nested with individual, and Tukey’s test for post-hoc analysis.

**Figure S13 (Related to Figure 6).**
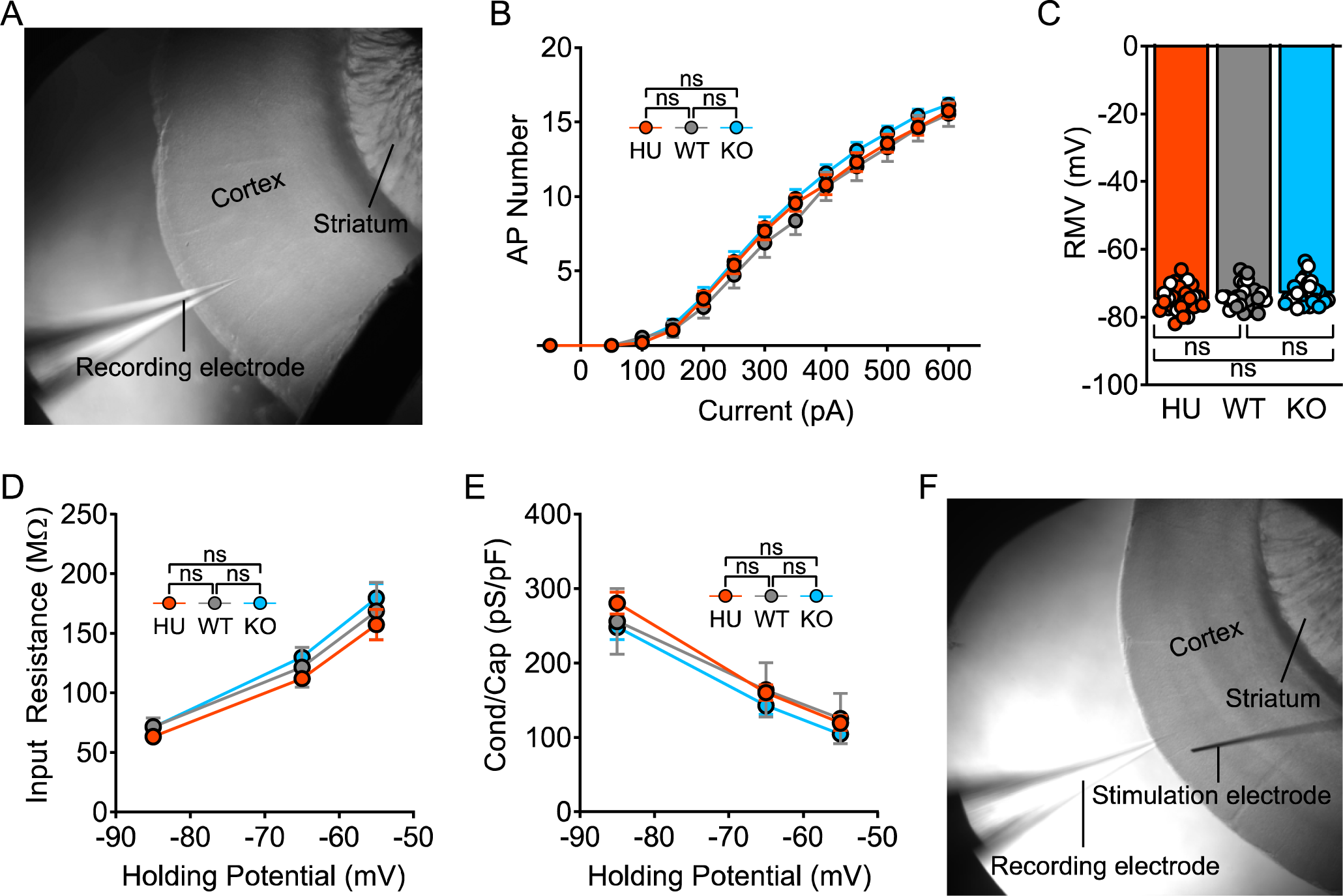
CLOCK does not affect the intrinsic properties of layer 2-4 excitatory neurons. **(A)** The microscope image shows our method to patch layer 2-4 neurons in the primary somatosensory cortex. **(B)** The number of action potential (AP) versus current amplitude curves indicates that CLOCK does not affect intrinsic excitability. **(C-E)** CLOCK does not alter **(C)** resting membrane voltage (RMV), **(D)** input resistance, and **(E)** conductance, indicating unchanged membrane properties. **(F)** The microscope image shows our approach to stimulate neurons in the same layer as the recording. Each data point is a neuron, and we sampled 4-5 mice per genotype and 4-8 neurons per mouse. We perform GLMM with genotype, sex, and x-axis (only for panels **B, D**, and **E**) as fixed factors, neuron as a random factor nested with individual, and Tukey’s test for post-hoc analysis.

**Figure S14 (Related to Figure 7).**
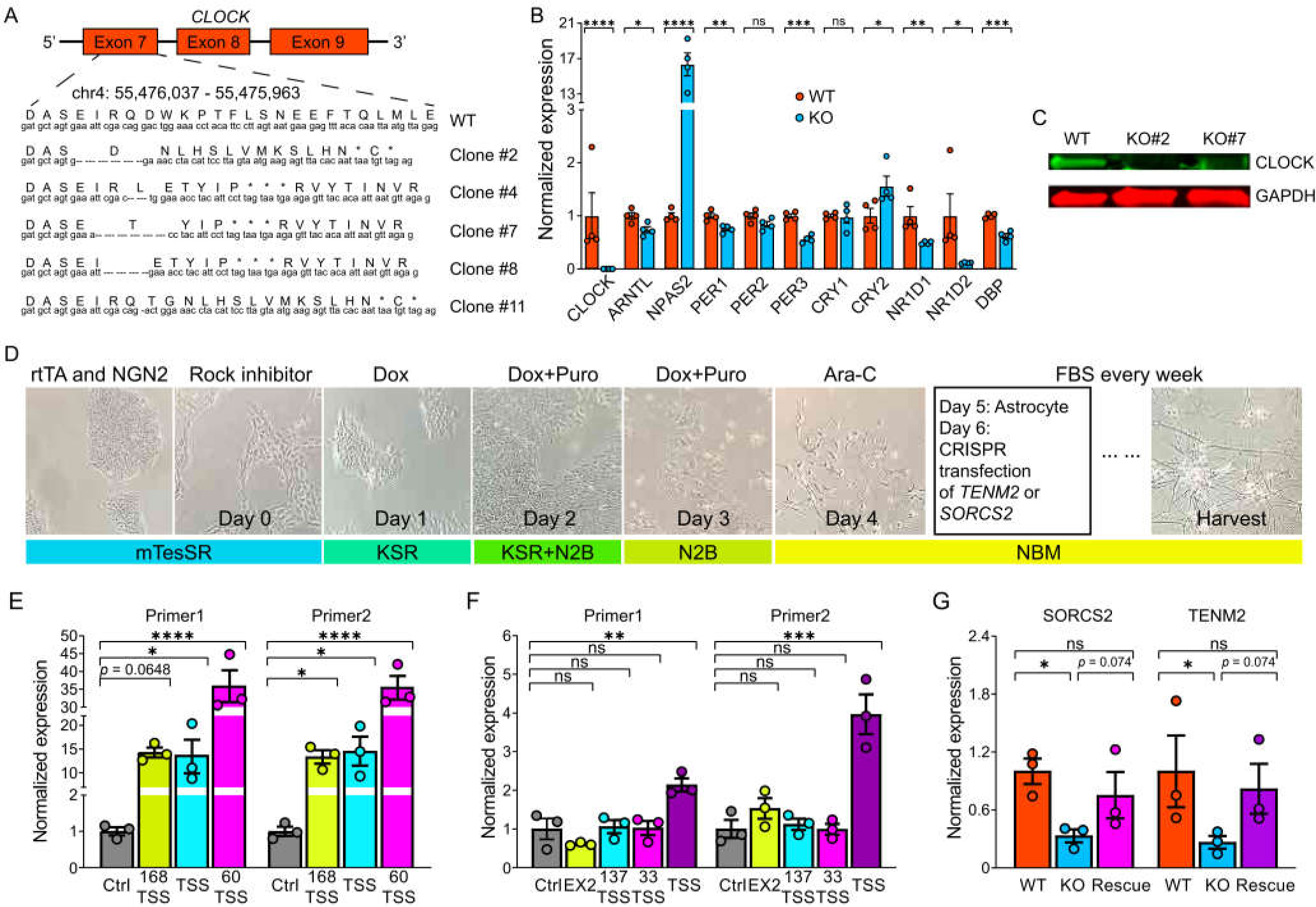
Generation of CLOCK KO iPSCs, differentiation to neurons, and CRISPR-Cas9 overexpression of TENM2 and SORCS2. **(A)** Schematic of sgRNAs using CRISPR-Cas9 targeting and deleted short nucleotide sequences, resulting in frame shifting in different clones. The upper line of capitalized characters are protein sequences, while the bottom lower-case characters are nucleotide sequences. Each dash symbol represents a deleted nucleotide, and the star sign represents a stop codon. **(B)** qRT-PCR shows CLOCK is not expressed in CLOCK KO iPSCs, and expression of most core circadian genes is altered. Each data point is a culture of iPSCs in a well on 24-well plate, and we used a Student’s t test. **(C)** Western blot of CLOCK (green) and GAPDH (red) İPSC lines. **(D)** Schematic of NGN2 fast neuron induction protocol to differentiate iPSCs to mature neurons. Images show cell and colony morphology across differentiation stages. Words above images are reagents that were added in each stage, while words below images are the culture medium corresponding to each stage. **(E-F)** qRT-PCR to test efficiency of sgRNAs that target possible promotor regions (x-axis) to overexpress **(E)** SORCS2 or **(F)** TENM2 in 293T cells. For each gene, we used two pairs of primers for qRT-PCR. TSS: transcription start site, and the number represents the nucleotide distance from the TSS. EX2, target in exon 2. Ctrl, empty plasmid without sgRNA. **(G)** qRT-PCR shows that expression of both SORCS2 and TENM2 decrease in CLOCK KO induced neurons while sgRNA overexpressioπ partially rescued their expression. Each data point is a culture of iPSCs in a well of a 24-well plate, and we applied Kruskal Wallis Test with Dunn’s test for post-hoc analysis.

**Figure S15 (Related to Figure 7).**
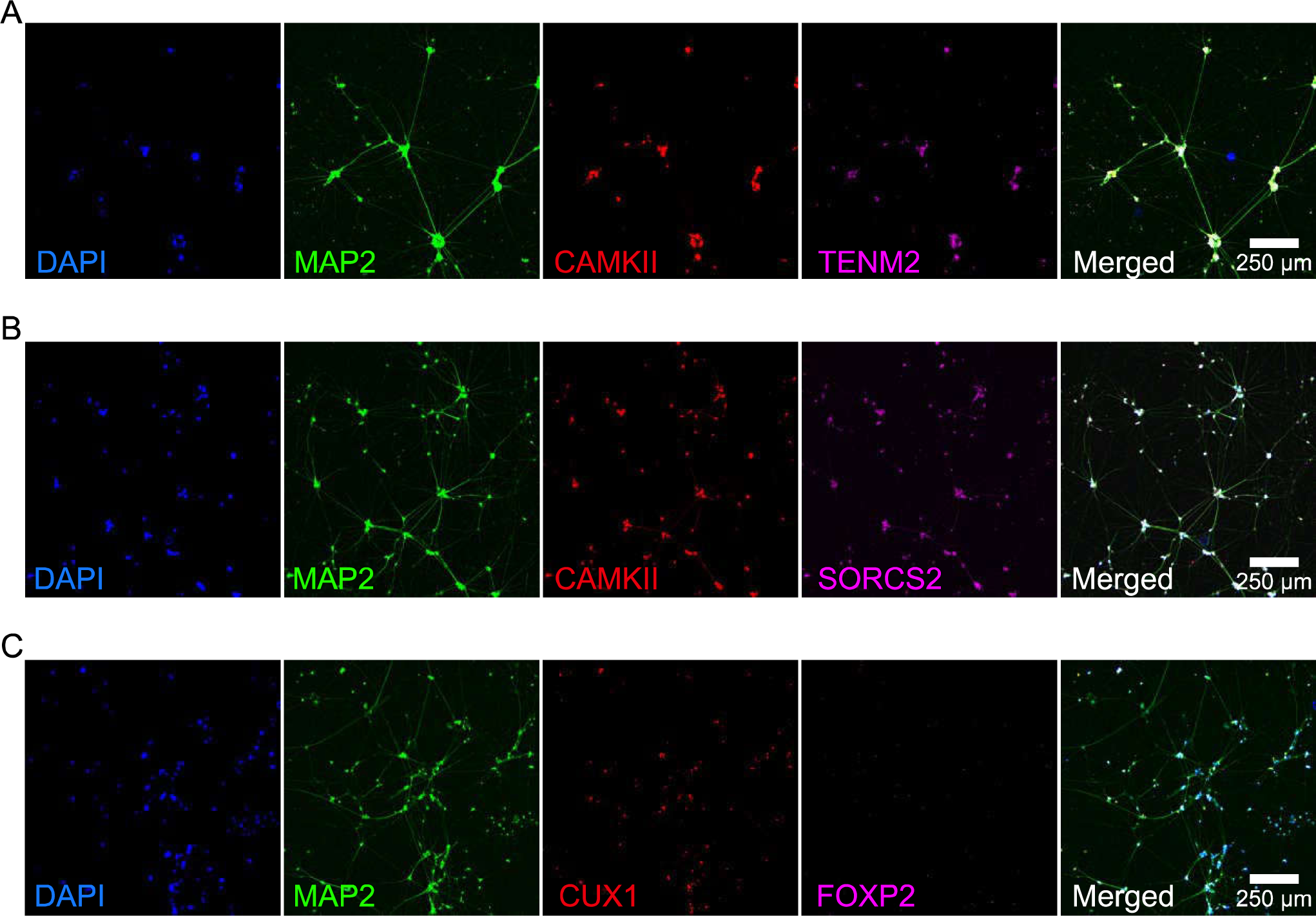
Identity of iPSC-derived neurons at Day 28 of differentiation. **(A-B)** The iPSC-derived neurons express neuronal markers of mature (MAP2, green) and excitatory (CAMKII, red) neurons with expression of **(A)** TENM2 and **(B)** SORCS2. **(C)** The iPSC-derived neurons express neuronal markers of layers 2-4 (CUX1, red) rather than layer 6 (FOXP2, magenta) cortical neurons.

## References

1. Antoch, M.P., Song, E.-J., Chang, A.-M., Vitaterna, M.H., Zhao, Y., Wilsbacher, L.D., Sangoram, A.M., King, D.P., Pinto, L.H., and Takahashi, J.S. (1997). Functional Identification of the Mouse Circadian Clock Gene by Transgenic BAC Rescue. Cell 89, 655–667.

2. Antunes, M., and Biala, G. (2012). The novel object recognition memory: neurobiology, test procedure, and its modifications. Cognitive Processing 13, 93–110.

3. Arshadi, C., Günther, U., Eddison, M., Harrington, K.I.S., and Ferreira, T.A. (2021). SNT: a unifying toolbox for quantification of neuronal anatomy. Nature Methods 18, 374–377.

4. Ayhan, F., Kulkarni, A., Berto, S., Sivaprakasam, K., Douglas, C., Lega, B.C., and Konopka, G. (2021). Resolving cellular and molecular diversity along the hippocampal anterior-to-posterior axis in humans. Neuron 109, 2091–2105.

5. Babbitt, C.C., Fedrigo, O., Pfefferle, A.D., Boyle, A.P., Horvath, J.E., Furey, T.S., and Wray, G.A. (2010). Both noncoding and protein-coding RNAs contribute to gene expression evolution in the primate brain. Genome Biology and Evolution 2, 67–79.

6. Bakken, T.E., Jorstad, N.L., Hu, Q., Lake, B.B., Tian, W., Kalmbach, B.E., Crow, M., Hodge, R.D., Krienen, F.M., Sorensen, S.A., et al. (2021). Comparative cellular analysis of motor cortex in human, marmoset and mouse. Nature 598, 111–119.

7. Baloch, S., Verma, R., Huang, H., Khurd, P., Clark, S., Yarowsky, P., Abel, T., Mori, S., and Davatzikos, C. (2008). Quantification of Brain Maturation and Growth Patterns in C57BL/6J Mice via Computational Neuroanatomy of Diffusion Tensor Images. Cerebral Cortex 19, 675–687.

8. Becht, E., McInnes, L., Healy, J., Dutertre, C.-A., Kwok, I.W.H., Ng, L.G., Ginhoux, F., and Newell, E.W. (2018). Dimensionality reduction for visualizing single-cell data using UMAP. Nature Biotechnology.

9. Berry, K.P., and Nedivi, E. (2017). Spine Dynamics: Are They All the Same? Neuron 96, 43–55.

10. Berto, S., and Nowick, K. (2018). Species-specific changes in a primate transcription factor network provide insights into the molecular evolution of the primate prefrontal cortex. Genome Biology and Evolution 10, 2023–2036.

11. Bianchi, S., Stimpson, C.D., Duka, T., Larsen, M.D., Janssen, W.G.M., Collins, Z., Bauernfeind, A.L., Schapiro, S.J., Baze, W.B., McArthur, M.J., et al. (2013). Synaptogenesis and development of pyramidal neuron dendritic morphology in the chimpanzee neocortex resembles humans. Proceedings of the National Academy of Sciences 110, 10395–10401.

12. Bissonette, G.B., and Powell, E.M. (2012). Reversal learning and attentional set-shifting in mice. Neuropharmacology 62, 1168–1174.

13. Boldog, E., Bakken, T.E., Hodge, R.D., Novotny, M., Aevermann, B.D., Baka, J., Bordé, S., Close, J.L., Diez-Fuertes, F., Ding, S.-L., et al. (2018). Transcriptomic and morphophysiological evidence for a specialized human cortical GABAergic cell type. Nature Neuroscience 21, 1185–1195.

14. Bozek, K., Wei, Y., Yan, Z., Liu, X., Xiong, J., Sugimoto, M., Tomita, M., Pääbo, S., Pieszek, R., Sherwood, C.C., et al. (2014). Exceptional Evolutionary Divergence of Human Muscle and Brain Metabolomes Parallels Human Cognitive and Physical Uniqueness. PLOS Biology 12, e1001871.

15. Butler, A., Hoffman, P., Smibert, P., Papalexi, E., and Satija, R. (2018). Integrating single-cell transcriptomic data across different conditions, technologies, and species. Nature Biotechnology 36, 411–420.

16. Caglayan, E., Liu, Y., and Konopka, G. (2022). Neuronal ambient RNA contamination causes misinterpreted and masked cell types in brain single-nuclei datasets. Neuron.

17. Can, A., Dao, D.T., Terrillion, C.E., Piantadosi, S.C., Bhat, S., and Gould, T.D. (2012). The Tail Suspension Test. Journal of Visualized Experiments: JoVE, 3769.

18. Charrier, C., Joshi, K., Coutinho-Budd, J., Kim, J.-E., Lambert, N., de Marchena, J., Jin, W.-L., Vanderhaeghen, P., Ghosh, A., Sassa, T., et al. (2012). Inhibition of SRGAP2 Function by Its Human-Specific Paralogs Induces Neoteny during Spine Maturation. Cell 149, 923–935.

19. Chen, C.-Y., Logan, R.W., Ma, T., Lewis, D.A., Tseng, G.C., Sibille, E., and McClung, C.A. (2016). Effects of aging on circadian patterns of gene expression in the human prefrontal cortex. Proceedings of the National Academy of Sciences 113, 206–211.

20. Chen, J., Bardes, E.E., Aronow, B.J., and Jegga, A.G. (2009). ToppGene Suite for gene list enrichment analysis and candidate gene prioritization. Nucleic Acids Research 37, W305–W311.

21. Chung, S., Lee, Eun J., Yun, S., Choe, Han K., Park, S.-B., Son, Hyo J., Kim, K.-S., Dluzen, Dean E., Lee, I., Hwang, O., et al. (2014). Impact of Circadian Nuclear Receptor REV-ERBα on Midbrain Dopamine Production and Mood Regulation. Cell 157, 858–868.

22. Concordet, J.-P., and Haeussler, M. (2018). CRISPOR: intuitive guide selection for CRISPR/Cas9 genome editing experiments and screens. Nucleic Acids Research 46, W242–W245.

23. Consortium, T.C.S.a.A. (2005). Initial sequence of the chimpanzee genome and comparison with the human genome. Nature 437, 69–87.

24. Dallmann, R., DeBruyne, J.P., and Weaver, D.R. (2011). Photic resetting and entrainment in CLOCK-deficient mice. J Biol Rhythms 26, 390–401.

25. DeBruyne, J.P., Noton, E., Lambert, C.M., Maywood, E.S., Weaver, D.R., and Reppert, S.M. (2006). A clock shock: mouse CLOCK is not required for circadian oscillator function. Neuron 50, 465–477.

26. del Toro, D., Carrasquero-Ordaz, M.A., Chu, A., Ruff, T., Shahin, M., Jackson, V.A., Chavent, M., Berbeira-Santana, M., Seyit-Bremer, G., Brignani, S., et al. (2020). Structural Basis of Teneurin-Latrophilin Interaction in Repulsive Guidance of Migrating Neurons. Cell 180, 323–339.e319.

27. Dennis, Megan Y., Nuttle, X., Sudmant, Peter H., Antonacci, F., Graves, Tina A., Nefedov, M., Rosenfeld, Jill A., Sajjadian, S., Malig, M., Kotkiewicz, H., et al. (2012). Evolution of Human-Specific Neural SRGAP2 Genes by Incomplete Segmental Duplication. Cell 149, 912–922.

28. Dobin, A., Davis, C.A., Schlesinger, F., Drenkow, J., Zaleski, C., Jha, S., Batut, P., Chaisson, M., and Gingeras, T.R. (2012). STAR: ultrafast universal RNA-seq aligner. Bioinformatics 29, 15–21.

29. Dougherty, J.D., Schmidt, E.F., Nakajima, M., and Heintz, N. (2010). Analytical approaches to RNA profiling data for the identification of genes enriched in specific cells. Nucleic Acids Research 38, 4218–4230.

30. Dicke, U., and Roth, G. (2016). Neuronal factors determining high intelligence. Philosophical Transactions of the Royal Society B: Biological Sciences 371, 20150180.

31. Easton, A., Arbuzova, J., and Turek, F.W. (2003). The circadian Clock mutation increases exploratory activity and escape-seeking behavior. Genes, Brain and Behavior 2, 11–19.

32. Elston, G.N., Benavides-Piccione, R., and DeFelipe, J. (2001). The Pyramidal Cell in Cognition: A Comparative Study in Human and Monkey. The Journal of Neuroscience 21, RC163–RC163.

33. Elston, G.N., Benavides-Piccione, R., Elston, A., Zietsch, B., Defelipe, J., Manger, P., Casagrande, V., and Kaas, J.H. (2006). Specializations of the granular prefrontal cortex of primates: Implications for cognitive processing. The Anatomical Record Part A: Discoveries in Molecular, Cellular, and Evolutionary Biology 288A, 26–35.

34. Enard, W., Gehre, S., Hammerschmidt, K., Hölter, S.M., Blass, T., Somel, M., Brückner, M.K., Schreiweis, C., Winter, C., and Sohr, R. (2009). A humanized version of Foxp2 affects cortico-basal ganglia circuits in mice. Cell 137, 961–971.

35. Enard, W., Przeworski, M., Fisher, S.E., Lai, C.S., Wiebe, V., Kitano, T., Monaco, A.P., and Pääbo, S. (2002). Molecular evolution of *FOXP2*, a gene involved in speech and language. Nature 418, 869–872.

36. Ferreira, T.A., Blackman, A.V., Oyrer, J., Jayabal, S., Chung, A.J., Watt, A.J., Sjöström, P.J., and van Meyel, D.J. (2014). Neuronal morphometry directly from bitmap images. Nature Methods 11, 982–984.

37. Finak, G., McDavid, A., Yajima, M., Deng, J., Gersuk, V., Shalek, A.K., Slichter, C.K., Miller, H.W., McElrath, M.J., Prlic, M., et al. (2015). MAST: a flexible statistical framework for assessing transcriptional changes and characterizing heterogeneity in single-cell RNA sequencing data. Genome Biology 16, 278.

38. Finlay, B., and Darlington, R. (1995). Linked regularities in the development and evolution of mammalian brains. Science 268, 1578–1584.

39. Fleming, S.J., Chaffin, M.D., Arduini, A., Akkad, A.-D., Banks, E., Marioni, J.C., Philippakis, A.A., Ellinor, P.T., and Babadi, M. (2022). Unsupervised removal of systematic background noise from droplet-based single-cell experiments using CellBender. bioRxiv, 791699.

40. Fontenot, M.R., Berto, S., Liu, Y., Werthmann, G., Douglas, C., Usui, N., Gleason, K., Tamminga, C.A., Takahashi, J.S., and Konopka, G. (2017). Novel transcriptional networks regulated by CLOCK in human neurons. Genes & Development 31, 2121–2135.

41. Fu, L., Pelicano, H., Liu, J., Huang, P., and Lee, C.C. (2002). The Circadian Gene Period2 Plays an Important Role in Tumor Suppression and DNA Damage Response In Vivo. Cell 111, 41–50.

42. Giza, J.I., Kim, J., Meyer, H.C., Anastasia, A., Dincheva, I., Zheng, C.I., Lopez, K., Bains, H., Yang, J., Bracken, C., et al. (2018). The BDNF Val66Met Prodomain Disassembles Dendritic Spines Altering Fear Extinction Circuitry and Behavior. Neuron 99, 163–178.e166.

43. Glerup, S., Bolcho, U., Mølgaard, S., Bøggild, S., Vaegter, C.B., Smith, A.H., Nieto-Gonzalez, J.L., Ovesen, P.L., Pedersen, L.F., Fjorback, A.N., et al. (2016). SorCS2 is required for BDNF-dependent plasticity in the hippocampus. Molecular Psychiatry 21, 1740–1751.

44. Grüneberg, H. (1938). An analysis of the &#x201c;pleiotropic&#x201d; effects of a new lethal mutation in the rat (*Mus norvegicus*). Proceedings of the Royal Society of London Series B - Biological Sciences 125, 123–144.

45. Hamilton, D.A., and Brigman, J.L. (2015). Behavioral flexibility in rats and mice: contributions of distinct frontocortical regions. Genes, Brain and Behavior 14, 4–21.

46. Hao, Y., Hao, S., Andersen-Nissen, E., Mauck, W.M., Zheng, S., Butler, A., Lee, M.J., Wilk, A.J., Darby, C., Zager, M., et al. (2021). Integrated analysis of multimodal single-cell data. Cell 184, 3573–3587.e3529.

47. Harris, K., Jensen, F., and Tsao, B. (1992). Three-dimensional structure of dendritic spines and synapses in rat hippocampus (CA1) at postnatal day 15 and adult ages: implications for the maturation of synaptic physiology and long-term potentiation. Journal of Neuroscience 12, 2685–2705.

48. Hayashi, Y., and Majewska, A.K. (2005). Dendritic Spine Geometry: Functional Implication and Regulation. Neuron 46, 529–532.

49. Heisler, J.M., Morales, J., Donegan, J.J., Jett, J.D., Redus, L., and O’Connor, J.C. (2015). The Attentional Set Shifting Task: A Measure of Cognitive Flexibility in Mice. JoVE, e51944.

50. Heyser, C.J. (2003). Assessment of developmental milestones in rodents. Current Protocols in Neuroscience 25, 8–18.

51. Ho, S.-M., Hartley, B.J., Tcw, J., Beaumont, M., Stafford, K., Slesinger, P.A., and Brennand, K.J. (2016). Rapid Ngn2-induction of excitatory neurons from hiPSC-derived neural progenitor cells. Methods 101, 113–124.

52. Hrvatin, S., Hochbaum, D.R., Nagy, M.A., Cicconet, M., Robertson, K., Cheadle, L., Zilionis, R., Ratner, A., Borges-Monroy, R., Klein, A.M., et al. (2018). Single-cell analysis of experience-dependent transcriptomic states in the mouse visual cortex. Nature Neuroscience 21, 120–129.

53. Kent, W.J., Sugnet, C.W., Furey, T.S., Roskin, K.M., Pringle, T.H., Zahler, A.M., Haussler, and David (2002). The human genome browser at UCSC. Genome Research 12, 996–1006.

54. Kang, H.J., Kawasawa, Y.I., Cheng, F., Zhu, Y., Xu, X., Li, M., Sousa, A.M.M., Pletikos, M., Meyer, K.A., Sedmak, G., et al. (2011). Spatio-temporal transcriptome of the human brain. Nature 478, 483–489.

55. Khrameeva, E., Kurochkin, I., Han, D., Guijarro, P., Kanton, S., Santel, M., Qian, Z., Rong, S., Mazin, P., Bulat, M., et al. (2019). Single-cell-resolution transcriptome map of human, chimpanzee, bonobo, and macaque brains. bioRxiv, 764936.

56. King, M., and Wilson, A. (1975). Evolution at two levels in humans and chimpanzees. Science 188, 107–116.

57. Kim, J.-Y., Grunke, S.D., Levites, Y., Golde, T.E., and Jankowsky, J.L. (2014). Intracerebroventricular viral injection of the neonatal mouse brain for persistent and widespread neuronal transduction. Journal of Visualized Experiments, e51863.

58. Kobayashi, Y., Ye, Z., and Hensch, Takao K. (2015). Clock Genes Control Cortical Critical Period Timing. Neuron 86, 264–275.

59. Kondratova, A.A., Dubrovsky, Y.V., Antoch, M.P., and Kondratov, R.V. (2010). Circadian clock proteins control adaptation to novel environment and memory formation. Aging 2, 285–297.

60. Konopka, G., Bomar, J.M., Winden, K., Coppola, G., Jonsson, Z.O., Gao, F., Peng, S., Preuss, T.M., Wohlschlegel, J.A., and Geschwind, D.H. (2009). Human-specific transcriptional regulation of CNS development genes by FOXP2. Nature 462, 213–217.

61. Konopka, G., Friedrich, T., Davis-Turak, J., Winden, K., Oldham, M.C., Gao, F., Chen, L., Wang, G.-Z., Luo, R., and Preuss, T.M. (2012). Human-specific transcriptional networks in the brain. Neuron 75, 601–617.

62. Konopka, G., and Geschwind, D.H. (2010). Human brain evolution: harnessing the genomics (r)evolution to link genes, cognition, and Behavior. Neuron 68, 231–244.

63. Kumar, V., Kim, K., Joseph, C., Kourrich, S., Yoo, S.H., Huang, H.C., Vitaterna, M.H., Pardo-Manuel de Villena, F., Churchill, G., Bonci, A., and Takahashi, J.S. (2013). C57BL/6N mutation in cytoplasmic FMRP interacting protein 2 regulates cocaine response. Science 342,1508–1512.

64. Labun, K., Montague, T.G., Krause, M., Torres Cleuren, Y.N., Tjeldnes, H., and Valen, E. (2019). CHOPCHOP v3: expanding the CRISPR web toolbox beyond genome editing. Nucleic Acids Research 47, W171–W174.

65. Lamont, E.W., Legault-Coutu, D., Cermakian, N., and Boivin, D.B. (2007). The role of circadian clock genes in mental disorders. Dialogues Clin Neurosci 9, 333–342.

66. Lane, N., and Martin, W. (2010). The energetics of genome complexity. Nature 467, 929–934.

67. Leggio, M.G., Mandolesi, L., Federico, F., Spirito, F., Ricci, B., Gelfo, F., and Petrosini, L. (2005). Environmental enrichment promotes improved spatial abilities and enhanced dendritic growth in the rat. Behavioural Brain Research 163, 78–90.

68. Leuner, B., and Gould, E. (2010). Dendritic Growth in Medial Prefrontal Cortex and Cognitive Flexibility Are Enhanced during the Postpartum Period. The Journal of Neuroscience 30, 13499–13503.

69. Levet, F., Tønnesen, J., Nägerl, U.V., and Sibarita, J.-B. (2020). SpineJ: A software tool for quantitative analysis of nanoscale spine morphology. Methods 174, 49–55.

70. Li, H., Handsaker, B., Wysoker, A., Fennell, T., Ruan, J., Homer, N., Marth, G., Abecasis, G., Durbin, R., and Subgroup, G.P.D.P. (2009). The Sequence Alignment/Map format and SAMtools. Bioinformatics 25, 2078–2079.

71. Li, J., Shalev-Benami, M., Sando, R., Jiang, X., Kibrom, A., Wang, J., Leon, K., Katanski, C., Nazarko, O., Lu, Y.C., et al. (2018). Structural Basis for Teneurin Function in Circuit-Wiring: A Toxin Motif at the Synapse. Cell 173, 735–748.e715.

72. Li, P., Fu, X., Smith, N.A., Ziobro, J., Curiel, J., Tenga, M.J., Martin, B., Freedman, S., Cea-Del Rio, C.A., Oboti, L., et al. (2017). Loss of CLOCK Results in Dysfunction of Brain Circuits Underlying Focal Epilepsy. Neuron 96, 387–401.e386.

73. Liao, Y., Smyth, G.K., and Shi, W. (2013). FeatureCounts: an efficient general purpose program for assigning sequence reads to genomic features. Bioinformatics 30, 923–930.

74. Lipton, J.O., Yuan, E.D., Boyle, L.M., Ebrahimi-Fakhari, D., Kwiatkowski, E., Nathan, A., Güttler, T., Davis, F., Asara, J.M., and Sahin, M. (2015). The circadian protein BMAL1 regulates translation in response to S6K1-mediated phosphorylation. Cell 161, 1138–1151.

75. Longair, M.H., Baker, D.A., and Armstrong, J.D. (2011). Simple Neurite Tracer: open source software for reconstruction, visualization and analysis of neuronal processes. Bioinformatics 27, 2453–2454.

76. Loo, L., Simon, J.M., Xing, L., McCoy, E.S., Niehaus, J.K., Guo, J., Anton, E.S., and Zylka, M.J. (2019). Single-cell transcriptomic analysis of mouse neocortical development. Nature Communications 10, 134.

77. Matharu, N., Rattanasopha, S., Tamura, S., Maliskova, L., Wang, Y., Bernard, A., Hardin, A., Eckalbar, W.L., Vaisse, C., and Ahituv, N. (2019). CRISPR-mediated activation of a promoter or enhancer rescues obesity caused by haploinsufficiency. Science 363, eaau0629.

78. Matsuo, T., Yamaguchi, S., Mitsui, S., Emi, A., Shimoda, F., and Okamura, H. (2003). Control Mechanism of the Circadian Clock for Timing of Cell Division in Vivo. Science 302, 255–259.

79. McAllister, A.K. (2000). Cellular and Molecular Mechanisms of Dendrite Growth. Cerebral Cortex 10, 963–973.

80. McGinnis, C.S., Murrow, L.M., and Gartner, Z.J. (2019). DoubletFinder: doublet detection in single-cell RNA sequencing data using artificial nearest neighbors. Cell Systems 8, 329–337.

81. McIlwain, K.L., Merriweather, M.Y., Yuva-Paylor, L.A., and Paylor, R. (2001). The use of behavioral test batteries: Effects of training history. Physiology & Behavior 73, 705–717.

82. Micheva, K.D., and Beaulieu, C. (1996). Quantitative aspects of synaptogenesis in the rat barrel field cortex with special reference to GABA circuitry. Journal of Comparative Neurology 373, 340–354.

83. Miller, B.H., McDearmon, E.L., Panda, S., Hayes, K.R., Zhang, J., Andrews, J.L., Antoch, M.P., Walker, J.R., Esser, K.A., Hogenesch, J.B., et al. (2007). Circadian and CLOCK-controlled regulation of the mouse transcriptome and cell proliferation. Proceedings of the National Academy of Sciences 104, 3342–3347.

84. Missler, M., Zhang, W., Rohlmann, A., Kattenstroth, G., Hammer, R.E., Gottmann, K., and Südhof, T.C. (2003). α-Neurexins couple Ca2+ channels to synaptic vesicle exocytosis. Nature 423, 939–948.

85. Moore, J.E., Purcaro, M.J., Pratt, H.E., Epstein, C.B., Shoresh, N., Adrian, J., Kawli, T., Davis, C.A., Dobin, A., Kaul, R., et al., ENCODE Project Consortium. (2020). Expanded encyclopaedias of DNA elements in the human and mouse genomes. Nature 583, 699–710.

86. Mure, L.S., Le, H.D., Benegiamo, G., Chang, M.W., Rios, L., Jillani, N., Ngotho, M., Kariuki, T., Dkhissi-Benyahya, O., Cooper, H.M., et al. (2018). Diurnal transcriptome atlas of a primate across major neural and peripheral tissues. Science 359, eaao0318.

87. Musiek, E.S. (2015). Circadian clock disruption in neurodegenerative diseases: cause and effect? Frontiers in Pharmacology 6.

88. Musiek, E.S., Lim, M.M., Yang, G., Bauer, A.Q., Qi, L., Lee, Y., Roh, J.H., Ortiz-Gonzalez, X., Dearborn, J.T., Culver, J.P., et al. (2013). Circadian clock proteins regulate neuronal redox homeostasis and neurodegeneration. The Journal of Clinical Investigation 123, 5389–5400.

89. Nagoshi, E., Saini, C., Bauer, C., Laroche, T., Naef, F., and Schibler, U. (2004). Circadian Gene Expression in Individual Fibroblasts: Cell-Autonomous and Self-Sustained Oscillators Pass Time to Daughter Cells. Cell 119, 693–705.

90. Necsulea, A., and Kaessmann, H. (2014). Evolutionary dynamics of coding and non-coding transcriptomes. Nature Reviews Genetics 15, 734–748.

91. O’Brien, C., Unruh, L., Zimmerman, C., Bradshaw, W.E., Holzapfel, C.M., and Cresko, W.A. (2013). Geography of the circadian gene clock and photoperiodic response in western North American populations of the three-spined stickleback Gasterosteus aculeatus. Journal of Fish Biology 82, 827–839.

92. Odawara, A., Saitoh, Y., Alhebshi, A.H., Gotoh, M., and Suzuki, I. (2014). Long-term electrophysiological activity and pharmacological response of a human induced pluripotent stem cell-derived neuron and astrocyte co-culture. Biochemical and Biophysical Research Communications 443, 1176–1181.

93. Ovcharenko, I., Loots, G.G., Giardine, B.M., Hou, M., Ma, J., Hardison, R.C., Stubbs, L., and Miller, W. (2005). Mulan: Multiple-sequence local alignment and visualization for studying function and evolution. Genome Research 15, 184–194.

94. Panda, S., Antoch, M.P., Miller, B.H., Su, A.I., Schook, A.B., Straume, M., Schultz, P.G., Kay, S.A., Takahashi, J.S., and Hogenesch, J.B. (2002). Coordinated Transcription of Key Pathways in the Mouse by the Circadian Clock. Cell 109, 307–320.

95. Pattabiraman, K., Muchnik, S.K., and Sestan, N. (2020). The evolution of the human brain and disease susceptibility. Current Opinion in Genetics & Development 65, 91–97.

96. Peters, A., and Kaiserman-Abramof, I.R. (1970). The small pyramidal neuron of the rat cerebral cortex. The perikaryon, dendrites and spines. American Journal of Anatomy 127, 321–355.

97. Preitner, N., Damiola, F., Luis Lopez, M., Zakany, J., Duboule, D., Albrecht, U., and Schibler, U. (2002). The Orphan Nuclear Receptor REV-ERBα Controls Circadian Transcription within the Positive Limb of the Mammalian Circadian Oscillator. Cell 110, 251–260.

98. Ramirez, D.M.O., and Kavalali, E.T. (2011). Differential regulation of spontaneous and evoked neurotransmitter release at central synapses. Current Opinion in Neurobiology 21, 275–282.

99. Ran, F.A., Hsu, P.D., Wright, J., Agarwala, V., Scott, D.A., and Zhang, F. (2013). Genome engineering using the CRISPR-Cas9 system. Nature Protocols 8, 2281–2308.

100. Rath, M.F., Rohde, K., Fahrenkrug, J., and Møller, M. (2013). Circadian clock components in the rat neocortex: daily dynamics, localization and regulation. Brain Structure and Function 218, 551–562.

101. Rath, M.F., Rovsing, L., and Møller, M. (2014). Circadian oscillators in the mouse brain: molecular clock components in the neocortex and cerebellar cortex. Cell and Tissue Research 357, 743–755.

102. Roybal, K., Theobold, D., Graham, A., DiNieri, J.A., Russo, S.J., Krishnan, V., Chakravarty, S., Peevey, J., Oehrlein, N., and Birnbaum, S. (2007). Mania-like behavior induced by disruption of CLOCK. Proceedings of the National Academy of Sciences 104, 6406–6411.

103. Saino, N., Bazzi, G., Gatti, E., Caprioli, M., Cecere, J.G., Possenti, C.D., Galimberti, A., Orioli, V., Bani, L., Rubolini, D., et al. (2015). Polymorphism at the Clock gene predicts phenology of long-distance migration in birds. Molecular Ecology 24, 1758–1773.

104. Saleem, Q., Anand, A., Jain, S., and Brahmachari, S.K. (2001). The polyglutamine motif is highly conserved at the Clock locus in various organisms and is not polymorphic in humans. Human Genetics 109, 136–142.

105. Sambrook, J., and Russell, D.W. (2006). Purification of nucleic acids by extraction with phenol:chloroform. Cold Spring Harbor Protocols 2006, pdb.prot4455.

106. Sayers, E.W., Bolton, E.E., Brister, J.R., Canese, K., Chan, J., Comeau, Donald C., Connor, R., Funk, K., Kelly, C., Kim, S., et al. (2021). Database resources of the national center for biotechnology information. Nucleic Acids Research 50, D20–D26.

107. Schindelin, J., Arganda-Carreras, I., Frise, E., Kaynig, V., Longair, M., Pietzsch, T., Preibisch, S., Rueden, C., Saalfeld, S., Schmid, B., et al. (2012). Fiji: an open-source platform for biological-image analysis. Nature Methods 9, 676–682.

108. Schmidt, E.R.E., Kupferman, J.V., Stackmann, M., and Polleux, F. (2019). The human-specific paralogs SRGAP2B and SRGAP2C differentially modulate SRGAP2A-dependent synaptic development. Scientific Reports 9, 18692.

109. Schmidt, E.R.E., and Polleux, F. (2022). Genetic Mechanisms Underlying the Evolution of Connectivity in the Human Cortex. Frontiers in Neural Circuits 15.

110. Schmidt, E.R.E., Zhao, H.T., Park, J.M., Dipoppa, M., Monsalve-Mercado, M.M., Dahan, J.B., Rodgers, C.C., Lejeune, A., Hillman, E.M.C., Miller, K.D., et al. (2021). A human-specific modifier of cortical connectivity and circuit function. Nature.

111. Schuch, J.B., Genro, J.P., Bastos, C.R., Ghisleni, G., and Tovo-Rodrigues, L. (2018). The role of CLOCK gene in psychiatric disorders: Evidence from human and animal research. American Journal of Medical Genetics Part B: Neuropsychiatric Genetics 177, 181–198.

112. Semple, B.D., Blomgren, K., Gimlin, K., Ferriero, D.M., and Noble-Haeusslein, L.J. (2013). Brain development in rodents and humans: Identifying benchmarks of maturation and vulnerability to injury across species. Progress in Neurobiology 106-107, 1–16.

113. Sherwood, C.C., Miller, S.B., Karl, M., Stimpson, C.D., Phillips, K.A., Jacobs, B., Hof, P.R., Raghanti, M.A., and Smaers, J.B. (2020). Invariant Synapse Density and Neuronal Connectivity Scaling in Primate Neocortical Evolution. Cerebral Cortex.

114. Shibata, M., Pattabiraman, K., Muchnik, S.K., Kaur, N., Morozov, Y.M., Cheng, X., Waxman, S.G., and Sestan, N. (2021). Hominini-specific regulation of CBLN2 increases prefrontal spinogenesis. Nature 598, 489–494.

115. Sholl, D. (1953). Dendritic organization in the neurons of the visual and motor cortices of the cat. Journal of anatomy 87, 387–406.

116. Siepka, S.M., and Takahashi, J.S. (2005). Methods to record circadian rhythm wheel running activity in mice. In Methods in Enzymology, M.W. Young, ed. (Academic Press), pp. 230–239.

117. Smith, T.S., Heger, A., and Sudbery, I. (2017). UMI-tools: modelling sequencing errors in Unique Molecular Identifiers to improve quantification accuracy. Genome Research, 491–499.

118. Striedter, G.F. (2005). Principles of brain evolution (Sunderland, MA: Sinauer).

119. Südhof, T.C., and Rothman, J.E. (2009). Membrane Fusion: Grappling with SNARE and SM Proteins. Science 323, 474–477.

120. Takahashi, J.S. (2017). Transcriptional architecture of the mammalian circadian clock. Nature Reviews Genetics 18, 164–179.

121. Turrigiano, G. (2012). Homeostatic synaptic plasticity: local and global mechanisms for stabilizing neuronal function. Cold Spring Harbor Perspectives in Biology 4, a005736.

122. Turek, F.W., Joshu, C., Kohsaka, A., Lin, E., Ivanova, G., McDearmon, E., Laposky, A., Losee-Olson, S., Easton, A., Jensen, D.R., et al. (2005). Obesity and Metabolic Syndrome in Circadian *Clock* Mutant Mice. Science 308, 1043–1045.

123. Ueno, H., Takahashi, Y., Suemitsu, S., Murakami, S., Kitamura, N., Wani, K., Matsumoto, Y., Okamoto, M., and Ishihara, T. (2020). Effects of repetitive gentle handling of male C57BL/6NCrl mice on comparative behavioural test results. Scientific Reports 10, 3509.

124. Untergasser, A., Cutcutache, I., Koressaar, T., Ye, J., Faircloth, B.C., Remm, M., and Rozen, S.G. (2012). Primer3-new capabilities and interfaces. Nucleic Acids Research 40, e115.

125. Usui, N., Araujo, D.J., Kulkarni, A., Co, M., Ellegood, J., Harper, M., Toriumi, K., Lerch, J.P., and Konopka, G. (2017a). Foxp1 regulation of neonatal vocalizations via cortical development. Genes & Development 31, 2039–2055.

126. Usui, N., Co, M., Harper, M., Rieger, M.A., Dougherty, J.D., and Konopka, G. (2017b). Sumoylation of FOXP2 regulates motor function and vocal communication through Purkinje cell development. Biological Psychiatry 81, 220–230.

127. Van der Zee, E.A., Havekes, R., Barf, R.P., Hut, R.A., Nijholt, I.M., Jacobs, E.H., and Gerkema, M.P. (2008). Circadian Time-Place Learning in Mice Depends on Cry Genes. Current Biology 18, 844–848.

128. Velmeshev, D., Schirmer, L., Jung, D., Haeussler, M., Perez, Y., Mayer, S., Bhaduri, A., Goyal, N., Rowitch, D.H., and Kriegstein, A.R. (2019). Single-cell genomics identifies cell type-specific molecular changes in autism. Science 364, 685–689.

129. Wagner, G.P., and Zhang, J. (2011). The pleiotropic structure of the genotype–phenotype map: the evolvability of complex organisms. Nature Reviews Genetics 12, 204–213.

130. Wilsbacher, L.D., Sangoram, A.M., Antoch, M.P., and Takahashi, J.S. (2000). The Mouse Clock Locus: Sequence and Comparative Analysis of 204 Kb from Mouse Chromosome 5. Genome Research 10, 1928–1940.

131. Winslow, J.T., and Camacho, F. (1995). Cholinergic modulation of a decrement in social investigation following repeated contacts between mice. Psychopharmacology 121, 164–172.

132. Xu, C., Li, Q., Efimova, O., He, L., Tatsumoto, S., Stepanova, V., Oishi, T., Udono, T., Yamaguchi, K., Shigenobu, S., et al. (2018). Human-specific features of special gene expression and regulation in eight brain regions. Genome Research 28, 1097–1110.

133. Xu, P., Berto, S., Kulkarni, A., Jeong, B., Joseph, C., Cox, K.H., Greenberg, M.E., Kim, T.-K., Konopka, G., and Takahashi, J.S. (2021). NPAS4 regulates the transcriptional response of the suprachiasmatic nucleus to light and circadian behavior. Neuron 109, 3268–3282.

134. Yang, M., and Crawley, J.N. (2001). Simple behavioral assessment of mouse olfaction. In Current Protocols in Neuroscience (John Wiley & Sons, Inc.).

135. Yao, Z., van Velthoven, C.T.J., Nguyen, T.N., Goldy, J., Sedeno-Cortes, A.E., Baftizadeh, F., Bertagnolli, D., Casper, T., Chiang, M., Crichton, K., et al. (2021). A taxonomy of transcriptomic cell types across the isocortex and hippocampal formation. Cell 184, 3222–3241.e3226.

136. Zhang, Y., Pak, C., Han, Y., Ahlenius, H., Zhang, Z., Chanda, S., Marro, S., Patzke, C., Acuna, C., Covy, J., et al. (2013). Rapid Single-Step Induction of Functional Neurons from Human Pluripotent Stem Cells. Neuron 78, 785–798.

137. Zheng, G.X.Y., Terry, J.M., Belgrader, P., Ryvkin, P., Bent, Z.W., Wilson, R., Ziraldo, S.B., Wheeler, T.D., McDermott, G.P., Zhu, J., et al. (2017). Massively parallel digital transcriptional profiling of single cells. Nature Communications 8, 14049.

138. Zhu, Y., Sousa, A.M.M., Gao, T., Skarica, M., Li, M., Santpere, G., Esteller-Cucala, P., Juan, D., Ferrández-Peral, L., Gulden, F.O., et al. (2018). Spatiotemporal transcriptomic divergence across human and macaque brain development. Science 362, eaat8077.

